# Reference Regulatory Element-Guided Gene Expression Analysis for Mechanistic Inference of Gene Regulatory Networks

**DOI:** 10.64898/2026.07.10.737848

**Authors:** Lixin Ren, Ishita Debnath, Zhana Duren

**Affiliations:** Center for Computational Biology and Bioinformatics, Department of Medical and Molecular Genetics, Indiana University School of Medicine, Indianapolis, IN 46202, USA

## Abstract

Regulatory genomics faces a depth–breadth gap: deep multi-omics provides regulatory detail but is difficult to scale, whereas broad expression datasets often lack the regulatory structure needed for mechanistic Gene Regulatory Network (GRN) analysis. We developed Regulatory Elements Guided Analysis (REGA), an interpretable framework that uses reference Regulatory Element (RE) catalogs to infer transcription factor (TF)–RE–gene programs from gene expression data. Across ChIP-seq, knockdown, Hi-C, *cis*- and *trans*-eQTL benchmarks, REGA prioritized functional REs, improved RE–gene and TF–gene inference over existing baselines, including methods using more data, and recovered coherent regulatory modules. In PsychENCODE snRNA-seq, REGA identified disease-associated modules and TF activities, linked regulatory dysregulation to genetic risk, and detected cross-cell-type neuronal–glial programs. In spatial transcriptomics, REGA linked cell-intrinsic regulatory programs with intercellular ligand–receptor communication; in Perturb-seq, it mapped perturbation responses to trait-associated regulatory architectures. REGA enables scalable, interpretable GRN analysis across expression datasets.

## Introduction

Modern regulatory genomics has generated two complementary but often disconnected resources: deep multi-omics datasets that resolve regulatory mechanisms in detail, and broad transcriptomic datasets that capture many samples, cell states, perturbations, and phenotypes. The former provides regulatory resolution but remains difficult to scale, whereas the latter provides biological breadth but often lacks the regulatory structure needed for mechanistic interpretation. Gene regulatory networks (GRNs) serve as the fundamental blueprints of cellular identity and function, orchestrating how genetic information translates into complex physiological traits and disease states^1–3^. Consequently, the reverse engineering of GRNs from high-dimensional biological data has been a central pursuit of computational biology for decades^4–7^. Early efforts relied predominantly on bulk gene expression data to infer regulatory relationships^8–10^. As technology advanced, methods evolved to incorporate single-cell RNA-sequencing (scRNA-seq^11,12^) and a rich tapestry of multi-omics modalities^13–19^, including chromatin accessibility (ATAC-seq^20^), physical interactions (Hi-C^21,22^), histone modifications^23^, and DNA methylation^24^. More recently, single-cell multiome assays, which simultaneously capture transcriptomic and epigenomic profiles from the exact same cell, have emerged as a prominent approach for dissecting these intricate regulatory layers^25–27^.

Despite these technological leaps, the accurate inference of GRNs remains a severe bottleneck in the field^8,9^. Traditional expression-only methods are notoriously limited, typically performing a mere <20% better than random guessing when compared against experimental ground truths^10,28^. To exploit epigenomic insights, frameworks like SCENIC+^29^ incorporate REs into the inference pipeline; however, they primarily use REs data post-hoc to connect and map TF-gene relationships previously detected by expression-only methods like GENIE3^10^, rather than utilizing RE information to actively guide the model toward learning better underlying TF-gene interactions. On the other end of the spectrum, advanced foundation models such as LINGER utilize external datasets as priors and fine-tune them using paired single-cell multiome data to push this accuracy to approximately 125% better than random^28^. These strategies are powerful when the required data are available, but their applicability is limited because majority of transcriptomic studies lack paired epigenomic profiles^3^. These multi-omics modalities remain impractical to acquire at scale due to severe financial constraints in large patient cohorts, technical barriers in specialized assays like perturb-seq^30,31^, and commercialization restrictions or low resolutions in emerging spatial transcriptomics technologies^32,33^. Moreover, simply combining multiple molecular modalities does not by itself yield mechanistic insight; regulatory mechanisms must be organized into interpretable relationships among TFs, REs, target genes, cellular contexts, phenotypes, and genetic variation. Thus, a major unmet need is a framework that can transfer regulatory knowledge from deep reference resources into broad expression datasets.

A growing opportunity comes from public epigenomic resources. Large-scale efforts such as ENCODE project^34^ and related atlases have generated high-quality reference RE catalogs across human and various model organisms^35–49^ (Fig. 1a, b). These reference catalogs provide reusable regulatory knowledge: they identify genomic regions likely to be functional in a given tissue or cell type, can be linked to nearby genes to define candidate *cis*-regulatory priors, and can be combined with TF motif information to infer potential TF–RE relationships. Together, reference REs and TF motifs define a prior regulatory scaffold connecting TFs, REs, and genes. This provides a way to guide expression-based GRN inference with regulatory structure, rather than relying on co-expression alone.

**Figure 1.**
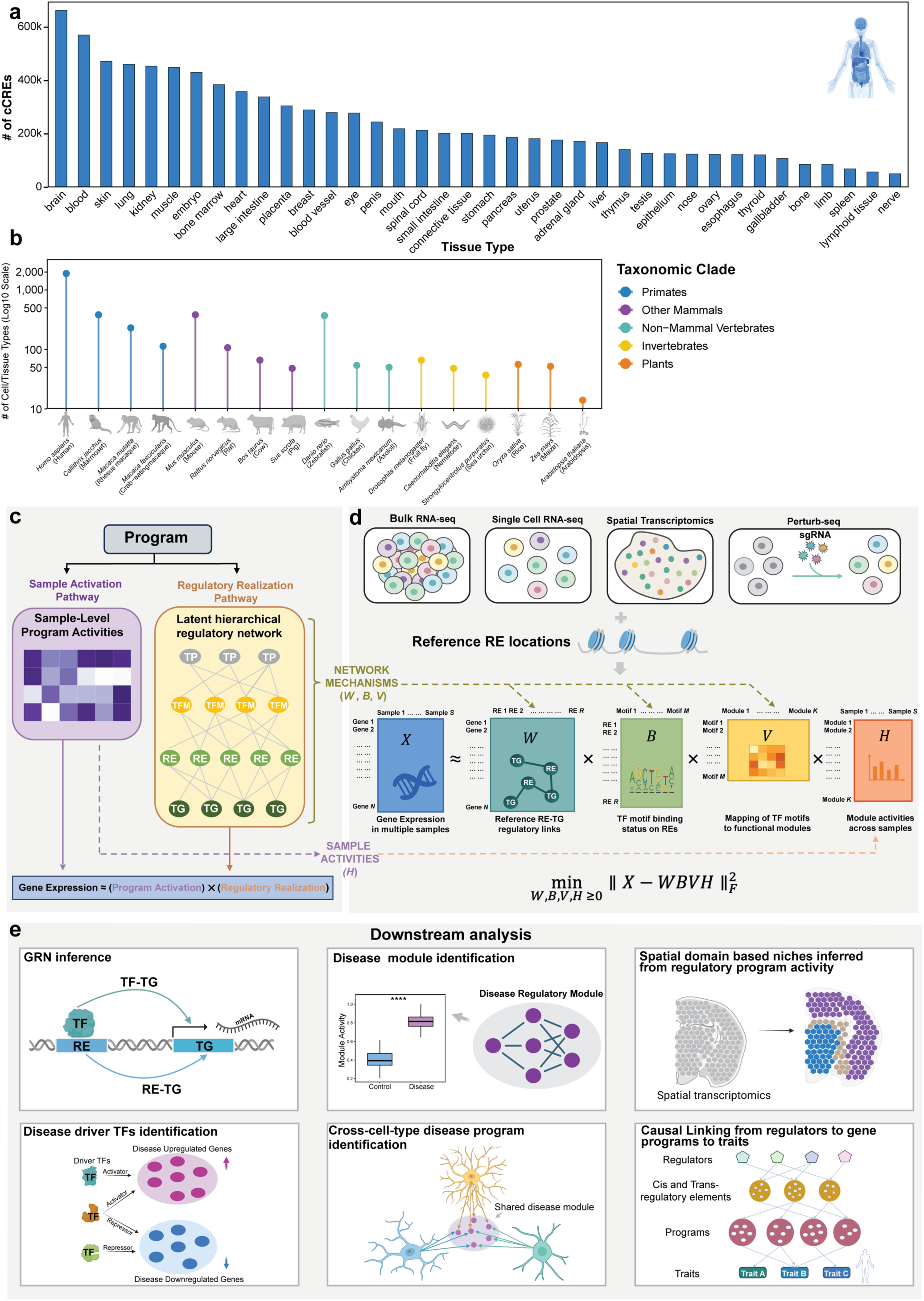
Schematic overview of the REGA framework. **(a)** Bar chart showing the number of ENCODE4 candidate cis-regulatory elements (cCREs) across human tissues, ordered by abundance. **(b)** Lollipop plot displaying the number of cell or tissue types (presented on a log10 scale) for which chromatin accessibility data is available, based on recent literature surveys. Diverse species are grouped and colored by taxonomic clade: Primates (blue), Other Mammals (purple), Non-Mammal Vertebrates (teal), Invertebrates (yellow), and Plants (orange). **(c)** Conceptual framework of REGA, illustrating the decomposition of transcriptional programs (TP) into two complementary pathways: a sample-specific activation pathway and a regulatory realization pathway, the latter representing a latent hierarchical regulatory network connecting target genes (TG), regulatory elements (RE), and transcription factor motifs (TFM). **(d)** Computational workflow of REGA. Taking an expression matrix (*X*) spanning diverse transcriptomic modalities — including bulk RNA-seq, single-cell RNA-seq, spatial transcriptomics, and Perturb-seq — and a reference RE set, REGA factorizes X into biologically interpretable matrices representing RE–TG links (*W*), TF motif binding on REs (*B*), motif–module associations (*V*), and sample-specific module activities (*H*). **(e)** Downstream applications enabled by REGA, including gene regulatory network inference, disease module identification, cross-cell-type disease program discovery, spatial regulatory domain analysis, driver TF prioritization, and regulator-to-trait linking.

To leverage this opportunity, we developed Regulatory Elements Guided Analysis (REGA), an interpretable hierarchical framework that uses reference RE catalogs and gene expression data to infer active TF–RE–target gene programs. REGA resolves the stark trade-off between data accessibility and performance by capturing a substantial portion of multi-omics accuracy while maintaining the low data burden of expression-only methods. Our approach is founded on two core biological and computational insights. First, co-regulated genes do not merely exhibit co-expression patterns; their *cis*-REs are also highly enriched for sequence motifs corresponding to active TFs. Second, while quantitative chromatin accessibility measured at the single-cell level promises high-resolution detail, it is ultimately so sparse and noisy that it provides marginal additional information for network inference compared to a consolidated, high-confidence peak list at the cell-type level. Knowing *where* the genome is accessible in a given cell type provides a robust, physically hardcoded blueprint^50,51^. By relying on this static “dictionary” of potential interactions and leaving it to the dynamic fluctuations of gene expression to reveal which connections are active, REGA bypasses technical noise and eliminates the need for expensive, cell/individual-level matched multi-omics data.

This manuscript presents and validates REGA, yielding several key contributions to the field. First, REGA leverages publicly available, non-sample-specific reference RE lists (Fig. 1a, b) alongside ubiquitous gene expression data as its sole inputs, eliminating the requirement for cell-level epigenome data and drastically lowering the data barrier for high-accuracy GRN inference. Second, our framework bridges the gap between expression-only methods and costly multi-omics approaches by simultaneously modeling RE–gene and TF–gene networks; because both TFs and REs converge on shared target genes, REGA effectively maps the complete TF–RE–TG regulatory axis to capture a physical layer of biology that expression-only methods cannot access, significantly improving the inference of direct TF–gene relationships (Fig. 1c). Third, this approach boasts broad and versatile applicability, demonstrating robust performance across four distinct types of transcriptomic data: bulk gene expression, single-cell expression, spatial transcriptomics, and perturb-seq data (Fig. 1d). Fourth, REGA estimates sample- or cell-level regulatory program activities, enabling modules to be associated with disease status, clinical phenotypes, genetic risk, and perturbation effects (Fig. 1e). Fifth, REGA provides a general framework for discovering cross-cell-type regulatory programs. Instead of analyzing each cell type independently and interpreting shared signals post hoc, REGA directly identifies regulatory architectures shared across cellular contexts and connects them to disease status, genetic risk, spatial neighborhood organization, and perturbation response. Together, REGA lowers the data and workflow barrier for mechanistic regulatory genomics by converting expression data plus reference REs into interpretable regulatory programs.

## Results

### Overview of REGA: reference regulatory element-guided inference of regulatory programs

We developed **Regulatory Elements Guided Analysis (REGA)** to infer interpretable gene regulatory programs from transcriptomic data using reference regulatory element (RE) catalogs. REGA takes as input a gene expression matrix and a context-relevant reference RE catalog, together with TF motif annotations, and uses these sources to define an initial regulatory scaffold connecting transcription factors (TFs), REs, and genes (Fig. 1a–c). Rather than relying on co-expression alone, REGA uses this scaffold to guide the discovery of regulatory programs supported by both coordinated gene expression and plausible upstream regulatory structure. The key idea is that truly co-regulated genes should show two forms of evidence: coordinated expression and shared upstream regulatory support. Co-expression alone is ambiguous, because correlated genes may be directly regulated or may simply share a common regulator. By incorporating RE and motif information, REGA helps distinguish gene modules that are merely correlated from those supported by a plausible regulatory mechanism.

REGA represents transcriptional regulation through a set of latent regulatory programs. Each program links three regulatory layers: downstream target genes, their associated REs, and upstream TFs. During training, REGA learns both the activity of each regulatory program across samples or cells and the regulatory realization of each program through the TF–RE–gene hierarchy (Fig. 1d). This design encourages REGA to learn programs that not only explain observed gene expression variation, but also remain consistent with biologically plausible regulatory paths (See Methods for detail).

The fundamental outputs of REGA are interpretable regulatory representations at multiple biological levels. First, REGA produces refined RE–gene, and TF–gene regulatory networks by computing attention scores between learned node embeddings. Second, it decodes the membership of each transcriptional program, identifying the TFs, REs, and genes that define each regulatory module. Third, it estimates sample-specific activities for REs, TFs, and transcriptional programs, providing a high-resolution view of how regulatory mechanisms vary across a population or condition series. Fourth, it computes a regulatory importance score for each individual RE. Together, these outputs allow REGA to reconstruct regulatory architecture from transcriptomic data while remaining anchored to prior biological knowledge derived from public epigenomic resources and TF-binding sequence preferences (Fig. 1e).

Finally, we extended this framework into a cross-cell-type model to capture coordinated regulatory dynamics across an individual’s tissues (Fig. 1e, Supplementary Fig. 1). In this mode, sample activity is shared across cell types for each person, representing their unique systemic identity, while the regulatory realization remains cell-type specific. By reconstructing cell-type-specific expression through this shared identity, REGA identifies cross-cell-type regulatory programs that are more likely to be associated with complex individual phenotypes and GWAS-linked traits. We validate the utility and accuracy of these inferred networks and program activities in the following sections across diverse biological contexts.

### REGA improves GRN inference and identifies co-regulatory modules in bulk RNA-seq

To evaluate REGA, we applied it to GTEx whole-blood bulk RNA-seq data^52^ using a 10x Genomics PBMC multiome ATAC-seq peak set as the RE reference (see Data Availability). To assess whether REGA prioritizes functional REs, we collected ENCODE^53^ ChIP-seq data for active promoter (H3K4me3^54^) and enhancer (H3K27ac^55^) histone modifications across cell types. As shown in Fig. 2a, the top-ranked REs based on inferred regulatory importance scores showed consistent and significant enrichment for active histone marks across all examined cell types, with average fold changes of 2.15 (range: 1.62–2.57) for H3K27ac and 2.42 (range: 2.23–2.63) for H3K4me3. To further validate the functional importance of the prioritized REs from an evolutionary perspective^56^, we compared the sequence average conservation scores between REs with high (’Top’) and low (’Bottom’) inferred regulatory importance. We performed this comparison across varying thresholds, ranging from the top/bottom 10% to 50%. As shown in Fig. 2b, REs in the top tier exhibited significantly higher conservation scores compared to those in the bottom tier. Thus, REGA prioritizes functional and evolutionarily conserved REs without using histone-mark or conservation information as input.

**Figure 2.**
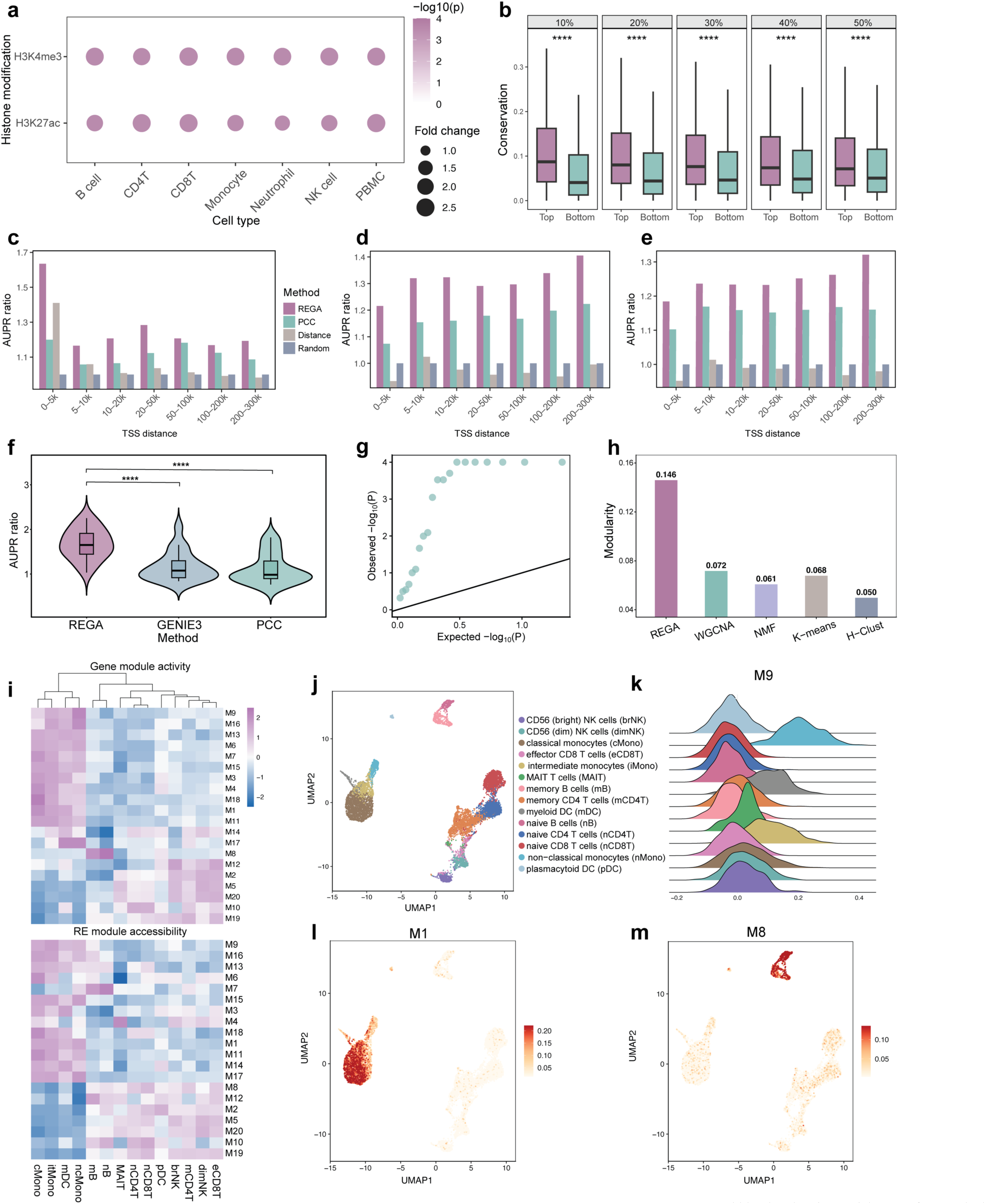
REGA improves gene regulatory network inference and identifies co-regulatory modules from bulk RNA-seq data. **(a)** Bubble plot showing enrichment of top-ranked REGA-inferred REs in active histone modification peaks (H3K4me3 and H3K27ac) across blood-related cell types. Bubble size reflects fold enrichment, and color indicates statistical significance (−log10(*P*), permutation test). **(b)** Boxplots comparing sequence conservation scores between REs with high versus low REGA-inferred regulatory importance across percentile thresholds (10–50%; one-sided t-test, **** *P* < 0.0001). **(c–e)** Bar charts showing area under the precision–recall curve (AUPR) ratios for predicting regulatory element–target gene (RE–TG) interactions in whole blood across seven genomic distance bins (0–5 kb to 200–300 kb), benchmarked against PCC, Distance, and Random baselines using three ground-truth datasets: **(c)** promoter-capture Hi-C, **(d)** GTEx whole-blood eQTLs, and **(e)** eQTLGen eQTLs. **(f)** Violin plots showing AUPR ratios for trans-regulatory potential prediction across 20 transcription factors (TFs), validated using ChIP-seq data (paired one-sided t-test, **** *P* < 0.0001). **(g)** Q–Q plot of permutation-based *P*-values from network enrichment analysis of REGA-derived modules within a trans-eQTL co-regulatory network. **(h)** Bar plot comparing Newman’s modularity (*Q*) scores across REGA, WGCNA, NMF, K-means, and hierarchical clustering (H-Clust). **(i)** Heatmaps showing pseudobulk RNA and ATAC module activities across PBMC cell types, ordered by hierarchical clustering. Heatmaps showing pseudobulk. RNA activity represents the average expression of module genes; ATAC activity reflects the average accessibility of module REs. **(j)** UMAP visualization of 14 annotated PBMC cell types. **(k)** Ridge plot showing the distribution of M9 module activity across cell types. **(l–m)** UMAP plots showing single-cell activity patterns of modules M1 and M8. Module activity was calculated using Seurat’s *AddModuleScore* based on log-normalized RNA expression.

To validate the *cis*-regulatory inferences of REGA, we utilized integrated promoter-capture Hi-C data covering 17 primary blood cell types^57^ and whole blood eQTL data from GTEx^52^ and eQTLGen^58^ as ground truth. We evaluated REGA’s ability to predict RE–gene interactions in bulk whole blood RNA-seq data, benchmarking it against three alternative approaches: a Pearson correlation coefficient (PCC) baseline derived from an external and independent single-cell multiome dataset, a genomic distance decay model, and a random baseline. We did not expect REGA to surpass PCC because it utilizes paired single-cell RNA and ATAC data, whereas REGA does not use ATAC data. To enable a robust assessment across genomic distances, predicted RE–gene pairs were stratified into seven bins ranging from 0–5 kb to 200–300 kb relative to transcription start sites (TSSs). Model performance was assessed using multiple complementary metrics, including area under Receiver Operating Characteristic (AUROC) curve, area under the precision–recall (AUPR) curve ratios, and precision, recall, and F1-score for the top-quartile (25%) of predicted interactions. Across all distance intervals, REGA consistently outperformed all baseline methods (Fig. 2c–e; Supplementary Fig. 2). Notably, despite relying solely on bulk RNA-seq data and a reference RE catalog, REGA outperformed the empirical expression-accessibility correlation baseline derived from matched single-cell multi-omics. Specifically, when evaluated against the pcHi-C ground truth, REGA achieved AUPR ratios ranging from 1.17 to 1.63 for different distance bins, corresponding to an average improvement of 27% over random prediction (Fig. 2c). In comparison, the PCC and genomic distance baselines achieved average improvements of 12% and 7%, respectively, indicating that the gain achieved by REGA was 2.25-fold and 3.86-fold greater than those of the PCC and distance baselines. Consistent performance gains were observed across independent validation using the GTEx and eQTLGen datasets (Fig. 2d–e). In the GTEx eQTL validation, REGA achieved AUPR ratios ranging from 1.22 to 1.41, with an average improvement of 31% over random prediction, compared with 16% for the PCC baseline and near-random performance for the genomic distance baseline. Similar improvements were observed in the eQTLGen validation, supporting the robustness of REGA across independent functional genetic benchmarks. Together, these results indicate that REGA captures both proximity-associated chromatin contacts and distal functional regulatory effects beyond simple genomic proximity.

To evaluate REGA’s ability to improve *trans*-regulatory inference in bulk whole blood RNA-seq data, we benchmarked its performance in predicting TF-gene interactions against two baseline methods: GENIE3^10^, a widely used and well-established method for GRN inference from expression data that has been shown to perform competitively across multiple benchmarking studies^59^, and a PCC baseline. We used ChIP-seq data for 20 TFs (Supplementary Table 1) across four blood-related cell types as ground truth^59^, defining putative TF target genes based on experimental TF-binding profiles. REGA demonstrated a substantial improvement in predictive performance, achieving an average AUPR increase of 68% over the random baseline, compared to 17% for the next best method, corresponding to an approximately fourfold relative gain. Importantly, this improvement was consistent across all 20 examined TFs in PBMCs, where REGA significantly outperformed both GENIE3 and PCC, yielding higher AUPR ratios (paired one-sided t-test, *P* < 0.0001; −log10(*P*) = 5.47 and 5.43, respectively; Fig. 2f). This advantage was further supported by additional evaluation metrics (Supplementary Fig. 3).

We next tested whether REGA identifies biologically coherent co-regulatory modules. We found modules were enriched for functionally related Gene Ontology terms (Supplementary Fig. 4a). Using a *trans*-eQTL-derived co-regulatory network from eQTLGen (where two genes are connected if they share a *trans*-SNP), 70% of REGA modules showed significant internal connectivity (Fig. 2g). REGA-derived modules also showed superior topology on this co-regulatory network, achieving a Newman’s Q modularity^60^ score of 0.146, more than double that of the next-best method, WGCNA^61^ (0.072) and other gene clustering methods. UMAP^62^ visualization of the joint latent space showed that REs, target genes, and TF motifs assigned to the same module formed tightly co-clustered groups, indicating that REGA learns a shared regulatory representation across modalities (Supplementary Fig. 4b-e). Finally, although inferred entirely from bulk data, REGA modules captured regulatory heterogeneity at single-cell resolution. Cross-referencing these modules with single-cell PBMC multiome data revealed concordant cell-type-specific patterns across module-associated gene expression and RE accessibility. Single-cell module activity analysis further confirmed population-specific regulatory patterns; for example, M9 was predominantly active in non-classical monocytes, M1 was enriched in monocytes, and M8 was specifically activated in B cells (Fig. 2j–m; Supplementary Fig. 5–6). Together, these results indicate that REGA-derived modules are functionally coherent, structurally organized, and capable of capturing cell-type-specific regulatory heterogeneity from complex bulk tissue profiles.

### REGA improves cell-type–specific GRN inference

To further evaluate REGA’s performance in cell-type-specific GRN inference, we applied it to the OneK1K cohort^63^, a large-scale single-cell RNA sequencing (scRNA-seq) dataset comprising 1.27 million peripheral blood mononuclear cells (PBMCs) from 982 donors. To assess REGA across distinct cellular contexts, we focused on three representative immune cell populations: CD4⁺ T cells, naive B cells, and CD14⁺ monocytes. These cell types were selected because matched TF ChIP-seq data were available for benchmarking regulatory interactions. Consistent with its performance in bulk RNA-seq data, REGA also reliably prioritizes functional REs in a cell-type-specific context. Comparison with ENCODE^53^ ChIP-seq data showed that the top-ranked REs identified by REGA were significantly enriched for active histone marks across these immune cell populations, with average fold changes of 1.80 for H3K27ac and 2.21 for H3K4me3 (Fig. 3a). From an evolutionary perspective, high-scoring REs in the same cell types also exhibited significantly higher sequence conservation scores than low-scoring REs across all tested percentile thresholds (10%–50%) (Fig. 3b; Supplementary Fig. 7a,b). Together, these results indicate that REGA preferentially captures functional and evolutionarily conserved regulatory elements at cell-type-specific resolution.

**Figure 3.**
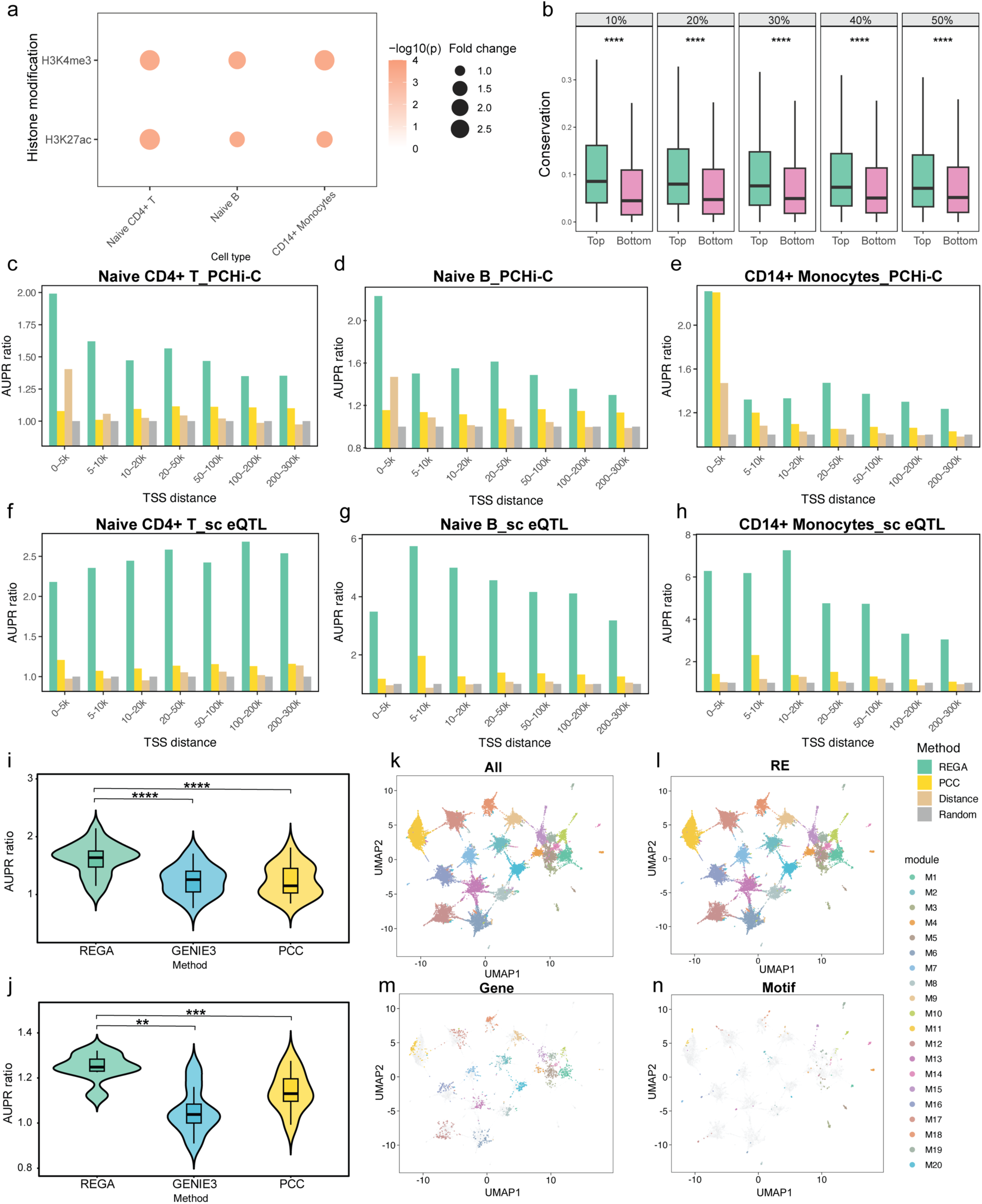
Systematic benchmarking of cell type-specific cis- and trans-regulatory inference performance. **(a)** Bubble plot showing enrichment of top-ranked REGA-inferred REs in H3K4me3 and H3K27ac ChIP-seq peaks across naïve CD4+ T cells, naïve B cells, and CD14+ monocytes. Bubble size reflects fold enrichment, and color indicates statistical significance (−log10 *P*, permutation test). **(b)** Boxplots comparing sequence conservation scores between REs with high versus low regulatory importance in naïve CD4+ T cells across percentile thresholds (10–50%; one-sided t-test, **** *P* < 0.0001). **(c–h)** Bar plots showing area under the precision–recall curve (AUPR) ratios for cell type-specific regulatory element–target gene (RE–TG) interaction prediction across seven genomic distance bins (0–5 kb to 200–300 kb), benchmarked against PCC, Distance, and Random baselines. Validation used promoter-capture Hi-C (pcHi-C) data for naïve CD4+ T cells (**c**), naïve B cells (**d**), and CD14+ monocytes (**e**), and cell type-specific eQTL data for the corresponding cell types (**f–h**). **(i)** Violin plots showing AUPR ratios for cell type-specific trans-regulatory potential prediction across 19 transcription factors (TFs), benchmarked against GENIE3 and PCC (paired one-sided t-test, **** *P* < 0.0001). **(j)** Violin plots showing AUPR ratios for predicting differentially expressed genes (DEGs) following TF perturbations in CD4+ T cells, using perturbation data from the KnockTF database (n = 8 TFs; paired one-sided t-test, ** *P* < 0.01, *** *P* < 0.001). **(k–n)** UMAP visualizations of the joint latent space of regulatory entities, including regulatory elements (REs), target genes, and transcription factor (TF) motifs, colored by module assignment. **(k)** Integrated view of all three entity types. **(l–n)** UMAP embeddings highlighting **(l)** REs, **(m)** target genes, and **(n)** TF motifs, respectively, with all unselected entities shown in light grey for structural context.

To assess the cell-type-specific *cis*-regulatory inferences of REGA, we used two independent ground truth datasets: cell-type-specific promoter-capture Hi-C^57^ (pcHi-C) profiles and single-cell eQTL^63^ (sc-eQTL) data derived from matched immune populations. We evaluated REGA’s ability to predict RE–gene interactions in CD4⁺ T cells, naive B cells, and CD14⁺ monocytes, and benchmarked it against three alternative approaches: (i) an empirical expression-accessibility PCC baseline derived from an independent 10x Genomics PBMC multiome dataset, (ii) a genomic distance decay model, and (iii) a random baseline. Across all three immune cell types, both ground truth datasets (pcHi-C, Fig. 3c–e; eQTL, Fig. 3f–h), and all tested TSS distance intervals, REGA consistently outperformed all baseline methods (Fig. 3c–h; Supplementary Fig. 7c–n). Specifically, when evaluated against the pcHi-C ground truth, REGA achieved AUPR ratios ranging from 1.35–1.99 in CD4⁺ T cells, 1.30–2.23 in naive B cells, and 1.24–2.31 in CD14⁺ monocytes (Fig. 3c–e). Similar performance gains were observed when using cell-type-matched eQTL datasets as ground truth (Fig. 3f–h), with AUPR ratios ranging from 2.18–2.68 in CD4⁺ T cells, 3.18–5.74 in naive B cells, and 3.05–7.26 in CD14⁺ monocytes. By comparison, the alternative baseline approaches exhibited limited predictive capabilities. The random baseline established the expected 1.0 performance threshold, with the genomic distance model performing only marginally above this level. Furthermore, while the empirical PCC baseline leveraged independent single-cell multiome data, it yielded only moderate improvements and remained substantially lower than REGA’s performance across most intervals.

To evaluate REGA’s performance in cell-type-specific *trans*-regulatory inference, we benchmarked its ability to predict TF–gene interactions against two established methods, GENIE3 and PCC. We first utilized putative TF targets derived from experimental ChIP-seq^64^ data (Supplementary Table 1) specific to CD4⁺ T cells, naive B cells, and CD14⁺ monocytes (n = 19 TFs) as ground truth. For the canonical CD4⁺ T cell regulator FOXP3^65^, REGA achieved an AUPR ratio of 1.73, compared to 0.76–1.06 for the baseline methods (Fig. 3i). A similar improvement was observed for ETS1, where REGA reached an AUPR ratio of 2.14, compared to 0.89–1.24 for the baselines. On average, both GENIE3 and PCC achieved an AUPR improvement of 0.24 over the random baseline, whereas REGA achieved an improvement of 0.62, corresponding to an approximately 2.8-fold increase. Consistent with these observations, REGA outperformed the baseline methods across all 19 transcription factors, yielding significantly higher AUPR ratios (paired one-sided Student’s *t*-test, *P* < 0.01 and *P* < 0.001; −log10(*P*) = 3.63 and 2.59, respectively; Fig. 3i). This advantage was observed consistently across multiple evaluation metrics (Supplementary Fig. 8a-d). To evaluate cell-type-specific *trans*-regulatory inference beyond physical binding, we next benchmarked REGA using TF perturbation data^64^ (Supplementary Table 2) as ground truth. Eight CD4⁺ T cell–specific datasets were collected from the KnockTF database^64^. As shown in Fig. 3j and Supplementary Fig. 8e-h, REGA consistently outperformed the baseline methods across all datasets.

We next examined the structural organization of cell-type-specific REGA-derived modules by visualizing the joint latent space of regulatory entities, including REs, target genes, and TF motifs, using UMAP. In this integrated representation, entities assigned to the same module formed tightly co-clustered groups, indicating that REGA learns a shared cross-modal regulatory representation (Fig. 3k; Supplementary Fig. 9). When individual entity types were highlighted within the joint manifold, REs (Fig. 3l), target genes (Fig. 3m), and TF motifs (Fig. 3n) consistently exhibited module-specific clustering patterns aligned with the global structure. Together, these results indicate that REGA integrates *cis*-REs, gene expression, and TF motif information to identify coherent co-regulatory modules.

### REGA identifies cell-type-specific disease-associated regulatory programs

To test whether REGA captures disease-relevant regulatory variation, we applied it to single-nucleus RNA-sequencing (snRNA-seq) data from PsychENCODE Phase II, which profiles the human prefrontal cortex across major neuropsychiatric disorders^66^. The analysis comprised 17 cell types in schizophrenia (SCZ), 16 in bipolar disorder (BIP) and 13 in autism spectrum disorder (ASD). Here, we use REGA to analyze each cell type and disorder separately. REGA-inferred TF activity was more sensitive than TF differential expression for identifying regulatory perturbations. For example, in SCZ L2.3.IT neurons, only five TFs were differentially expressed, whereas REGA identified 34 dysregulated TF activities (Fig. 4a,b), indicating that TF abundance alone is an incomplete proxy for regulatory function. We therefore defined driver TFs as factors whose REGA-inferred activity was significantly associated with disease status within each cell type. Across disorders, REGA identified cell-type-specific driver TFs with both increased and decreased activity (Fig. 4c; Supplementary Fig. 12a, b; Supplementary Table 3-5). In SCZ, these included known disease-relevant TFs such as EGR1, POU3F2, EGR3, JUN, and TCF4^67–73^. Notably, REGA localized TCF4, a reproducible SCZ risk locus^72,73^, to L6b neurons and Lamp5 interneurons, where it showed opposite disease-associated activity patterns (Fig. 4c). Given the role of L6b neurons in cortical circuit organization and higher-order thalamocortical communication^74,75^, this result places a canonical SCZ risk gene into a defined cellular context that is consistent with neurodevelopmental models of disease^76,77^.

**Figure 4.**
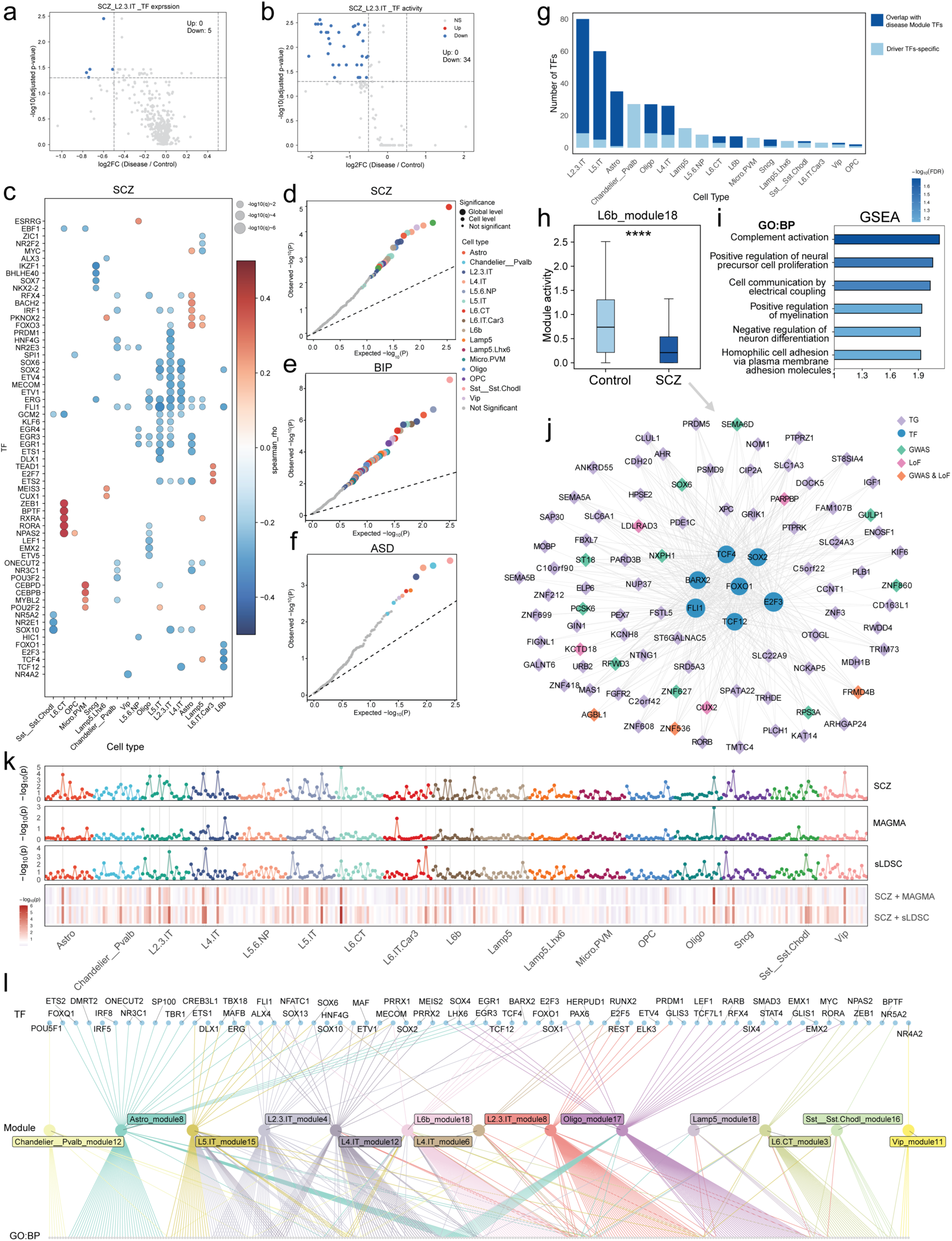
REGA identifies cell type-specific disease regulatory modules. (a–b) Volcano plots comparing transcription factor (TF) dysregulation between schizophrenia (SCZ) and control samples in L2.3.IT cells, based on **(a)** direct TF expression and **(b)** REGA-inferred TF activity. REGA-inferred activity captures substantially more disease-associated regulatory changes (Wilcoxon rank-sum test, FDR < 0.05, |log2FC| > 0.5). **(c)** Dot plot showing representative cell-type-specific driver TFs in SCZ. Dot color represents the Spearman correlation coefficient between TF activity and disease status, and dot size reflects statistical significance (−log10 FDR; global FDR < 0.1). **(d–f)** Q–Q plots showing Spearman correlation *P*-values between REGA-derived module activities and disease status for schizophrenia (SCZ; **d**), bipolar disorder (BIP; **e**), and autism spectrum disorder (ASD; **f**). Dot size denotes significance tier (global FDR < 0.05, cell-level FDR < 0.05, or non-significant), and color indicates cell type. **(g)** Stacked bar plot showing overlap between cell-type-specific driver TFs and TFs assigned to disease-associated modules across cell types. **(h)** Box plot showing significantly reduced activity of L6b module 18 in SCZ compared with control samples (two-sided Wilcoxon rank-sum test, **** *P* < 0.0001). **(i)** Bar chart showing representative Gene Ontology Biological Process (GO:BP) terms enriched in L6b module 18 identified by Gene Set Enrichment Analysis (GSEA), highlighting neurodevelopmental, cell communication, and immune-related pathways. **(j)** TF–target gene (TG) regulatory network of L6b module 18, with TGs colored according to genetic evidence: GWAS (teal), loss-of-function (LoF; pink), GWAS & LoF (orange), or standard TGs (purple). **(k)** Multi-track plot integrating module–disease associations with MAGMA and stratified LD score regression (sLDSC) genetic evidence across cell types; heatmaps show Fisher-combined significance scores. **(l)** Tripartite network linking driver TFs, genetically supported disease modules, and enriched GO:BP terms in SCZ.

We next assessed whether REGA-derived module activities were associated with disease status. REGA identified widespread disease-associated modules across disorders, including 29 modules in SCZ, 70 in BIP, and 10 in ASD (Fig. 4d–f; Supplementary Fig. 10). These modules showed cell-type-specific activity changes between disease and control samples (Supplementary Fig. 11). Gene set enrichment analysis further showed that many disease-associated modules exhibited coherent functional enrichment patterns (Supplementary Table 6-8). Driver TFs also strongly overlapped with TFs assigned to disease-associated modules (Fig. 4g; Supplementary Fig. 12c, d). For example, in SCZ L2.3.IT neurons, 94% of driver TFs were linked to disease-associated modules, supporting concordance between upstream TF dysregulation and downstream disease-associated regulatory programs.

To illustrate the mechanistic resolution of REGA, we focused on SCZ-associated L6b module 18, which contained TCF4 and showed reduced activity in SCZ relative to controls (Fig. 4h). Functional annotation linked this module to neurodevelopmental and neuronal communication processes, including neural precursor cell proliferation, electrical coupling, myelination, and complement activation (Fig. 4i). The reconstructed TF–target network was organized around central regulators including TCF4, SOX2, FOXO1, E2F3, TCF12, FLI1, and BARX2 (Fig. 4j). Importantly, many downstream targets carried independent SCZ genetic support, including SEMA6D and NXPH1 from common variant association, DLRAD3 and PARPBP from rare LoF burden, and ZNF536, FRMD4B, and AGBL1 from both evidence types. Thus, L6b module 18 links a central SCZ risk TF, disease-altered module activity, and genetically supported downstream targets within one regulatory program. A similar TF-centered architecture was observed for Astro module 8, suggesting that genetic risk also converges on broader astrocyte regulatory programs (Supplementary Fig. 13).

To further evaluate the disease relevance of REGA-derived modules, we integrated independent genetic risk evidence across cell types. For each module, we assessed three complementary sources of evidence: association between module activity and SCZ status, gene-level associations from GWAS using MAGMA^78^, and heritability enrichment of module-specific REs using stratified LD score regression^79^ (sLDSC) (Fig. 4k). Across cell types, multiple modules showed concordant signals across these analyses, with elevated −log10(*P*) values observed in the SCZ, MAGMA, and sLDSC tracks. These signals were particularly enriched in neuronal populations, indicating that modules active in these cell types are associated with both transcriptional dysregulation and genetic risk. Fisher’s combined probability test identified core modules supported by both disease association and genetic evidence (Supplementary Fig. 14a). For example, identified as disease-associated based on their regulatory activity, L6b module 18 also showed strong genetic support via sLDSC support, while Astro module 8 and Oligo module 17 demonstrated concordant evidence across multiple genetic layers (Fig. 4k). Similar patterns were observed in BIP and ASD (Supplementary Fig. 14b,c and 15), indicating that REGA modules capture disease-relevant regulatory programs supported by independent genetic evidence.

To investigate whether independent cell types share common pathological mechanisms, we constructed a hierarchical network integrating driver TFs, REGA-derived modules, and enriched Gene Ontology biological process (GO:BP) terms based on genetically supported modules from different cell types in SCZ (Fig. 4l). Multiple modules exhibited overlapping GO terms and shared driver TFs (Supplementary Fig. 16), indicating that distinct cell-type-specific modules may converge onto common regulatory processes. For example, Astro module 8 and Oligo module 17 showed shared functional enrichment and partially overlapping regulatory inputs. Similarly, modules derived from excitatory neuronal populations, including L2/3 IT, L4 IT, and L5 IT, displayed substantial overlap in both GO terms and driver TFs, suggesting coordinated regulatory programs across neuronal subtypes. These observations suggest that disease-associated regulatory modules may not be restricted to individual cell types, but instead reflect coordinated regulatory processes involving multiple cell types.

### REGA identifies across cell-type shared disease regulatory programs

To test whether neuropsychiatric disorders operate through coordinated, multicellular networks rather than isolated cell types, we sought to address two key unknowns: whether shared, cross-cell-type regulatory programs exist, and whether they are explicitly associated with disease status. We extended REGA to detect shared trans-regulatory program across nine well-sampled cell types in prefrontal cortex, testing 20 distinct modules per disorder. REGA identified disease-associated cross-cell-type activity across all three disorders evaluated: four modules in SCZ (modules 14, 20, 1, and 18), two in BIP (modules 6 and 4), and one in ASD (module 11) achieved statistical significance (FDR-adjusted *P* < 0.05; Fig. 5a–c). Quantile–quantile plots and disease-control activity comparisons showed robust disease-associated signals across multiple cell types (Fig. 5d; Supplementary Fig. 17a–f). The fact that these shared regulatory programs emerged consistently across all three distinct neuropsychiatric conditions strongly indicates that these findings are not a product of random chance, but rather reflect crucial, coordinated regulatory program successfully captured by our model.

**Figure 5.**
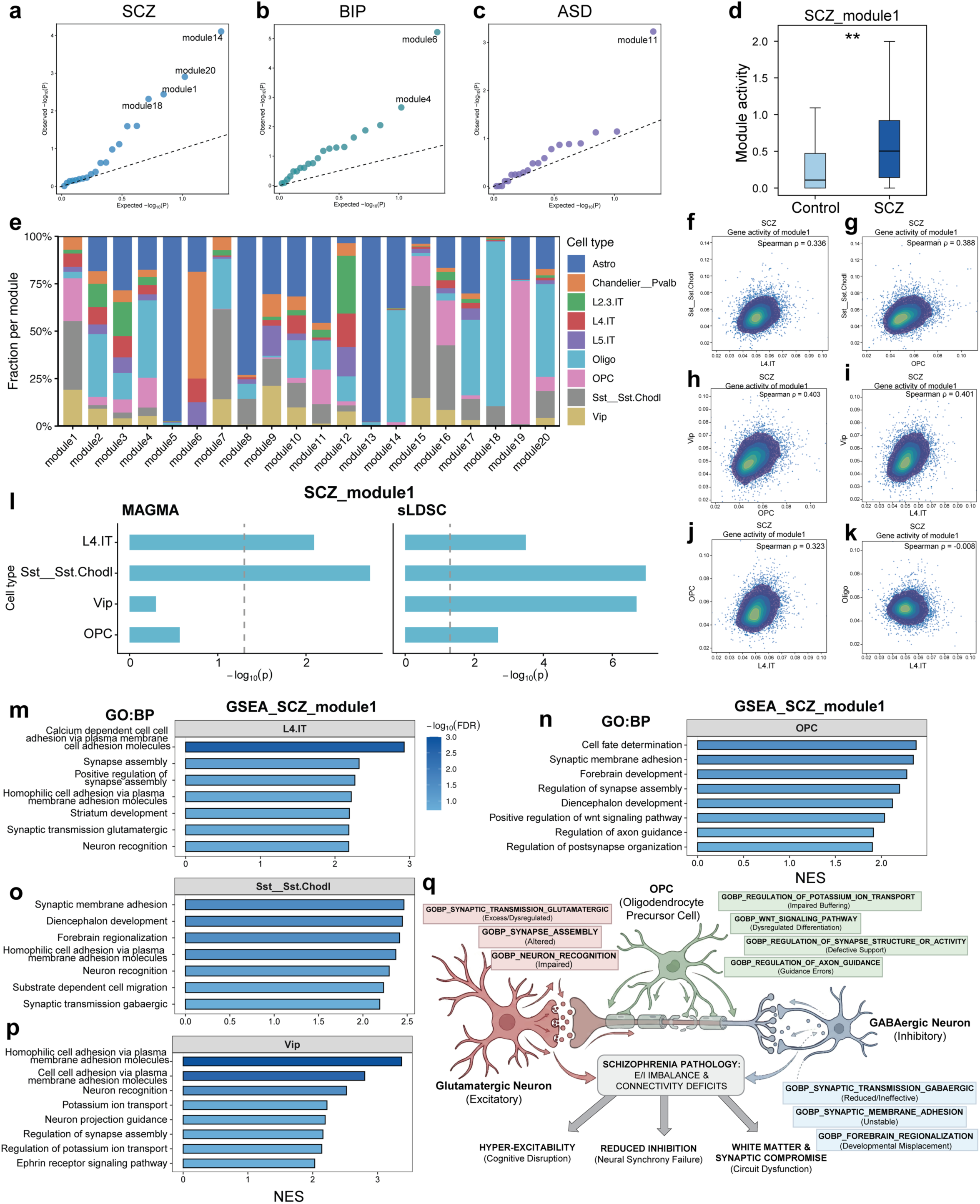
REGA identifies cross-cell-type disease regulatory programs. (a–c) Q–Q plots showing associations between cross-cell-type shared module activities and disease status for schizophrenia (SCZ; **a**), bipolar disorder (BIP; **b**), and autism spectrum disorder (ASD; **c**). Each point represents a cross-cell-type module, and the diagonal line indicates the expected null distribution. **(d)** Boxplot showing significantly increased REGA-inferred activity of SCZ module 1 in SCZ compared with control samples (two-sided Wilcoxon rank-sum test, ** *P* < 0.01). **(e)** Stacked bar plots showing the cell type composition of cross-cell-type shared modules. **(f–k)** Pairwise scatter plots comparing normalized gene loadings of shared genes in SCZ module 1 across cell types. Color gradients indicate local point density, and Spearman correlation coefficients quantify concordance across cell types. **(l)** MAGMA and stratified LD score regression (sLDSC) enrichment analyses of SCZ module 1 across cell types. Dashed lines indicate the significance threshold (*P* = 0.05). **(m–p)** Cell type-specific Gene Set Enrichment Analysis (GSEA) results for SCZ module 1 in L4.IT, OPC, Sst_Sst_Chodl, and Vip cells, highlighting schizophrenia-related biological pathways. **(q)** Conceptual model illustrating how the REGA-identified SCZ module 1 coordinates dysfunction across glutamatergic neurons, GABAergic neurons, and oligodendrocyte precursor cells (OPCs), contributing to excitation/inhibition imbalance, connectivity deficits, and circuit dysfunction in schizophrenia.

We next examined the cellular composition of shared disease-associated modules (Fig. 5e). SCZ modules showed distinct architectures: module 1 was broadly distributed across neuronal and glial populations, whereas module 14 was primarily glial, with major contributions from astrocytes and oligodendrocytes. Modules 18 and 20 were largely oligodendroglia, with additional astrocytic and other cell-type contributions. Disease-associated modules in BIP and ASD also showed mixed but structured cellular compositions, suggesting that shared regulatory programs are not restricted to a single lineage (Supplementary Fig. 17g,h).

SCZ module 1 provided a representative example of cross-cell-type regulatory organization. Its activity was significantly increased in SCZ samples relative to controls (Fig. 5d), and its gene loadings were positively correlated across multiple neuronal and glial cell-type pairs (Spearman’s ρ = 0.323–0.403; Fig. 5f–j). However, this coordination is context-dependent; for instance, module activity correlated positively between excitatory neurons and oligodendrocyte precursor cells (OPCs), but not mature oligodendrocytes (ρ = −0.008; Fig. 5k). Independent genetic evidence from MAGMA (gene-level) and sLDSC (RE-level) confirmed that the genetic risk associated with module 1 is distributed across cell types via distinct molecular mechanisms (Fig. 5l). This analysis revealed consistent genetic signals across multiple cell types. In particular, regulatory programs in L4 IT and Sst_Sst_Chodl neurons showed enrichment in both MAGMA and sLDSC analyses, suggesting genetic support from both coding and non-coding variants. In contrast, Vip neurons and OPCs showed limited enrichment in MAGMA but exhibited stronger signals in sLDSC. Together, these patterns suggest that genetic risk associated with SCZ module 1 is distributed across cell types and may involve distinct contributions from coding and regulatory variation.

Functional dissection of SCZ module 1 via cell-type-specific Gene Set Enrichment Analysis (GSEA) revealed a striking pattern of functional complementation, demonstrating how a single shared module orchestrates lineage-specific processes to achieve a unified pathological effect (Supplementary Table 9). This analysis revealed a clear pattern of functional complementation, indicating that this shared regulatory module is linked to distinct, lineage-specific processes (Fig. 5m–p). Notably, the module exhibited consistent cross-lineage enrichment for structural connectivity programs, with terms such as cell-cell adhesion, synaptic membrane adhesion, and neuron recognition ranking among the top enrichments across multiple neuronal types (Fig. 5m, o, p). Beyond this shared structural foundation, the module was associated with distinct functional pathways: in excitatory L4 IT neurons, it primarily involved glutamatergic synaptic transmission and synapse assembly (Fig. 5m), whereas in inhibitory interneurons, it highlighted pathways related to GABAergic transmission, potassium ion transport, and projection guidance (Fig. 5o, p). Concurrently, within OPCs, the module was enriched for developmental and supportive programs, including Wnt signaling, cell fate determination, and axon guidance (Fig. 5n). Synthesizing these diverse enrichments, we propose a conceptual model of cross-cell-type cooperative pathogenesis associated with SCZ module 1 (Fig. 5q). In this framework, the shared regulatory program links widespread alterations across the cortical circuit: glutamatergic hyper-excitability, diminished GABAergic inhibition, and OPC-mediated synaptic and white matter changes. Together, these cell-type–specific alterations likely contribute to the excitation/inhibition (E/I) imbalance and structural connectivity deficits characteristic of schizophrenia^80–82^.

To further characterize the diversity of regulatory architectures, we examined SCZ module 14 as a complementary, glial-dominant case that contrasts with the broader cross-lineage structure of module 1. Module 14 showed a positive astrocyte–oligodendrocyte correlation but weak correlations with neuronal populations (Supplementary Fig. 18a–c). Consistent with this pattern, the genetic signals associated with module 14 differed from those observed for module 1. While astrocyte and oligodendrocyte partitions showed limited enrichment in MAGMA, they exhibited enrichment in sLDSC analysis (Supplementary Fig. 18d), suggesting a contribution from regulatory variation. GSEA further revealed cell-type-specific patterns within this module. In astrocytes, enrichment was observed for excitatory ligand-gated ion channel activity and sodium channel activity (Supplementary Fig. 18e,g), whereas in oligodendrocytes, enrichment was observed for inhibitory ligand-gated ion channel activity, GABA-gated chloride channels, and voltage-gated calcium channel activity (Supplementary Fig. 18f,h). These findings suggest that SCZ module 14 may coordinate complementary glial processes, with astrocytes and oligodendrocytes contributing to excitatory- and inhibitory-related functions, respectively, providing a potential glial parallel to the excitation/inhibition imbalance implicated in schizophrenia^82–84^. More broadly, the identification of similar cross-cell-type modules in BIP and ASD (Fig. 5b, c) suggests that such cross-lineage regulatory patterns may represent a shared feature across complex neuropsychiatric conditions.

### REGA connects cell-intrinsic regulatory programs with intercellular communication

We applied REGA to spatial transcriptomics data to evaluate its ability to identify region-specific regulatory modules, utilizing the gene expression modality of P22 mouse brain spatial RNA–ATAC dataset^85^. We assessed module importance by calculating module contribution scores, which represent the overall regulatory signal for each module (See Methods for detail). REGA identified ten modules, of which seven showed high contribution scores and clear spatial organization, with Moran’s I values up to 0.79 (Fig. 6a; Supplementary Fig. 19, 20a). In contrast, modules 2, 5, and 9 had low contribution scores and weak spatial autocorrelation. Thus, although REGA does not use spatial neighborhood information, it recovered spatially coherent regulatory domains.

**Figure 6.**
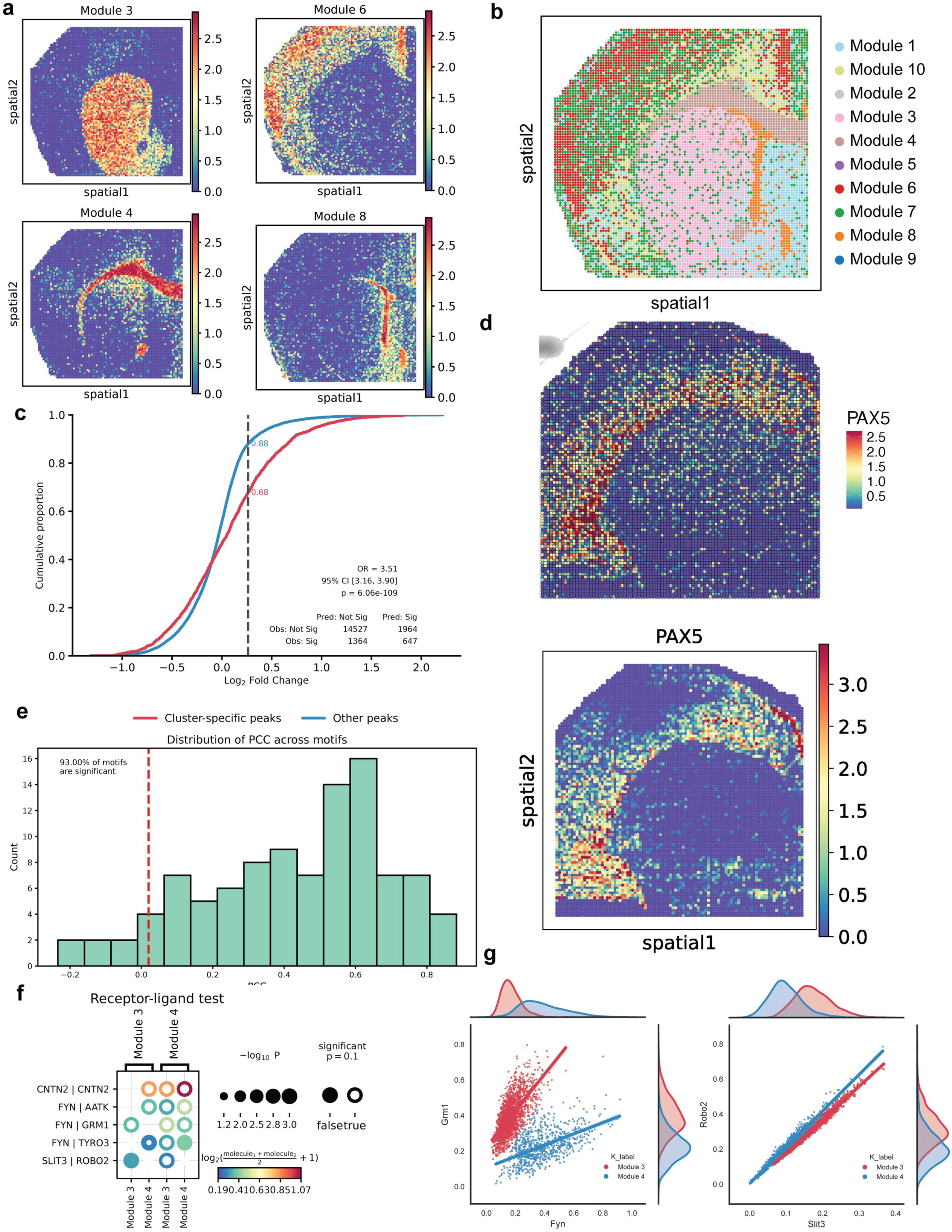
REGA captures regulatory modules in spatial transcriptomic data. **(a)** Spatial visualization of REGA module activity scores in a postnatal day 22 (P22) mouse brain coronal section, showing distinct spatial patterns for modules 3, 4, 6, and 8. **(b)** Spatial distribution of module assignments across the tissue, with spots colored by their dominant module label based on relative module proportions (n = 9,215 spots). **(c)** Empirical cumulative distribution function (ECDF) of log2 fold change (log2FC) in model- predicted chromatin accessibility for RNA cluster-specific peaks (red), defined as peaks correlated with at least one gene within 300 kb (PCC > 0.1, FDR < 0.05, log2FC > log2(1.2)), versus background non-cluster-specific peaks (blue; n = 1,682 peaks). The dotted line indicates the log2(1.2) threshold. Cluster-specific peaks showed enriched accessibility, with a pooled odds ratio of 3.51 (95% CI: 3.16–3.90) between predicted and observed patterns. **(d)** Spatial maps of Pax5 transcription factor (TF) activity inferred by chromVAR using predicted spatial ATAC profiles (top) and by REGA-inferred TF activity (bottom). **(e)** Distribution of Pearson correlation coefficients (PCCs) between REGA-inferred TF activity and chromVAR scores for the top 10 TFs per module. The red vertical line indicates the FDR significance threshold (α = 0.05), defined as the minimum PCC among significant TFs. **(f)** Ligand–receptor interactions between modules 3 and 4. Dot size represents −log10(*P*), and color indicates interaction strength (log-transformed mean expression of ligand–receptor pairs). Significant interactions (*P* < 0.1) highlight module-specific signaling. **(g)** Scatter plots showing expression levels of indicated gene pairs across spots from module 3 (red) and module 4 (blue), with linear regression fits, 95% confidence intervals, and marginal histograms showing gene expression distributions.

Module activity scores further linked these regulatory domains to known brain anatomy. Assigning each spot to its highest-activity module revealed strong agreement with Allen Brain Atlas annotations: modules 10, 7, and 6 mapped mainly to cortical regions, module 3 to the striatum, module 4 to the corpus callosum, module 8 to fiber tract/lateral ventricle-associated regions, and module 1 to the lateral septal nucleus (Fig. 6b; Supplementary Fig. 20b). GSEA supported these assignments; for example, module 8 was enriched for neural progenitor and neural stem cell processes, consistent with the progenitor-rich lateral ventricular region (supplementary table 10).

We next asked whether REGA could recover chromatin accessibility patterns without using epigenomic data as input. At the individual peak level, 95% of 1,432 module-assigned peaks (normalized module weight > 0.5) showed significant positive correlations between REGA-inferred chromatin activity and observed ATAC accessibility across spatial spots (Supplementary Fig. 21a). To determine if REGA captures the cell type specific open chromatin regions, we analyzed the consistency of its inferred chromatin activity across 11 cell types obtained from RNA information in the original study. We focused on 1,682 peaks that were correlated with at least one gene within a 300 kb window (correlation > 0.1) and are “cluster-specific” in at least 1 cluster, totaling 18,502 peak-cluster pairs. We defined a peak-cluster pair as “cluster-specific” if it exhibited a fold change > 1.2 and a significant difference relative to other clusters (FDR < 0.05, two-sample *t*-test). In the experimental ATAC data, 2,011 pairs were identified as cluster-specific, while 2,611 pairs met these criteria in REGA’s inferred chromatin activity. Notably, 647 pairs showed significant overlap between the two datasets, representing a strong enrichment (OR = 3.51, 95% CI: 3.16–3.90; Fig. 6c). This substantial agreement confirms that REGA’s inferred activity successfully recapitulates cell type and region-specific accessibility changes, even without utilizing any epigenomic or ATAC-seq data as input.

To assess whether REGA captures spatial patterns of TF activity, we selected the top 10 motifs from each module and computed the PCC between chromVAR deviation scores, based on observed ATAC data, and the corresponding motif activity inferred by REGA. The distribution of PCC values indicated predominantly positive correlations across motifs, with 93 out of 100 showing statistically significant positive associations (FDR< 0.05) (Fig. 6e, Supplementary Fig. 21b). For example, the PAX5 motif from module 10 exhibited strong spatial agreement between inferred activity and chromVAR deviation (PCC=0.71, Fig. 6d). Similar patterns were observed for additional motifs across different modules (Supplementary Fig. 22). These results demonstrate that REGA is able to capture spatially resolved TF activity patterns, supporting its effectiveness in modeling transcriptional regulatory landscapes.

Furthermore, we hypothesized that spatially adjacent modules may interact via ligand–receptor signaling. We focused on modules 3 and 4, which are spatially adjacent, and identified genes enriched in each module to examine potential interactions. This analysis revealed several significant ligand–receptor pairs (FYN–GRM1, FYN–AATK, FYN–TYRO3, SLIT3–ROBO2, CNTN2–CNTN2), suggesting inter-module communication (Fig. 6f). Prior studies support these interactions, including roles for FYN–TYRO3 signaling in oligodendrocyte myelination and CNTN2 in axon guidance^86,87^. We next examined the co-expression patterns of these pairs across spots in modules 3 and 4 to characterize their spatial coordination. For the FYN–GRM1 pair, we observe distinct, module-specific distributions: while the ligand and receptor were correlated within the spots of each individual module, their patterns were clearly separated into two distinct regression lines. Furthermore, FYN (ligand) expression was enriched in module 4, whereas GRM1 (receptor) was enriched in module 3, indicating a clear signaling direction from module 4 to module 3. In contrast, for the SLIT3–ROBO2 pair, the ligand and receptor were also correlated within both modules, but their distributions showed substantial overlap with less pronounced separation between the module-specific patterns. Crucially, the expression bias was reversed, —SLIT3 (ligand) was more highly expressed in module 3 and ROBO2 (receptor) in module 4,—suggesting a reciprocal signaling direction from module 3 to module 4 (Fig. 6g). Other pairs also displayed intermediate and shared signaling patterns, suggesting different types of interactions between modules (Supplementary Fig. 20c, d). Together, these results demonstrate that modules 3 and 4 engage in sophisticated, multi-directional communication using distinct ligand–receptor channels, highlighting REGA’s unique capability to decipher nuanced, spatially organized cell–cell communication programs across neighboring tissue regions.

### REGA maps perturbation responses to trait-associated regulatory architectures

To test whether REGA can recover interpretable regulatory programs from perturbation data, we applied it to the K562 Perturb-seq dataset from Replogle et al., containing 1,087 gene perturbations^30,88^. After aggregation by perturbation target, REGA decomposed the pseudobulk gene-by-perturbation matrix into regulatory modules that jointly connected perturbed regulators, downstream target gene programs, and TFs (Fig. 7a; Supplementary Table 11). This output is central: REGA does not merely infer regulator–gene associations, but organizes perturbation responses into programs in which TFs provide context between perturbed genes and downstream transcriptional effects.

**Figure 7.**
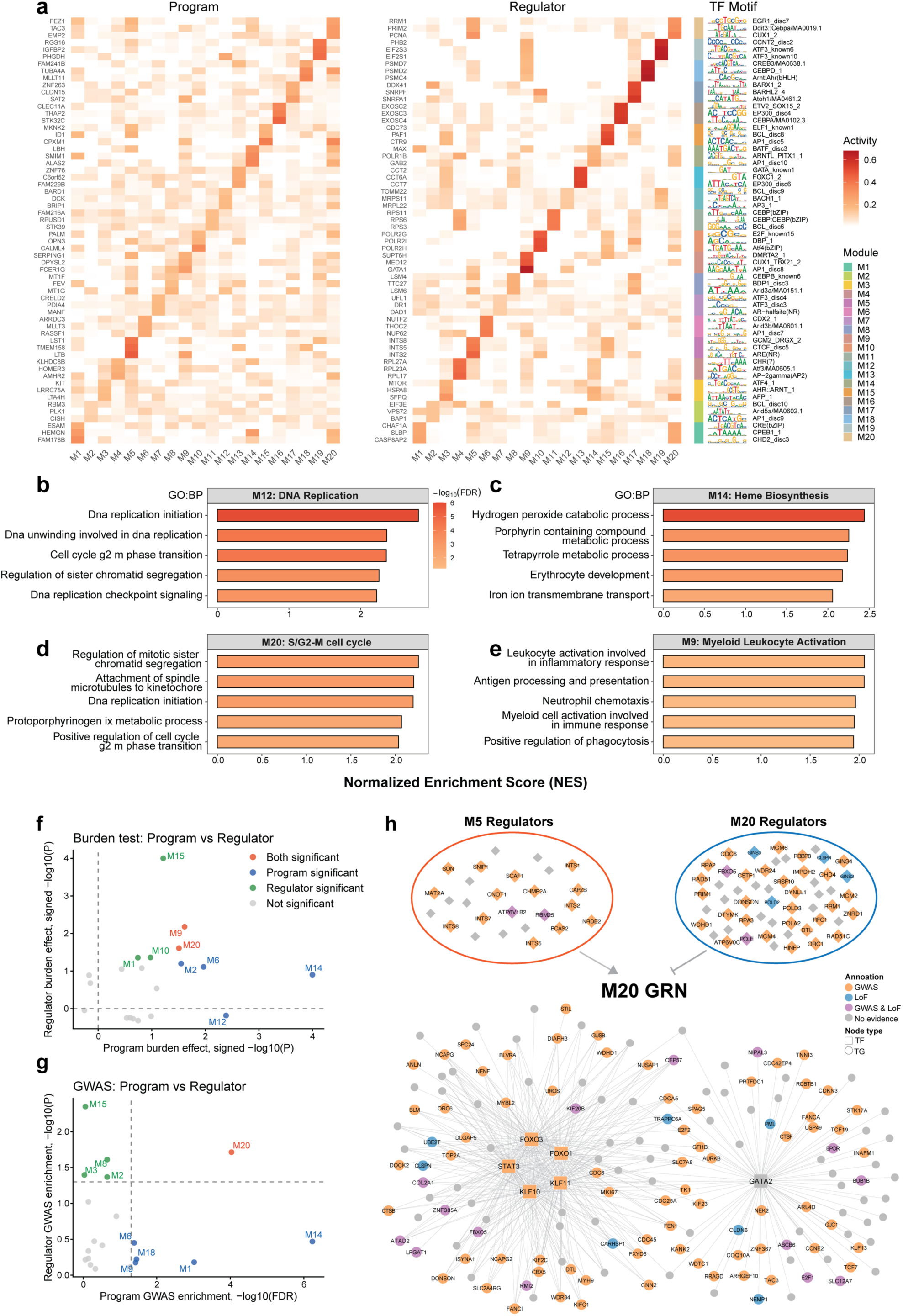
Application of REGA to perturbation data. **(a)** Heatmaps show module activities of the top three target genes (“Program”) and regulators (“Regulator”) for each REGA-derived module; darker red indicates higher activity. The right panel shows sequence logos of the top three TF binding motifs, with side colors denoting module assignment. **(b–e)** Curated functional annotation of representative modules. Bar plots show representative GO:BP terms for **(b)** M12 (DNA replication), **(c)** M14 (heme biosynthesis), **(d)** M20 (S/G2-M cell cycle), and **(e)** M9 (myeloid leukocyte activation). Bar length represents normalized enrichment score (NES), and color indicates −log10 FDR. **(f–g)** Genetic association comparison between module programs and regulators. Scatter plots compare burden effects **(f)** and MAGMA GWAS enrichment **(g)** for module target genes (“Program”) and regulators. Each point represents a module. Burden effects are signed −log10(*P*) from two-sided permutation tests (10,000 iterations). Program GWAS enrichment is shown as −log10(FDR), whereas regulator enrichment uses unadjusted −log10(*P*). Colors indicate joint significance status. **(h)** TF–TG regulatory network of the RDW-associated M20 module. Squares denote TFs and circles denote target genes. Node colors indicate RDW genetic support: grey, none; orange, MAGMA GWAS; blue, LoF burden; purple, both. Upper panels show M5 and M20 regulators connected to the M20 network, with M5 inferred to activate the M20 gene program.

The inferred modules showed biological coherence. Gene set enrichment analysis of downstream genes demonstrated that REGA separated perturbation responses into distinct functional programs (Fig. 7b–e, Supplementary Table 12-13). M12 and M20 captured different cell-cycle states: M12 was enriched for DNA replication initiation, whereas M20 was enriched for S/G2–M transition and sister chromatid segregation (Fig. 7b,d). M14 represented an erythroid program, with enrichment for heme biosynthesis and porphyrin metabolism (Fig. 7c). In contrast, M9 captured a myeloid/immune activation program involving leukocyte activation and neutrophil chemotaxis (Fig. 7e). These enrichments show that REGA partitions perturbation effects into functionally specific programs.

The TFs assigned to each module supported these annotations. M14 contained canonical erythroid TFs, including GATA1, TAL1, NFE2, and LYL1, consistent with its heme/erythroid enrichment. M5, enriched for myeloid activation, apoptosis, and stress-response processes, contained oxidative-stress-associated TFs such as JUN, JUNB, JUND, NFE2L2, and BACH1. M1 included proliferation-associated TFs such as MYC, E2F1, and E2F2, whereas M9 included immune/myeloid-associated regulators such as BHLHE40 and STAT6 (Supplementary Fig. 23a). Thus, REGA modules were supported by concordance between downstream gene function and upstream TF identity.

We next tested whether REGA modules were linked to erythroid traits: mean corpuscular hemoglobin (MCH), red cell distribution width (RDW), and immature reticulocyte fraction (IRF). We evaluated genetic support separately for downstream target gene programs and corresponding regulators following a recent study by Ota et al.^89^. Common variant signals were assessed by MAGMA-based GWAS enrichment, and rare variant effects were assessed using loss-of-function (LoF) burden test gene-level effect sizes, with significance determined by permutation testing. REGA revealed heterogeneous but structured trait-association patterns across modules (Fig. 7f–g; Supplementary Fig. 24).

In RDW, M20 showed concordant support at program and regulator layers in GWAS and LoF analyses, identifying it as a strongly RDW-associated regulatory program. M9 also showed concordant program-regulator LoF burden signals. Other modules showed layer-specific evidence. M14, the erythroid/heme module, displayed strong program-level association by GWAS enrichment, but weaker regulator-level support, suggesting that RDW-associated common variants converge mainly on the downstream erythroid program. Conversely, M15 showed stronger regulator-level signals, particularly in LoF burden and GWAS-based regulator analyses, indicating that some associations are more visible upstream. Similar layered patterns were observed for MCH and IRF. These results show that REGA can distinguish whether trait-associated genetic variation is concentrated in downstream programs, upstream regulators, or both.

TF-level genetic evidence further strengthened these conclusions. A substantial fraction of module-specific TFs showed significant MAGMA-derived GWAS associations (Supplementary Fig. 23b). In M20, five of six TFs carried significant GWAS signals for RDW, matching the strong program- and regulator-level evidence for this module. Several TFs also showed concordant common- and rare-variant support. For RDW, M15 TFs RUNX1 (LoF: −log10(*P*)=5.37; GWAS: −log10(*P*)=6.14) and SREBF2 (LoF: −log10(*P*)=4.47; GWAS: −log10(*P*)=6.73) were supported by both analyses. For MCH, E2F2 in M1 showed strong concordant evidence (LoF: −log10(*P*)=6.04; GWAS: −log10(*P*)=9.30).

Finally, the RDW-associated M20 GRN integrated these signals into an explicit regulatory architecture (Fig. 7h). M20 was organized around TFs FOXO1, FOXO3, GATA2, STAT3, KLF10, and KLF11, which connected to broad downstream target genes. All M20 TFs except GATA2 showed significant RDW GWAS evidence. Although GATA2 itself did not reach GWAS significance, many of its downstream targets carried GWAS, LoF, or combined genetic evidence, indicating that trait-associated signals can converge on downstream regulatory programs even when upstream TFs are not directly significant. Beyond intra-module effects, the network integrated cross-module input from M5, which was inferred to activate the M20 program. In REGA’s cross-module response matrix (Supplementary Fig. 23c), diagonal signals typically reflect baseline repression, where knocking down a regulator upregulates its matched gene program. Conversely, REGA utilizes off-diagonal signals to capture higher-order activation or repression between distinct modules; through this logic, the off-diagonal response between M5 and M20 revealed a clear activation effect. This cross-module link is validated by individual M5 regulators, such as ATP6V1B2 and RBM25, which carried independent GWAS and LoF evidence. Together, these structured responses demonstrate that REGA successfully links cross-module dynamics, master TFs, downstream programs, and genetic evidence into coherent, trait-associated regulatory architectures.

## Discussions and conclusions

In this study, we present REGA, an interpretable framework for inferring gene regulatory programs from gene expression data guided by reference regulatory element catalogs. A central challenge in gene regulatory network inference is that high-resolution regulatory modeling often depends on matched transcriptomic and epigenomic data, while most clinical, perturbation, and spatial transcriptomic datasets contain gene expression alone. REGA addresses this gap by using reference REs as a prior regulatory blueprint and allowing expression variation to identify active regulatory programs. This design enables REGA to model the TF–RE–target gene axis without requiring sample-level ATAC-seq.

Across benchmarking analyses, REGA improved both *cis*- and *trans*-regulatory inference. It prioritized functional REs and improved RE–gene linking despite not using ATAC profiles. Performance in pcHi-C-based evaluations decreased with genomic distance, consistent with the distance dependence of promoter-centered chromatin contacts, whereas eQTL-based evaluations remained more stable at distal intervals, as eQTLs capture functional regulatory effects on expression beyond physical contact frequency alone. REGA nevertheless outperformed all baseline methods at distal regions, including interactions more than 50 kb from the TSS (Fig. 2c–e), indicating that it captures both proximity-driven contacts and distal functional regulation beyond simple genomic proximity. For *trans*-regulation, REGA substantially improved TF–target inference relative to expression-only methods. Although TF binding and functional regulation are often discordant^90^, REGA-derived TF–target relationships were enriched for both ChIP-seq binding evidence and perturbation-defined regulatory targets. This convergence across orthogonal modalities indicates that REGA captures functionally relevant regulatory interactions beyond static TF occupancy or co-expression alone. Importantly, REGA did not only improve individual regulatory links, but also organized TFs, REs, and target genes into coherent modules. This modular structure is critical because biological regulation is rarely mediated by isolated pairwise interactions; instead, regulatory effects often emerge from coordinated programs involving multiple regulators, regulatory elements, and downstream genes.

The applications of REGA further demonstrate its broad utility. In disease snRNA-seq data, REGA detected disease-associated TF activity and regulatory modules that were not fully captured by differential expression, showing that regulatory dysregulation can occur without changes in TF abundance. In schizophrenia, REGA linked driver TFs, altered regulatory modules, and genetic risk evidence, supporting its ability to connect transcriptomic changes with upstream regulatory mechanisms and inherited disease risk. The cross-cell-type analysis further suggested that neuropsychiatric disease may arise from coordinated regulatory disruption across neuronal and glial systems, rather than from isolated changes within a single cell type.

In spatial transcriptomics, REGA recovered anatomical regulatory domains and connected spatial gene regulatory programs with ligand–receptor signaling between neighboring modules. This provides a framework for spatial niche analysis in which regulatory states within cells and communication between tissue regions can be studied together. In Perturb-seq, REGA connected perturbed genes to TF–RE–target gene programs and linked cross-module perturbation responses to trait-associated genetic evidence, demonstrating its ability to interpret perturbation effects within broader regulatory architectures.

Several limitations remain. REGA depends on the quality and relevance of the reference REs catalog and may miss condition-specific or cell-state-specific elements absent from the reference. TF activity inferred by REGA should also be interpreted as regulatory activity rather than direct biochemical activity, since TF function can be influenced by cofactors, chromatin context, and post-translational regulation. In addition, ligand–receptor interactions, genetic enrichments, and perturbation-derived regulatory links generate mechanistic hypotheses but do not by themselves prove causality.

Together, these results establish REGA as a scalable and interpretable approach for GRN inference from expression data. By integrating reference regulatory elements with transcriptomic variation, REGA lowers the data barrier for regulatory network analysis and enables mechanistic interpretation across bulk, single-cell, spatial, and perturbation datasets.

## Methods

### REGA model

REGA is a computational framework that integrates gene expression profiles with reference regulatory element data to infer the hierarchical regulatory information flow governing gene expression, spanning from latent transcriptional programs and transcription factors (TFs) to *cis*-regulatory elements (REs) and observed gene expression. At its core, REGA is based on Skeleton-Guided Interpretable Hierarchical Graph Representation Learning. The model represents transcriptional regulation through a small set of interpretable transcriptional programs, each of which is expected to capture a mechanistically supported regulatory module composed of co-expressed genes, their associated REs, and upstream TFs. To define the regulatory search space, REGA constructs a hierarchical skeleton knowledge graph following the biological flow of regulation from TFs or motifs to REs and then to genes for each cell type or tissue context (Fig. 1c). This skeleton is not treated as fixed; instead, it provides a biologically grounded initialization and constraint that can be refined during training, allowing REGA to adapt prior regulatory knowledge to the transcriptomic cohort while preserving biological plausibility.

REGA connects transcriptional programs to observed gene expression through two coupled components: the Regulatory Realization Pathway and Sample-specific Regulatory Program Activation. The Regulatory Realization Pathway uses the skeleton to define biologically plausible routes of regulation, from TFs to REs and then to genes, and learns latent embeddings for transcriptional programs, TFs, REs, and genes within this hierarchy. Dot products between regulatory node embeddings are used as attention scores to quantify regulatory compatibility: RE–gene attention scores refine the *cis*-regulatory network, and TF–gene attention scores summarize *trans*-regulatory influence on genes. In parallel, the Sample-specific Regulatory Program Activation pathway estimates the activity of each transcriptional program in each sample. These program activities generate a sample-level regulatory representation and propagate through the learned hierarchy to obtain sample-specific TF, RE, and gene activities. Finally, dot products between the sample-level representation and gene embeddings reconstruct the observed gene expression matrix, which serves as the primary training objective (Fig. 1d). Together, the two pathways impose a co-generative constraint: gene expression can be explained only when the relevant program is active in a sample and can be realized through a biologically plausible regulatory path in the skeleton. Thus, REGA is encouraged to learn mechanisms that are both predictive of expression and consistent with prior regulatory knowledge. By encoding this regulatory cascade directly into the model architecture, REGA enables mechanistic interpretability at each layer of the inferred regulatory hierarchy.

The inputs to REGA consist of a gene expression matrix after sequencing-depth correction and covariate adjustment, denoted as

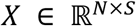

where *N* denotes the number of genes and *S* the number of samples, together with a list of *R* REs. To characterize TF binding at regulatory elements, predefined TF motifs are scanned on the RE sequences using the HOMER software, yielding a binary RE–TF motif matching matrix

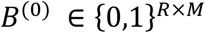

where 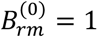 indicates that motif *m* is detected at regulatory element *r*. To bridge REs and their target genes, we construct an initial gene–RE linkage matrix

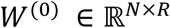

where *N* denotes the number of genes and *R* denotes the number of RE. For each gene, regulatory elements located within a 300 kb window centered at its transcription start site (TSS) are considered associated. To reflect the biological observation that proximal regulatory elements tend to exert stronger effects on gene expression, the gene–RE linkage weights are initialized using a distance-dependent exponential decay function:

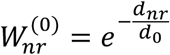

where *d_nr_* denotes the genomic distance between regulatory element *r* and the TSS of gene *n*, and *d*_0_ is a decay constant controlling the rate at which regulatory influence decreases with distance (set to 100 kb in this study). This sparse initialization guides the model toward biologically cis-regulatory interactions, with linkage weights refined during optimization.

We model each gene expression value as a weighted sum of *K* transcriptional programs:

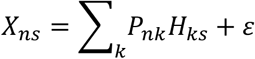

where *P_nk_*. is the contribution of program *k* to gene *n*, *H_ks_* is the activity of *k*-th program in *s*-th sample, and *ε* is Gaussian noise. The matrix *H* ∈ *R^k^*^×*s*^, referred to as the Sample-specific Regulatory Program Activation, provides a low-dimensional representation of samples in the space of transcriptional programs, and captures sample-specific regulatory states. Genes with similar program-level contributions are interpreted as belonging to the same co-regulated gene modules.

To connect transcriptional programs to the underlying regulatory architecture, REGA further models *P* through the Regulatory Realization Pathway. Each program is regulated by a set of TFs, represented by *V_mk_*., the effect of TF *m* motif on program *k*. To link programs back to genes, we model *P_nk_*. as the combined effect of nearby REs and the TFs bound to them:

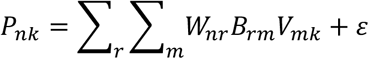

where *W_nr_*_)_ is the regulatory effect of RE *r* on gene *n*, and *ε* is Gaussian noise. All parameters are constrained to be non-negative. In matrix form, the model reduces to a four factors non-negative matrix factorization of the expression matrix

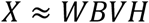

which captures how gene expression arises from the coordinated action of REs, TFs, and transcriptional programs across samples. Model parameters are learned by solving the following non-convex optimization problem.

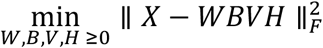

We designed a multiplicative update algorithm to solve this optimization problem. The update rules for the model variables are as follows:

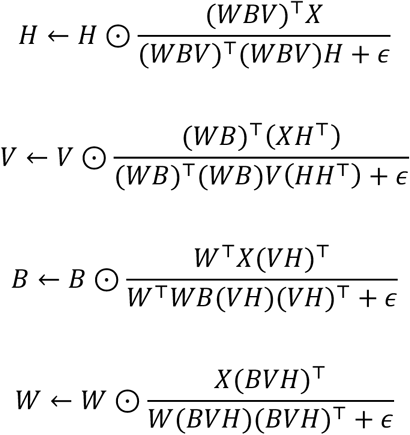

In addition to the distribution choice, the only tuning parameter is the number of transcriptional program *K*.

Formally, REGA represents gene regulation as a hierarchical regulatory network composed of multiple interconnected layers, including latent transcriptional programs, transcription factor activities, cis-regulatory elements, and gene expression. Each layer corresponds to a distinct regulatory abstraction, and regulatory information is modeled to propagate directionally across layers. Latent transcriptional programs capture coordinated regulatory activity shared across genes, giving rise to co-regulated gene modules, while transcription factors and cis-regulatory elements mediate how these programs influence gene expression. This hierarchical formulation enables REGA to learn structured regulatory representations rather than abstract latent factors.

### Multi-view extension of REGA for cross-cell-type analysis

In complex tissues, cells of different types collaborate to perform essential biological functions. While inter-individual variation in gene expression is often attributed to cell-type-specific transcriptional programs, it can also be shaped by coordinated regulatory changes across multiple cell types^91^. Such inter-cellular dependencies suggest that regulatory logic is not purely cell-autonomous, but can be synchronized across the tissue environment.

To disentangle these relationships, we reformulate REGA framework into a multi-view formulation for cross-cell-type analysis. For each cell type *i*, we construct pseudobulk data across donors and model expression as

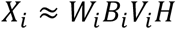

where *W_i_*, *B_i_*, and *V_i_* are cell type specific, capturing cis- and trans-regulatory components in each cell type. The donor-level embedding *H*, however, is shared across cell types and represents a cross-cell-type regulatory program space. By aggregating information across multiple cellular views, *H* provides a higher-level convergent regulatory representation with great potential to be highly informative for capturing disease phenotypes. To jointly model gene expression across cell types under this formulation, model parameters are learned by minimizing the aggregate reconstruction error across all cellular contexts, leading to the following objective function:

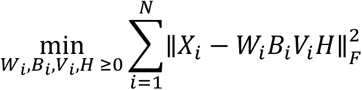

We proposed a multiplicative update algorithm to solve this joint four-factor non-negative matrix factorization model. Update of variables are as follows:

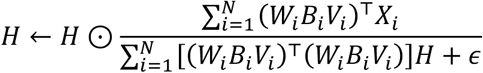

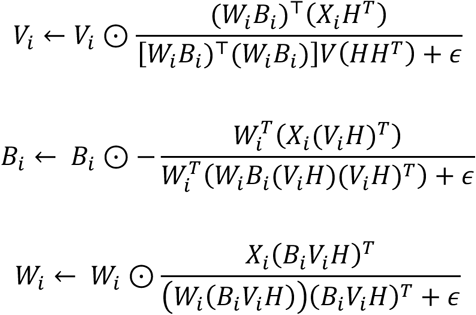

### Data collection and preprocess

Bulk RNA-seq data from whole blood were obtained from the GTEx project^52^ (https://www.gtexportal.org), and an aggregated ATAC-seq peak set derived from a public 10x Genomics peripheral blood mononuclear cells (PBMCs) multiome dataset was used as the reference set of regulatory elements (REs), downloaded from the 10x Genomics website (https://www.10xgenomics.com/datasets). OneK1K single-cell RNA-seq (scRNA-seq) data from PBMCs^63^ were downloaded from the Gene Expression Omnibus (https://www.ncbi.nlm.nih.gov/geo/) under accession number GSE196830, and the same aggregated PBMC multiome ATAC-seq peak set was used as the reference RE set. For all scRNA-seq datasets, pseudobulk expression profiles were generated by aggregating gene expression across cells within each sample, resulting in gene-by-sample matrices used as model inputs. Single-nucleus RNA-seq (snRNA-seq) data for schizophrenia (SCZ), bipolar disorder (BIP), and autism spectrum disorder (ASD) in the adult dorsolateral prefrontal cortex (DLPFC) were obtained from brainScope^66^ (Brain Single-Cell Omics for PsychENCODE Resource; https://brainscope.gersteinlab.org/), and cell type-specific reference RE sets were also obtained from brainScope, based on single-cell candidate cis-regulatory regions identified from snATAC-seq and snMultiome data. Spatial ATAC–RNA-seq data (spatial assay for transposase-accessible chromatin and RNA using sequencing) from mouse brain coronal sections at postnatal day 22 (P22) were obtained from Zhang et al.^85^ via GEO (GSE205055). The dataset was generated at a spatial resolution of 20 μm and comprised paired transcriptomic and chromatin accessibility measurements across approximately 10,000 spatial spots; the gene-by-spot matrix and corresponding reference RE set derived from this dataset were used as model inputs. Perturb-seq data from Replogle et al.^30^ in K562 cells were downloaded via GEARS^88^ (https://dataverse.harvard.edu/api/access/datafile/7458695), comprising 1,087 high-quality gene perturbations. The reference RE set for K562 cells was derived from ENCODE^53^ ATAC-seq data (https://www.encodeproject.org/). Pseudobulk profiles were generated by aggregating gene expression across cells sharing the same perturbation target, and the resulting gene-by-perturbation matrix was used as model input.

### RE importance score

To quantify the importance of each RE, we defined an RE importance score by integrating each RE’s regulatory strength toward target genes with its overall activity across the dataset. Let *W* ∈ ℝ*^N^*^×*R*^ denote the gene-by-RE regulatory strength matrix, where *N* is the number of target genes and *R* is the number of REs. For the *r*-th RE, its total regulatory strength was calculated as

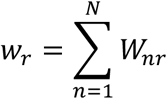

We then computed the reconstructed RE activity matrix as

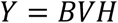

where *B*, *V*, and *H* are factor matrices learned by REGA. and *Y_rs_* represents the activity of RE *r* in sample *s*. The overall activity of RE *r*across the dataset was summarized as

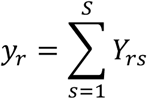

The RE importance score was then defined as

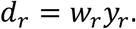

REs with larger *d_r_* values were considered to have higher regulatory importance, reflecting both stronger target-gene regulatory effects and higher overall activity across the dataset.

### Epigenetic validation of prioritized REs

To evaluate the functional relevance of regulatory elements (REs) prioritized by our model, we assessed their spatial overlap with histone modification ChIP–seq peaks (e.g., H3K27ac and H3K4me3) derived from matched cell types. We focused on the top 2,000 REs ranked by their model-inferred importance scores. To control for potential biases introduced by variable region lengths, both the prioritized REs and background ATAC–seq peaks were normalized to fixed 500-bp windows centered on their midpoints (±250 bp). Overlap significance was quantified using a permutation test with 10,000 iterations, randomly sampling 2,000 background ATAC-seq peaks per iteration to construct a null distribution. Enrichment was quantified as the fold change:

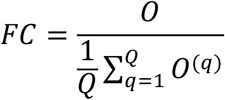

where *O* denotes the observed overlap count and *O*^(*q*)^ denotes the overlap count in the *q*-th permutation (*Q* = 10,000). Empirical P-values were computed as:

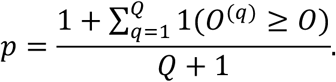

### *Cis-* and *trans*-regulatory inference using latent regulatory embeddings

To infer *cis*- and *trans*-regulatory relationships, REGA represents target genes, REs, and TF motifs in a shared latent regulatory module space. We defined latent node embeddings for genes, REs, and TF motifs as

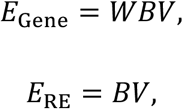

and

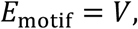

where *E*_Gene_ ∈ ℝ*^N^*^×*K*^, *E*_RE_ ∈ ℝ*^R^*^×*K*^, and *E*_motif_ ∈ ℝ*^M^*^×*K*^ . Here, *E*_Gene_ represents gene embeddings obtained by aggregating motif-module information across gene-associated REs, *E*_BC_ represents RE embeddings derived from motif composition and module membership, and *E*_motif_ represents motif embeddings in the same latent regulatory space.

*Cis*-regulatory strength between target genes and REs was defined by the dot product attention score between gene and RE embeddings:

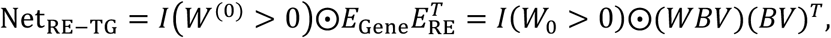

where Net_RE-TG_ ∈ ℝ*^N^*^×*R*^, *I*(*W*^(0)^ > 0) is a binary mask indicating RE–gene links in the prior skeleton. Each entry Net_RE-TG,*nr*)_ represents the inferred regulatory strength between target gene *n* and RE *r*. This formulation can be interpreted as an attention-like compatibility score, in which each gene embedding acts as a query and each RE embedding acts as a key in the latent regulatory module space. REs with embeddings more compatible with a given gene embedding receive higher inferred *cis*-regulatory scores

Similarly, motif-level trans-regulatory strength between target genes and TF motifs was defined by the dot product between gene and motif embeddings:

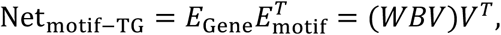

where Net_motif-TG_ ∈ ℝ*^N^*^×*M*^ . Each entry Net_TF-TG,*nt*_ represents the inferred regulatory strength between target gene *n* and TF motif *m*, after integrating gene-associated RE activity and latent motif-module structure. This score can also be interpreted as an attention-like compatibility score, where the gene embedding acts as a query and the motif embedding acts as a key in the latent regulatory module space. Motifs with embeddings more compatible with a given gene embedding receive higher inferred trans-regulatory scores. To obtain TF-level regulatory scores, motif-level scores were mapped to TFs using a motif–TF matching matrix. Let *C* ∈ ℝ*^M^*^×*T*^ denote the motif–TF matching matrix, where *T* is the number of TFs. The TF–TG regulatory strength matrix was computed as

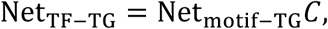

where Net_TF-TG_ ∈ ℝ*^N^*^×*T*^ . Each entry Net_IK,I?,+L_ represents the inferred regulatory strength between target gene *n* and TF *t*, after integrating RE–TG regulatory activity, motif binding information, latent regulatory module structure, and motif–TF matching.

### Module assignment

To assign regulatory entities to REGA-inferred modules, we designed a two-channel parallel assignment strategy based on REGA-derived node embeddings. Node embedding matrices were generated for different entity types, including

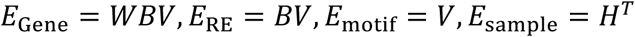

representing embeddings for genes, REs, TF motifs, and samples, respectively.

Step 1: Node embedding normalization

For each input module activity matrix

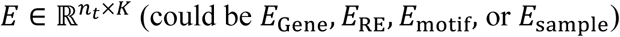

where rows represent entities and columns represent REGA modules, and *n_t_* denotes the number of entities for a given entity type, we first applied a two-step L1 normalization to make scores comparable across both modules and entities.

Specifically, column-wise normalization was performed as

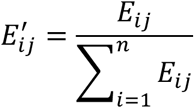

followed by row-wise normalization

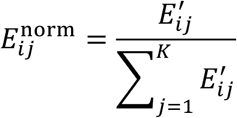

Where

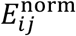

denotes the normalized score used for module assignment.

Step 2: Column-wise top-fraction channel.

For each module, entities ranked within the top user-defined fraction (e.g., top 10%) of normalized scores were selected. This channel prioritizes entities with the strongest relative contribution to each module.

Step 3: Row-wise absolute-threshold channel.

Each entity was assigned to the module with the maximum normalized score if that score exceeded a user-defined threshold (e.g., *p* = 0.25), requiring the selected module to explain at least 25% of the feature’s normalized module weight.

Step 4: Parallel union.

For each module, assignments from the top-fraction and absolute-threshold channels were combined by union to generate candidate entity–module assignments.

After candidate assignments were obtained, users could choose between mutually exclusive hard assignments and overlapping soft assignments. For hard assignment, entities assigned to multiple modules were resolved by retaining only the module with the highest normalized score, producing a one-to-one entity-to-module mapping for genes, REs, TF motifs, and samples. For soft assignment, entities were allowed to remain assigned to multiple modules when selected by more than one module. Thus, the hard assignment mode emphasizes module specificity, whereas the soft assignment mode preserves overlapping regulatory membership.

The top-fraction value and absolute threshold were user-adjustable parameters. Setting either parameter to zero disabled the corresponding channel, allowing module assignment to be performed using only the remaining channel.

For the multi-cell-type extension of REGA, the module assignment procedure remained unchanged except for the column-wise top-fraction channel. Specifically, all cell-specific normalized matrices were merged into a single global matrix, with rows indexed as *celltype_entity*. Top-fraction selection was then performed globally across all entities, allowing entities from different cell types to compete within the same module ranking.

### Identification of disease-associated driver TFs

To identify disease-associated driver TFs, we first defined sample-level TF activity by integrating motif activity, motif–TF annotation, and TF expression within each cell type. For each cell type, let *VH* ∈ ℝ*^M^*^×*S*^ denote the sample-level motif activity matrix, where *M* is the number of motifs and *S* is the number of samples. Let *X*^TF^ ∈ ℝ*^T^*^×*S*^ denote the TF expression matrix, where *T* is the number of transcription factors. A binary motif–TF matching matrix *C* ∈ {0,1}*^T^*^×*M*^was constructed from the motif–TF mapping file, where

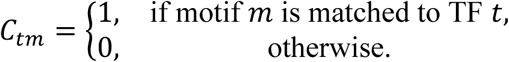

For each TF–motif pair, we calculated the Pearson correlation coefficient between TF expression and motif activity across matched samples:

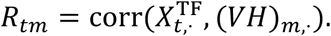

To retain TF–motif relationships supported by positive sample-level association, correlations below 0.3 were set to zero:

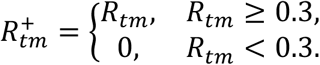

The TF–motif weight matrix was then defined as the element-wise product of the binary motif–TF matching matrix and the filtered correlation matrix:

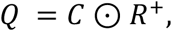

Motif-derived TF activity was calculated as the weighted sum of motif activities:

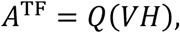

where *A*^TF^ ∈ ℝ*^T^*^×*S*^ represents the motif-derived activity score of each TF in each sample.

To incorporate both TF regulatory activity and TF expression abundance, the final sample-level TF activity score was defined as the element-wise product of TF expression and motif-derived TF activity:

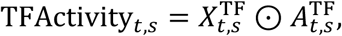

where *t* denotes a TF and *s* denotes a sample. This expression-weighted activity score was used as the TF activity measure in downstream analyses.

For disease association analysis, samples were classified according to binary disease status, with control samples coded as 0 and disease samples coded as 1. Within each cell type, Spearman correlation analysis was performed between TF activity and disease status across matched samples:

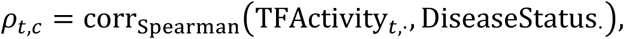

where *c* denotes the cell type. *P*-values from all tested TF–cell type pairs were jointly corrected for multiple testing using the Benjamini–Hochberg false discovery rate procedure. TF–cell type pairs with global FDR-adjusted *q* < 0.1 were defined as cell-type-specific disease-associated driver TFs.

### Evaluation metrics

The performance of gene GRN inference was evaluated using multiple complementary metrics, including the area under the precision–recall curve (AUPR), the area under the receiver operating characteristic curve (AUROC), and precision, recall, and F1-score calculated using the top 25% of predicted regulatory interactions. The AUPR ratio was defined as the ratio between the observed AUPR of a method and the expected AUPR of a random predictor. For a random predictor, the expected AUPR equals the fraction of positive samples in the dataset. Thus, the AUPR ratio was calculated as

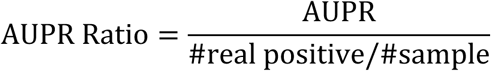

where #real positive/#samplerepresents the fraction of positive samples in the dataset. The AUPR ratio therefore quantifies the fold improvement of a predictor relative to random prediction.

### *Trans*-eQTL co-regulatory network enrichment analysis

To evaluate the biological coherence of REGA-derived gene modules, we performed network enrichment analysis using a weighted *trans*-eQTL co-regulatory network constructed from eQTLGen Consortium data. In this network, an edge was placed between two genes if they shared at least one common trans-acting single-nucleotide polymorphism (SNP) regulating both genes, with the edge weight defined as the number of shared *trans*-acting SNPs. For each REGA-derived module, internal connectivity was quantified as the sum of edge weights among genes assigned to the module within the co-regulatory network. Statistical significance was assessed using a permutation-based strategy. Specifically, 10,000 size-matched random gene sets were sampled from the same background gene pool, and their weighted internal connectivity scores were used to construct a null distribution. Empirical *P*-values were calculated as the proportion of random gene sets with weighted internal connectivity greater than or equal to the observed value. Empirical *P*-values were corrected across modules using the Benjamini–Hochberg false discovery rate procedure, and modules with FDR-adjusted *P* < 0.05 were considered significantly enriched for *trans*-eQTL co-regulatory connectivity.

### MAGMA

Single-gene and module-level GWAS enrichment analyses were performed using MAGMA^78^ v1.10. GWAS summary statistics were mapped to genes using the default MAGMA SNP-to-gene annotation procedure, and linkage disequilibrium (LD) structure was estimated using the 1000 Genomes Phase 3 European (EUR) reference panel^92^. For module-level analyses, module gene sets were defined as genes assigned to the same REGA-derived module. Benjamini–Hochberg false discovery rate (FDR) correction was applied across all tested genes for single-gene analyses and across all tested modules for module-level analyses.

### Stratified LD score regression

Partitioned heritability enrichment was evaluated using stratified LD score regression (sLDSC; v1.0.1)^79^. REGA module-specific regulatory annotations were constructed from regulatory elements (REs) assigned to modules based on the *BV* matrix, and LD scores were computed using a 1-centimorgan (cM) window with the 1000 Genomes Phase 3 European (EUR) reference panel^92^ and HapMap3 SNPs. For each GWAS summary statistic, sLDSC was performed jointly with the baselineLD v2.2 model^93^. Allele frequencies and regression weights were obtained from 1000 Genomes Phase 3 European reference resources, with regression weights based on HapMap3 SNPs excluding the major histocompatibility complex (MHC) region. Module-level heritability enrichment was quantified using regression coefficients and corresponding *P*-values from the sLDSC output.

### Module-level burden effect analysis

Following the burden effect enrichment framework described by Ota et al.^89^, genetic associations between REGA-identified modules and complex traits were evaluated. For each module, two module-assigned components were analyzed separately: program genes, defined as genes assigned to the module, and regulators, defined as perturbation target genes assigned to the same module. For both program genes and regulators, burden effects were quantified as the average loss-of-function (LoF) effect size (*γ*) of the corresponding gene set. GeneBayes^94^ posterior mean LoF effect sizes (*γ*) and selective constraint scores (*S*_het_) were obtained from Ota et al.^89^. To control for the confounding effect of selective constraint, background genes expressed in the relevant cellular context were stratified into ten bins according to their *S*_het_ scores. For each program-gene or regulator set, a null distribution was generated using 10,000 permutations. In each permutation, the same number of background genes was sampled while preserving the *S*_het_ bin distribution of the observed gene set. Statistical significance was assessed using two-sided empirical *P*-values by comparing the observed average *γ* with the corresponding null distribution. The direction of the trait association was determined by comparing the observed mean with the null mean: a significantly positive average *γ* was interpreted as a repressing effect on the trait, whereas a negative average *γ* was interpreted as a promoting effect.

### Contribution scores and selection of REs and TF motifs for spatial transcriptomics analysis

To characterize module-level contributions in spatial transcriptomics data, we quantified the contribution of each latent module using both gene-side embeddings and spot-level embedding. Let *E*_Gene_ = *WBV* ∈ ℝ*^N^*^×*K*^denote the gene embedding matrix and let *H* ∈ ℝ*^K^*^×*S*^ denote the module-by-spot embedding matrix for the spatial dataset, where *K* is the number of modules and *S* is the number of spatial spots. For each module *k*, we computed a module contribution score as the product of the total gene-side contribution and the total spatial activity of that module:

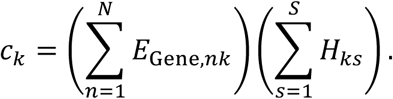

Equivalently, the vector of module contribution scores was computed as the diagonal of the outer product between the column sums of *E*_Gene_ and the row sums of *H* . Modules with higher *c*. values were considered to make larger contributions to the spatial regulatory architecture.

To assess whether REGA recovered chromatin accessibility patterns in spatial data, we selected module-associated REs from the RE embedding matrix *E*_BC_ = *BV*. REs were selected using the row-wise absolute-threshold channel of the module assignment strategy described above. Specifically, an RE was retained if its maximum normalized module-assignment score exceeded the specified threshold. This procedure yielded approximately 1,432 module-associated REs for downstream validation.

To prioritize TF motifs for spatial regulatory interpretation, we first selected TF motifs using the same row-wise absolute-threshold channel applied to the motif embedding matrix *E*_motif_ = *V*. We further ranked TF motifs within each module using a motif-by-module weighted score matrix. Let *s* ∈ ℝ^<×(^ denote a vector of motif abundance scores obtained by summing the RE-by-motif binding matrix *B* across REs:

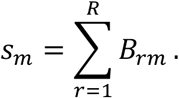

The weighted motif-by-module score matrix was then computed as

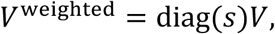

where *V*^weighted^ ∈ ℝ*^M^*^×*K*^. TF motifs were ranked within each module according to *V*^weighted^, and the top 10 motifs per module were selected for downstream analysis.

#### GSEA

Gene set enrichment analysis (GSEA)^95^ was performed using the preranked mode implemented in the Python package GSEApy^96^. For each module, genes were ranked according to their module-specific activity scores in the *WBV*(gene-by-module) matrix. Prior to analysis, the *WBV* matrix was subjected to L1 normalization across columns followed by rows. Ranked gene lists for each module were then used as input for preranked GSEA. Enrichment significance was evaluated using normalized enrichment scores (NES), nominal P-values, and Benjamini–Hochberg false discovery rate (FDR)-adjusted q-values.

## Data availability

All datasets used in this study are publicly available. Bulk RNA-seq data from whole blood were obtained from the GTEx project^52^ (https://www.gtexportal.org). Single-cell RNA-seq (scRNA-seq) data from the OneK1K cohort are available from the Gene Expression Omnibus (GEO; https://www.ncbi.nlm.nih.gov/geo/) under accession number GSE196830^63^. Single-nucleus RNA-seq (snRNA-seq) data for schizophrenia (SCZ), bipolar disorder (BIP), and autism spectrum disorder (ASD) in the adult DLPFC, along with corresponding candidate cis-regulatory regions, were obtained from brainScope^66^ (https://brainscope.gersteinlab.org/). Spatial ATAC–RNA-seq data from mouse brain coronal sections at postnatal day 22 (P22) were collected from GEO under accession number GSE205055^85^. Perturb-seq data in K562 cells from Replogle et al.^30^ were downloaded via GEARS^88^ (https://dataverse.harvard.edu/api/access/datafile/7458695), and K562 ATAC-seq data were obtained from ENCODE^53^ (https://www.encodeproject.org/). The PBMC multiome data were obtained from the 10x Genomics website (https://www.10xgenomics.com/datasets). Histone modification data for H3K4me3 and H3K27ac across multiple blood-related cell types, including B cells, CD4⁺ and CD8⁺ T cells, monocytes, neutrophils, and NK cells, were obtained from the ENCODE consortium^53^. Promoter-capture Hi-C data were obtained from Javierre et al.^57^. Whole blood eQTL data were obtained from the GTEx^52^ consortium and eQTLGen^58^ consortium (https://www.eqtlgen.org/). Single-cell eQTL data were obtained from the OneK1K cohort^63^ (https://onek1k.org/). TF ChIP-seq data were obtained from the Cistrome database^59^ (http://cistrome.org/), and TF perturbation data were obtained from the KnockTF^64^ database (http://www.licpathway.net/KnockTF/index.html). GWAS summary statistics for schizophrenia, autism spectrum disorder, and bipolar disorder were downloaded from the Psychiatric Genomics Consortium (PGC)^97–99^ (https://pgc.unc.edu/for-researchers/download-results/). GWAS summary statistics for blood traits were obtained from the GWAS Catalog^100^ (https://www.ebi.ac.uk/gwas/). Loss-of-function (LoF) burden test summary statistics from the UK Biobank^101^ were also obtained from the GWAS Catalog^100^.

## Code availability

The software is available at GitHub (https://github.com/Durenlab/REGA).

## Acknowledgments

This work was partially supported by NIH grants R35GM150513 and R21DA060503.

---

**Supplementary Figure 1.**
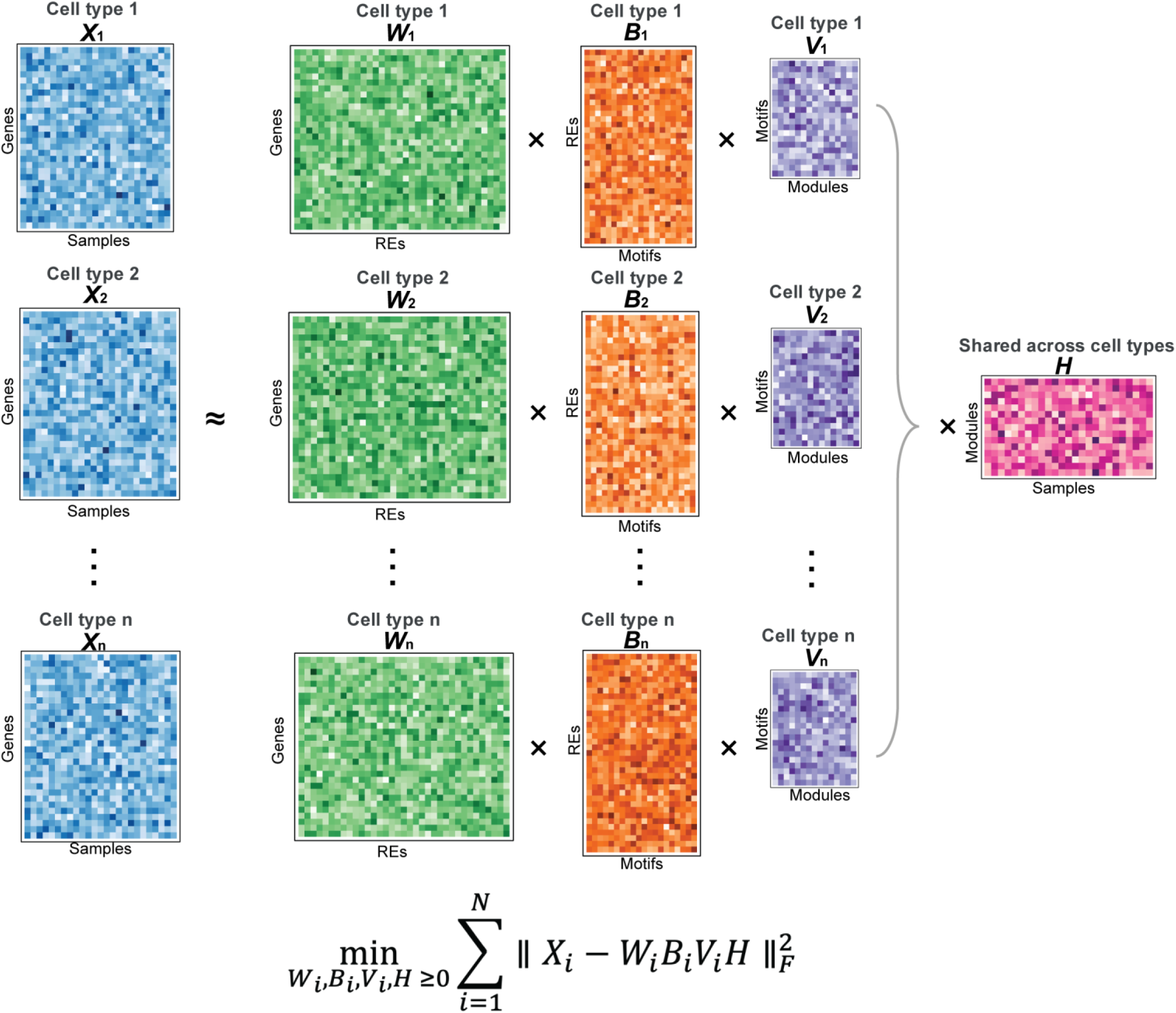
Multi-view extension of REGA for cross-cell-type regulatory program inference. Schematic overview of the multi-view REGA framework for cross-cell-type analysis. For each cell type *i*, pseudobulk gene expression profiles across samples (*Xᵢ*) are modeled using cell type-specific cis- and trans-regulatory components, including regulatory element (RE)–gene associations (*Wᵢ*), TF motif binding profiles (*Bᵢ*), and motif–module representations (*Vᵢ*). A shared latent embedding (*H*) is jointly learned across cell types to capture sample-level regulatory programs shared across cellular contexts. By integrating multiple cellular views, the model minimizes the aggregate reconstruction error across cell types, enabling the identification of coordinated cross-cell-type regulatory programs. The corresponding optimization objective is shown below.

**Supplementary Figure 2.**
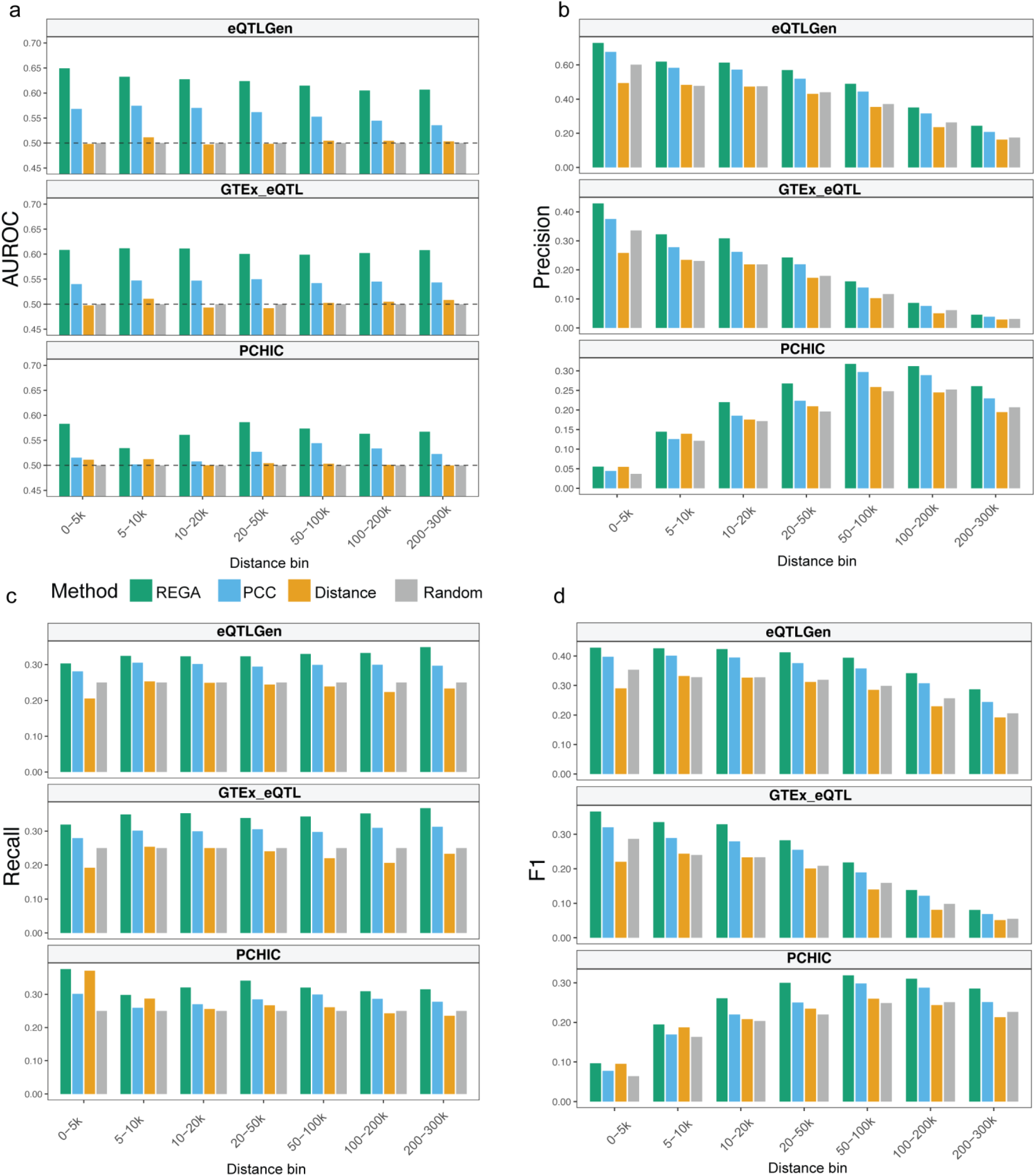
Performance assessment of cis-regulatory strength inferred by REGA in bulk RNA-seq data. Bar plots show the performance of REGA in predicting regulatory element–target gene (RE–TG) interactions across genomic distance bins (0–5 kb to 200–300 kb relative to the transcription start site (TSS)) using GTEx whole-blood bulk RNA-seq data. Performance was evaluated against three independent ground-truth datasets: eQTLGen eQTLs, GTEx whole-blood eQTLs, and promoter-capture Hi-C (pcHi-C), shown as separate rows in each panel. REGA was benchmarked against three alternative methods: PCC (Pearson correlation between RE chromatin accessibility and target gene expression, computed using an independent 10x Genomics PBMC multiome dataset), Distance (a genomic distance-based decay function), and Random (a uniform baseline). Panels show **(a)** area under the receiver operating characteristic curve (AUROC), **(b)** precision, **(c)** recall, and **(d)** F1-score. Precision, recall, and F1-score were evaluated using the top quartile (25%) of predicted RE–TG interactions within each distance bin. The dashed line in **(a)** indicates the expected AUROC of a random classifier (0.5).

**Supplementary Figure 3.**
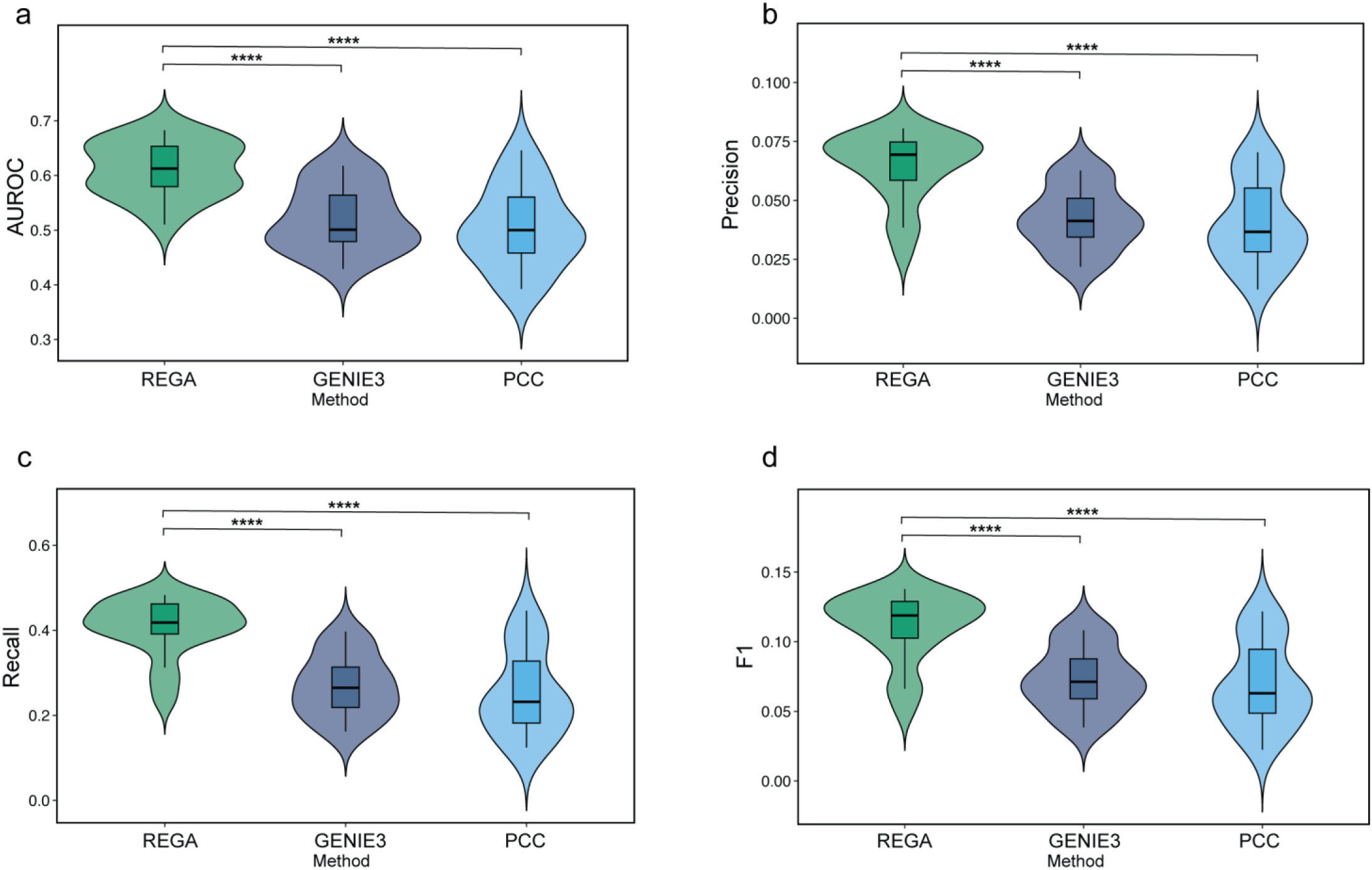
Performance assessment of trans-regulatory inference by REGA in bulk RNA-seq data. Violin plots show the performance of REGA for predicting trans-regulatory targets across transcription factors (TFs), benchmarked against GENIE3 and PCC. Ground-truth TF–target interactions were defined using experimental ChIP-seq data from blood-related cell types. Panels show **(a)** area under the receiver operating characteristic curve (AUROC), **(b)** precision, **(c)** recall, and **(d)** F1-score. Precision, recall, and F1-score were computed using the top quartile (25%) of predicted TF–target interactions. Boxplots within violins indicate the median and interquartile range. Statistical significance was assessed using paired one-sided Student’s *t*-tests (n = 20 TFs; **** *P* < 0.0001).

**Supplementary Figure 4.**
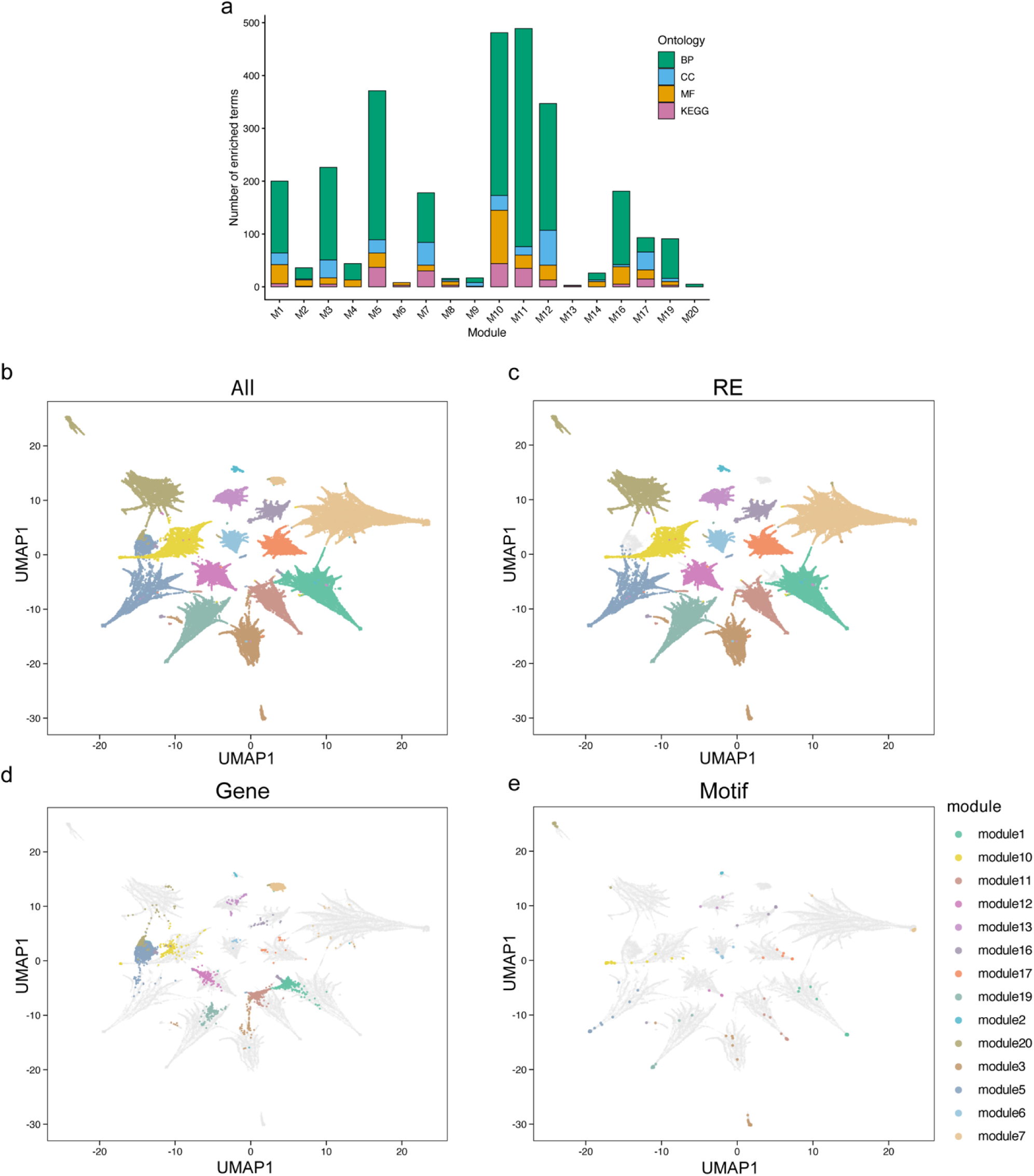
Functional enrichment and joint latent-space visualization of REGA-identified co-regulatory modules derived from bulk RNA-seq data. **(a)** Gene Ontology (GO) and KEGG enrichment analysis of REGA-derived modules. Stacked bar plots show the number of significantly enriched terms for each module across Biological Process (BP), Cellular Component (CC), Molecular Function (MF), and KEGG categories, highlighting the functional specificity of REGA-inferred modules. **(b–e)** Joint UMAP embedding of REGA-derived co-regulatory modules. UMAP visualizations show the shared latent space of regulatory elements (REs), target genes, and transcription factor (TF) motifs based on REGA-inferred module activities. Points represent individual entities colored by module assignment. **(b)** Integrated UMAP embedding of all entity types, showing that REs, genes, and motifs assigned to the same module co-cluster in the latent space. **(c–e)** UMAP embeddings highlighting **(c)** REs, **(d)** target genes, and **(e)** TF motifs, respectively. Highlighted entities are colored by module assignment, while all remaining entities are shown in grey for structural context.

**Supplementary Figure 5.**
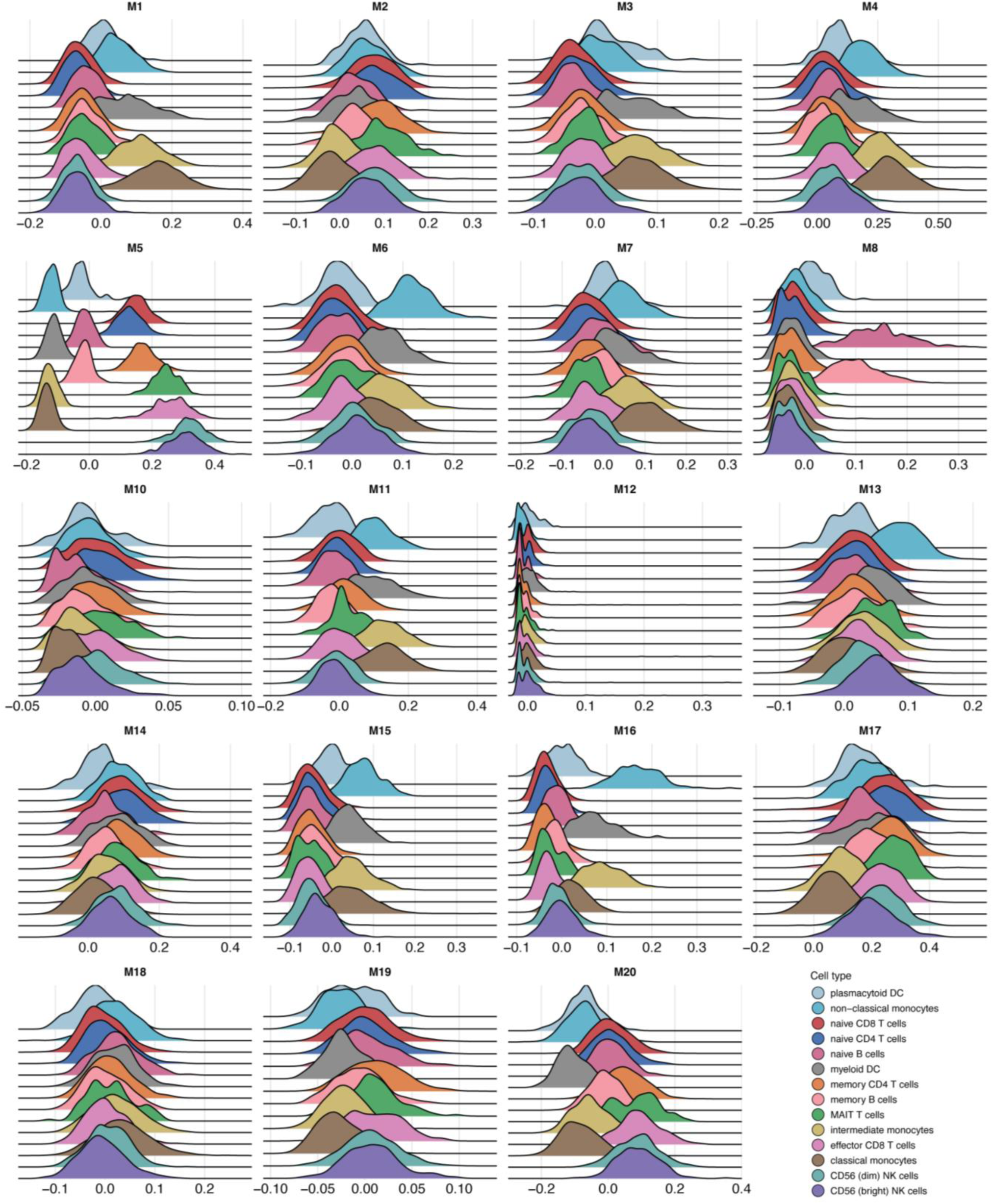
Ridge plots of REGA-derived module activities across blood cell types at single-cell resolution. Ridge plots show the distribution of REGA-derived module activity scores across blood cell types using a PBMC single-cell multiome dataset. Module activity was computed from log-normalized single-cell gene expression profiles using Seurat’s *AddModuleScore* function. Each panel corresponds to an individual module, and each curve represents the distribution of module activity within a specific cell type. The height of each ridge reflects the density of module scores across cells. Consistent with the pseudobulk analysis, REGA-derived modules exhibit distinct cell-type-specific activation patterns.

**Supplementary Figure 6.**
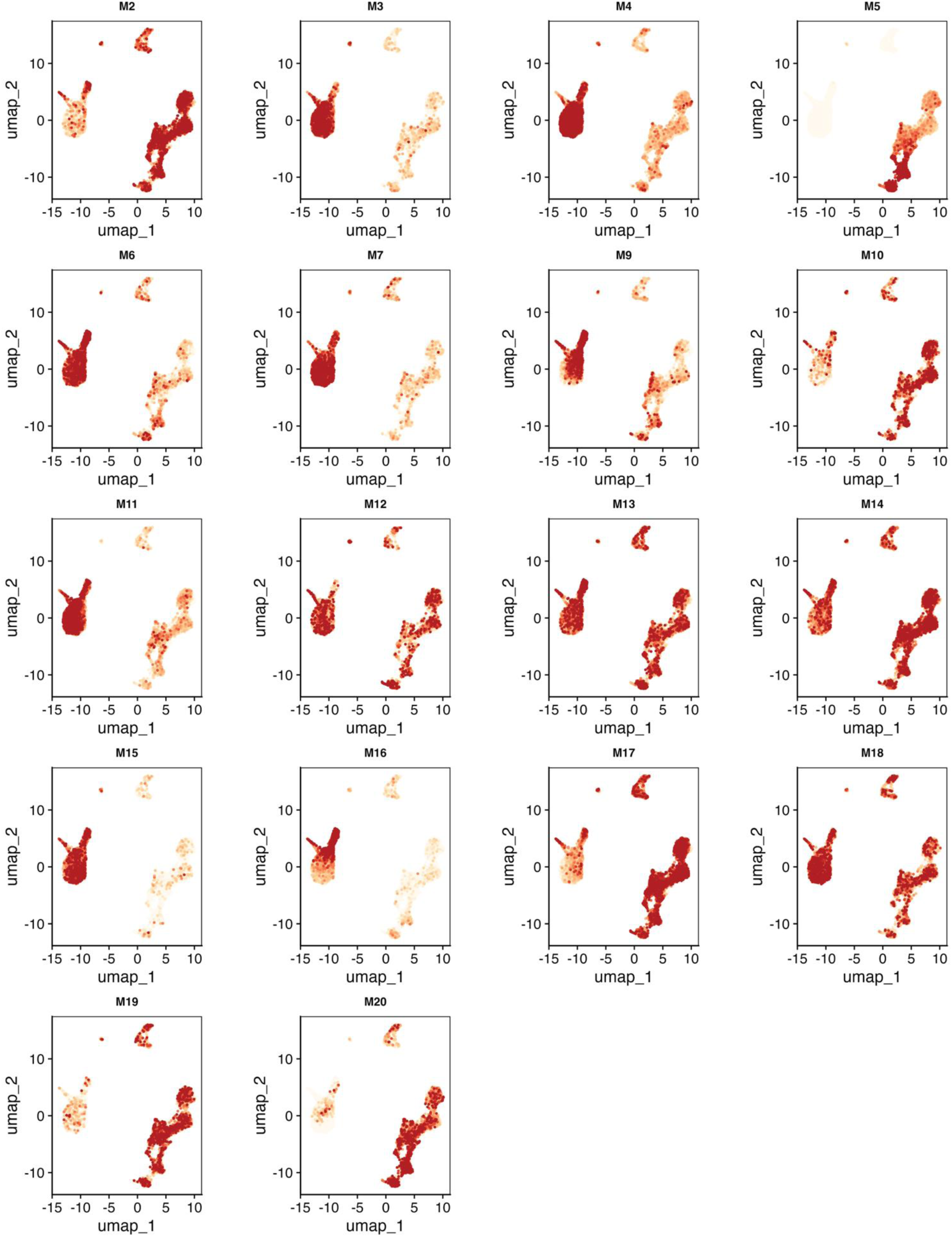
UMAP visualization of REGA-derived module activities at single-cell resolution. UMAP plots show the activity of REGA-derived modules across single cells in a PBMC single-cell multiome dataset. Each point represents a single cell and is colored by module activity. Module activity was calculated from log-normalized gene expression profiles using Seurat’s *AddModuleScore* function, with higher values indicating stronger module activation. Each panel corresponds to an individual module.

**Supplementary Figure 7.**
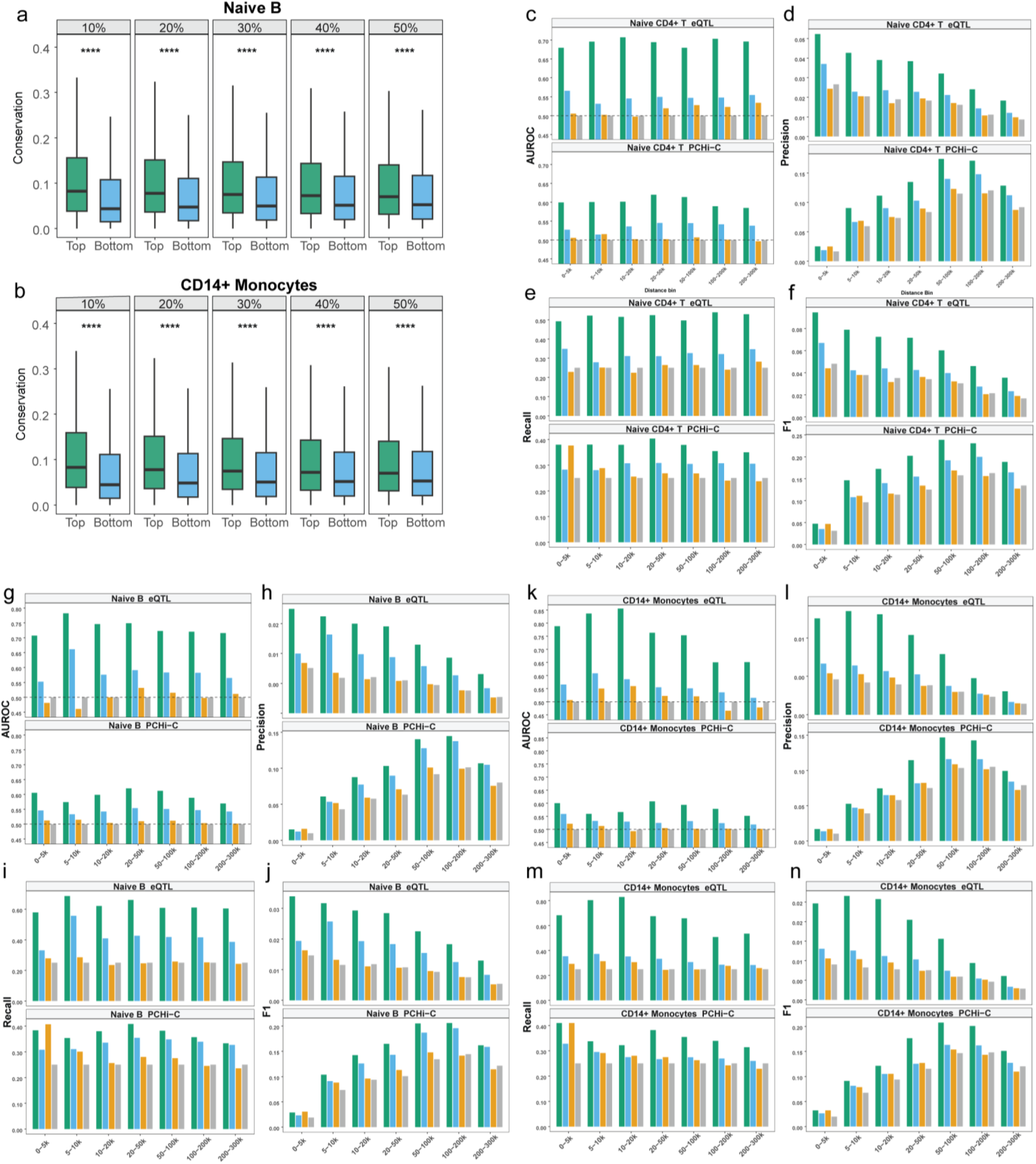
Performance assessment of cell type-specific cis-regulatory inference by REGA. (a–b) Evolutionary conservation of regulatory elements (REs) prioritized by REGA in naïve B cells and CD14+ monocytes. Boxplots compare mean sequence conservation scores between REs with high (“Top”) and low (“Bottom”) inferred regulatory importance across percentile thresholds (10–50%). Statistical significance was assessed using a one-sided *t*-test (**** *P* < 0.0001). **(c–n)** Performance evaluation of REGA for predicting regulatory element–target gene (RE–TG) interactions across genomic distance bins (0–5 kb to 200–300 kb relative to the transcription start site (TSS)) using single-cell RNA-seq data. Performance was evaluated using cell type-specific eQTL and promoter-capture Hi-C (pcHi-C) datasets as ground truth for naïve CD4+ T cells (**c–f**), naïve B cells (**g–j**), and CD14+ monocytes (**k–n**). REGA was benchmarked against three alternative methods: PCC (Pearson correlation between RE chromatin accessibility and target gene expression, computed using an independent 10x Genomics PBMC multiome dataset), Distance (a genomic distance-based decay function), and Random (a uniform baseline). Panels show area under the receiver operating characteristic curve (AUROC; **c, g, k**), precision (**d, h, l**), recall (**e, i, m**), and F1-score (**f, j, n**). Precision, recall, and F1-score were evaluated using the top quartile (25%) of predicted RE–TG interactions within each distance bin. The dashed line in the AUROC panels indicates the expected performance of a random classifier (0.5).

**Supplementary Figure 8.**
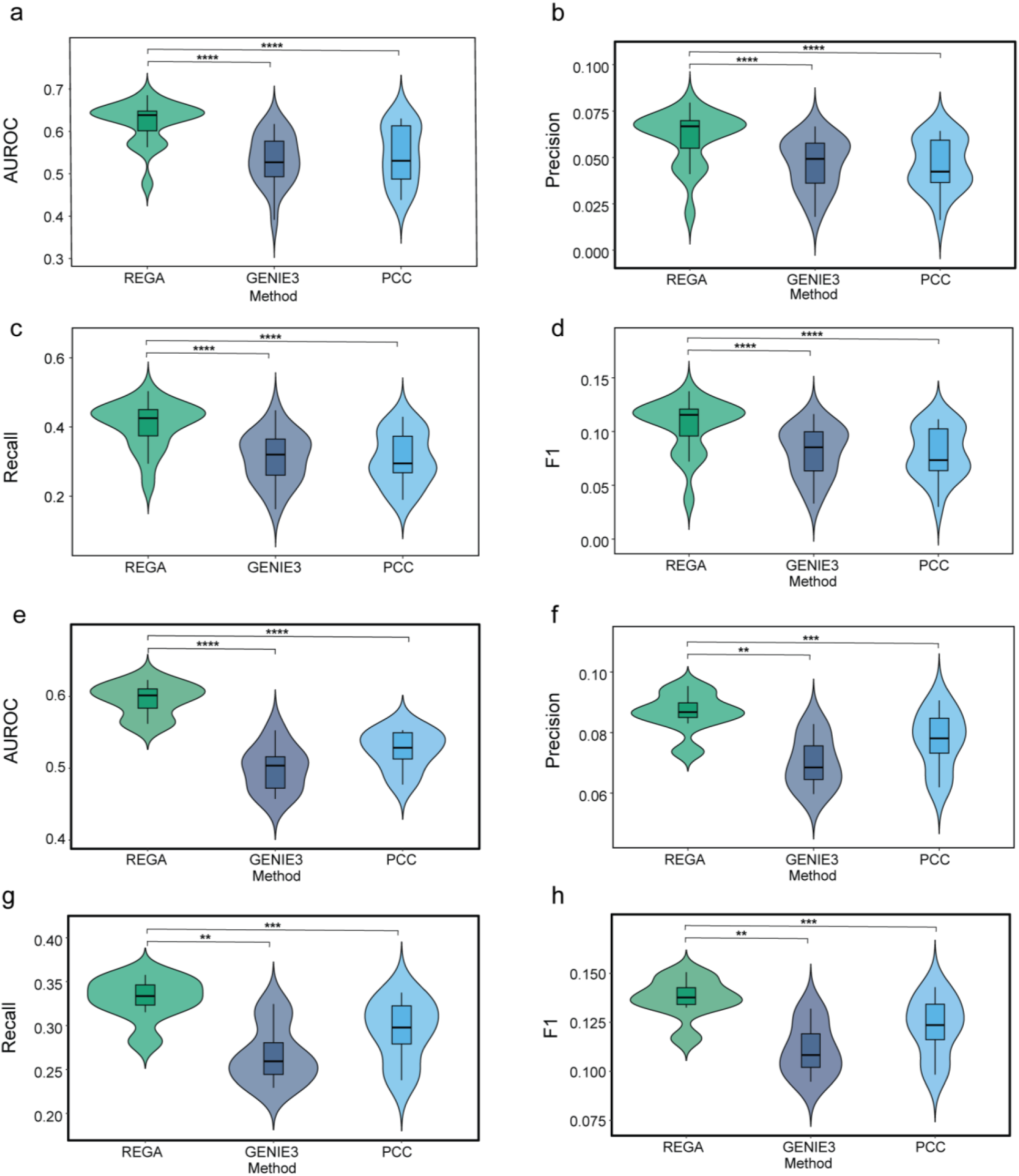
Performance assessment of cell type-specific trans-regulatory inference by REGA. (a–d) Evaluation of cell type-specific trans-regulatory target prediction using TF ChIP-seq data as ground truth. Violin plots show the performance of REGA, GENIE3, and PCC for predicting TF–target interactions across transcription factors (TFs) in CD4+ T cells, naïve B cells, and CD14+ monocytes. Ground-truth TF–target interactions were defined using experimental ChIP-seq data. Panels show **(a)** area under the receiver operating characteristic curve (AUROC), **(b)** precision, **(c)** recall, and **(d)** F1-score. Precision, recall, and F1-score were computed using the top quartile (25%) of predicted TF–target interactions. Statistical significance was assessed using paired one-sided Student’s *t*-tests (n = 19 TFs; **** *P* < 0.0001). **(e–h)** Validation of cell type-specific trans-regulatory inference using TF perturbation data from CD4+ T cells. Violin plots show the performance of REGA, GENIE3, and PCC for predicting direct differentially expressed genes (DEGs) following TF perturbation. Ground truth was defined using eight CD4+ T cell-specific perturbation datasets curated from the KnockTF database. Panels show **(e)** AUROC, **(f)** precision, **(g)** recall, and **(h)** F1-score. Precision, recall, and F1-score were computed using the top quartile (25%) of predicted TF–target interactions. Statistical significance was assessed using paired one-sided Student’s *t*-tests (** *P* < 0.01, *** *P* < 0.001).

**Supplementary Figure 9.**
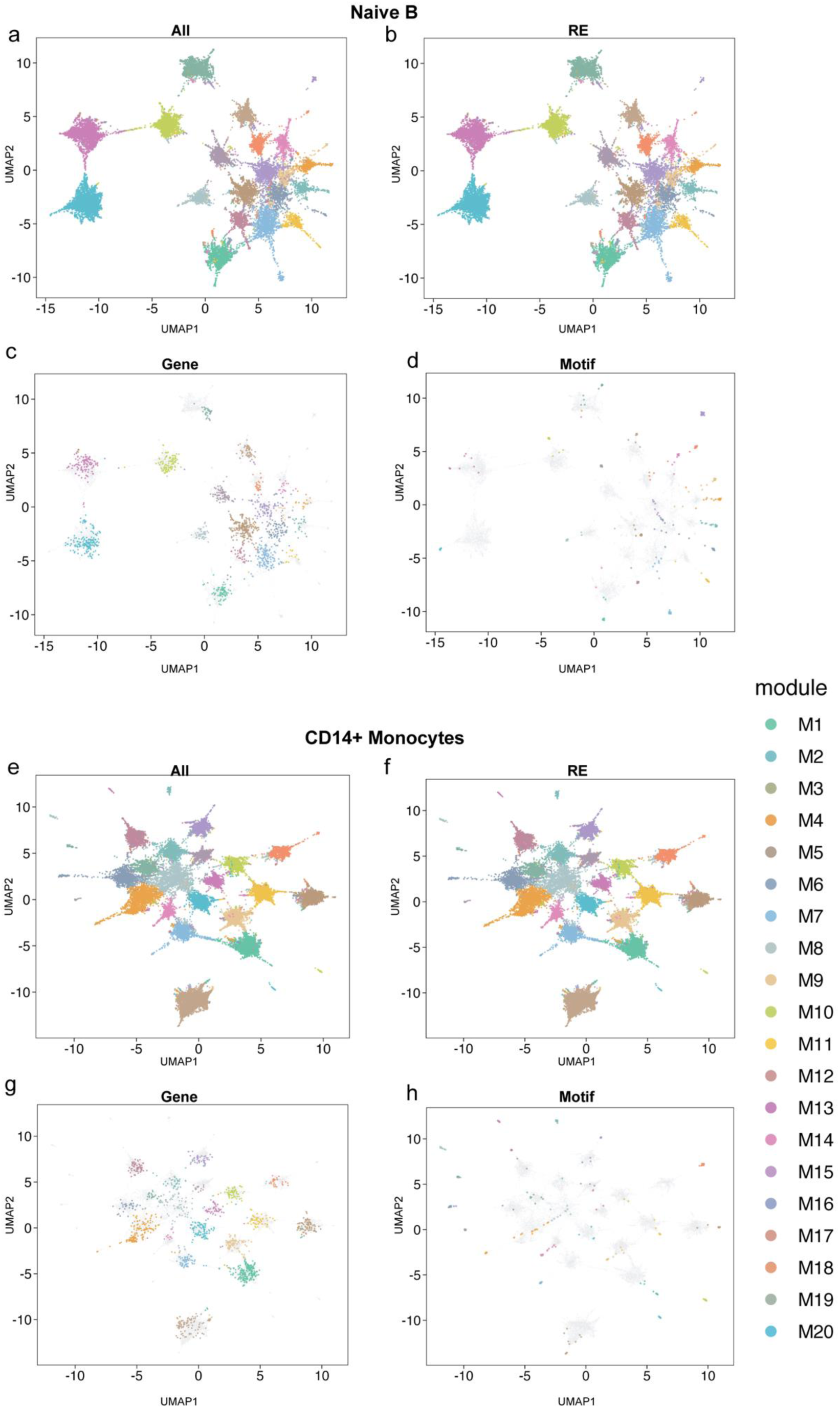
Joint latent-space visualization of REGA-derived co-regulatory modules in naïve B cells and CD14+ monocytes.(a–h) UMAP visualizations showing the joint latent space of regulatory elements (REs), target genes, and transcription factor (TF) motifs based on REGA-inferred module activities. Points represent individual entities colored by module assignment. (a–d) UMAP embeddings for naïve B cells. (a) Integrated UMAP embedding of all entity types (“All”), showing co-clustering of REs, genes, and motifs assigned to the same module. (b–d) UMAP embeddings highlighting (b) REs, (c) target genes, and (d) TF motifs, respectively. (e–h) UMAP embeddings for CD14+ monocytes. (e) Integrated UMAP embedding of all entity types (“All”). (f–h) UMAP embeddings highlighting (f) REs, (g) target genes, and (h) TF motifs, respectively. In panels highlighting specific entity types, selected entities are colored by module assignment, while all remaining entities are shown in light grey for structural context.

**Supplementary Figure 10.**
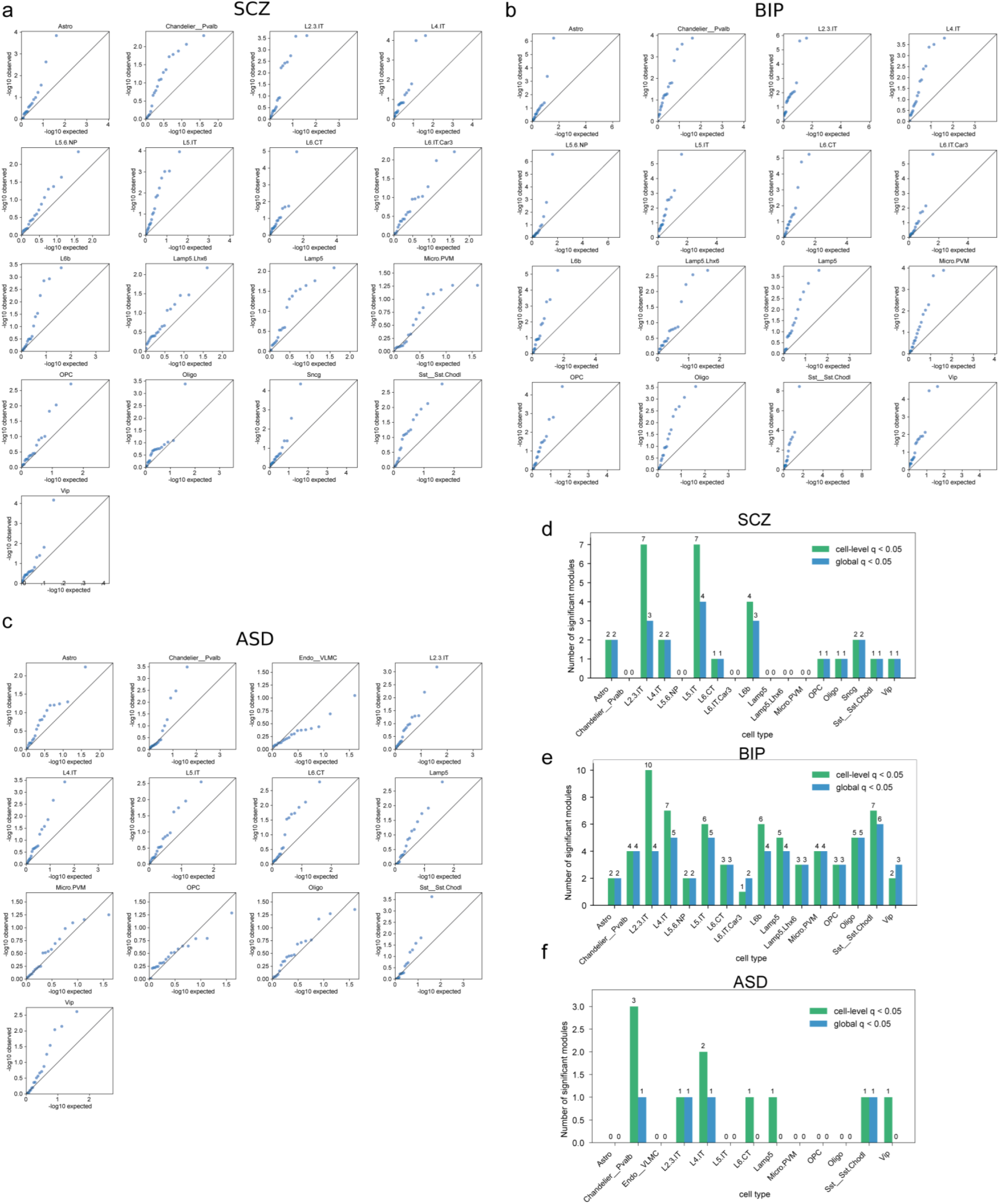
Cell type-specific disease-associated regulatory modules across neuropsychiatric disorders. (a–c) Q–Q plots showing associations between REGA-derived sample-level module activities and disease status across cell types for schizophrenia (SCZ; **a**), bipolar disorder (BIP; **b**), and autism spectrum disorder (ASD; **c**). Observed versus expected −log10(*P*) values from Spearman correlation analyses are shown for individual cell types. Deviation from the diagonal indicates enrichment of disease-associated modules within the corresponding cell type. **(d–f)** Number of significant disease-associated modules identified across cell types for SCZ (**d**), BIP (**e**), and ASD (**f**). Bars indicate the number of modules significant at the cell type-specific level (cell-level *q* < 0.05) or after global multiple-testing correction across all cell types and modules (global *q* < 0.05). These results demonstrate widespread disease-associated regulatory modules across diverse neural cell types.

**Supplementary Figure 11.**
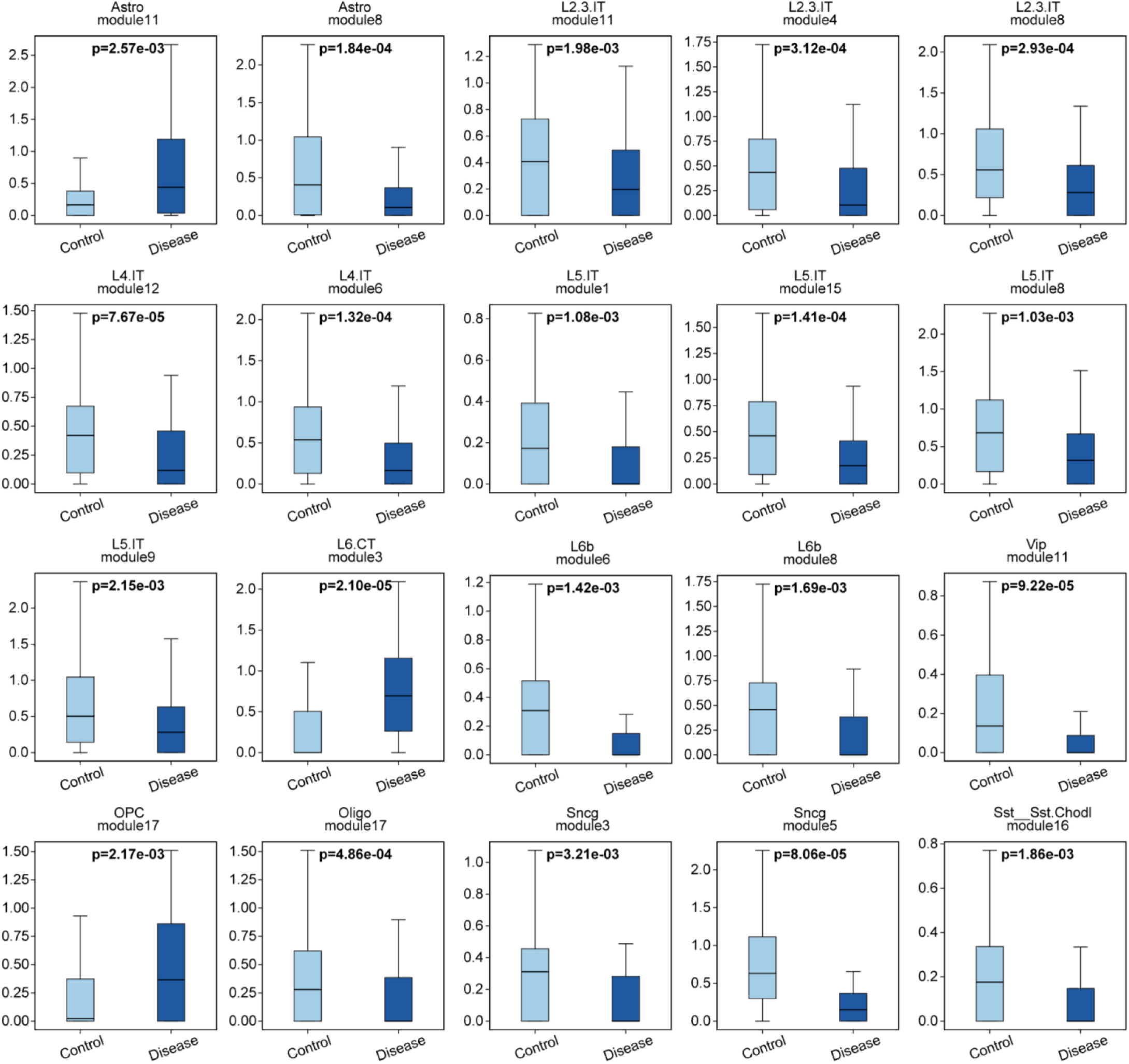
Differential activity of SCZ-associated regulatory modules between healthy control and SCZ samples. Boxplots show the distribution of REGA-inferred module activities between healthy control (light blue) and schizophrenia (SCZ) disease samples (dark blue) for representative SCZ-associated modules across cell types. The central line indicates the median, box bounds represent the interquartile range (IQR), and whiskers extend to the minimum and maximum values. Statistical significance was assessed using two-sided Wilcoxon rank-sum tests. Corresponding *P*- values are shown above each panel.

**Supplementary Figure 12.**
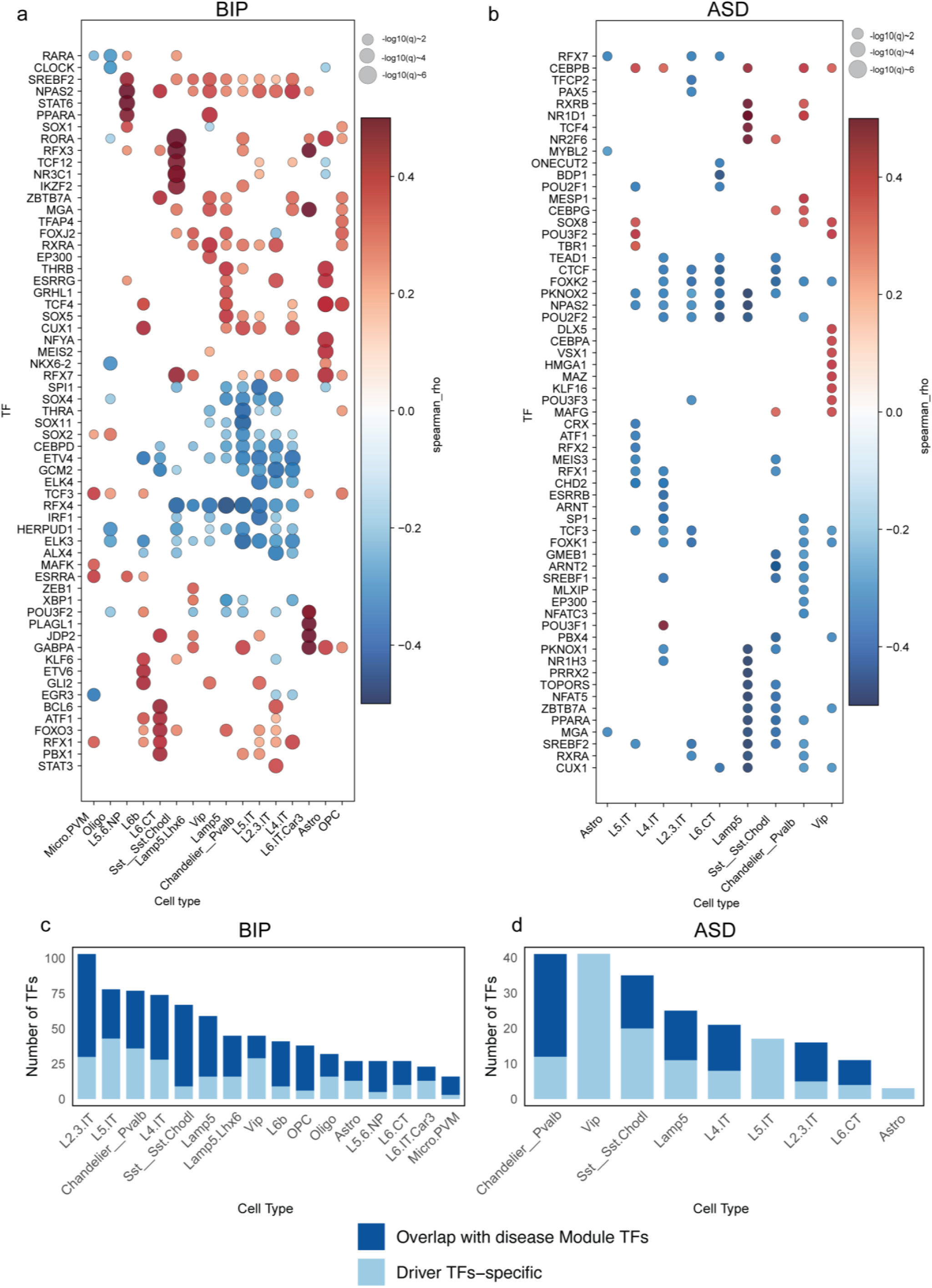
Cell type-specific driver transcription factors in bipolar disorder (BIP) and autism spectrum disorder (ASD). (a–b) Representative cell type-specific driver transcription factors (TFs) in BIP (**a**) and ASD (**b**). Dot plots show the association between REGA-inferred TF regulatory activity and disease status across cell types. The y-axis lists representative driver TFs, and the x-axis indicates the corresponding cell types. Dot color represents the Spearman correlation coefficient (ρ) between inferred TF activity and disease status, with red and blue indicating positive and negative associations, respectively. Dot size reflects statistical significance (−log10(*q*)). Driver TFs were identified using global false discovery rate (FDR) correction across all tested TFs and cell types, with significance defined as FDR < 0.1. **(c–d)** Overlap between cell type-specific driver TFs and disease-associated module TFs in BIP (**c**) and ASD (**d**). Stacked bar plots show the total number of identified driver TFs across cell types. Bars are partitioned into TFs overlapping with disease-associated module TFs (dark blue) and driver TF-specific TFs without module overlap (light blue). Cell types are ordered by the total number of identified driver TFs.

**Supplementary Figure 13.**
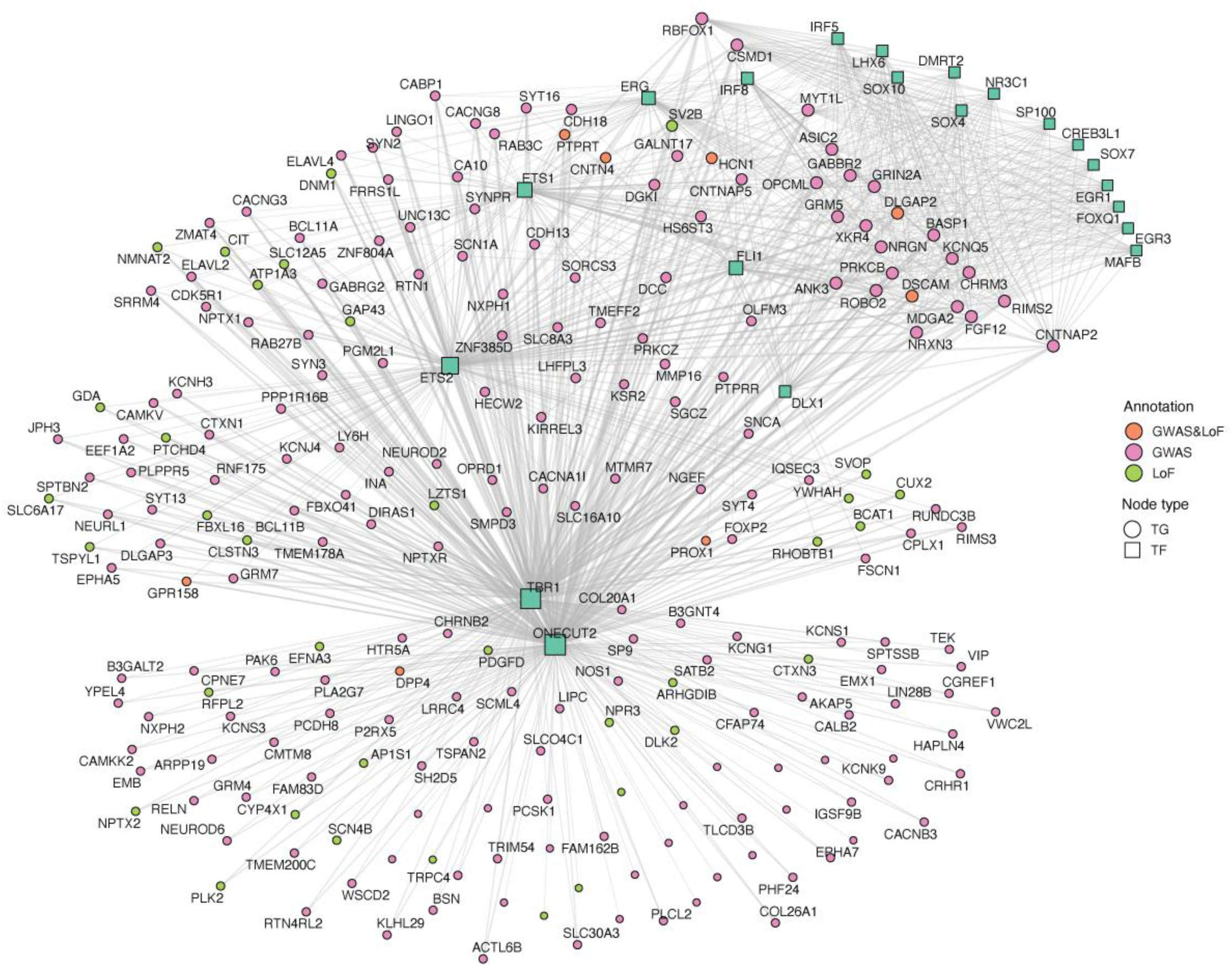
TF–TG regulatory network of the SCZ-associated Astro module 8 integrating genetic evidence. Network visualization of transcription factor (TF)–target gene (TG) interactions within the SCZ-associated Astro module 8 after restricting displayed target genes to those supported by independent genetic evidence. TFs are represented as square nodes, and target genes are represented as circular nodes. Edges denote REGA-inferred regulatory interactions between TFs and their downstream targets. Target genes are color-coded according to the type of supporting genetic evidence to highlight the disease relevance of the module. TGs with significant common variant associations identified through MAGMA gene-level analysis of schizophrenia (SCZ) GWAS data (FDR < 0.05) are shown in purple (GWAS). TGs harboring significant rare loss-of-function (LoF) variant associations identified through gene burden testing (FDR < 0.05) are shown in green (LoF). TGs supported by both GWAS and LoF evidence are highlighted in orange (GWAS & LoF).

**Supplementary Figure 14.**
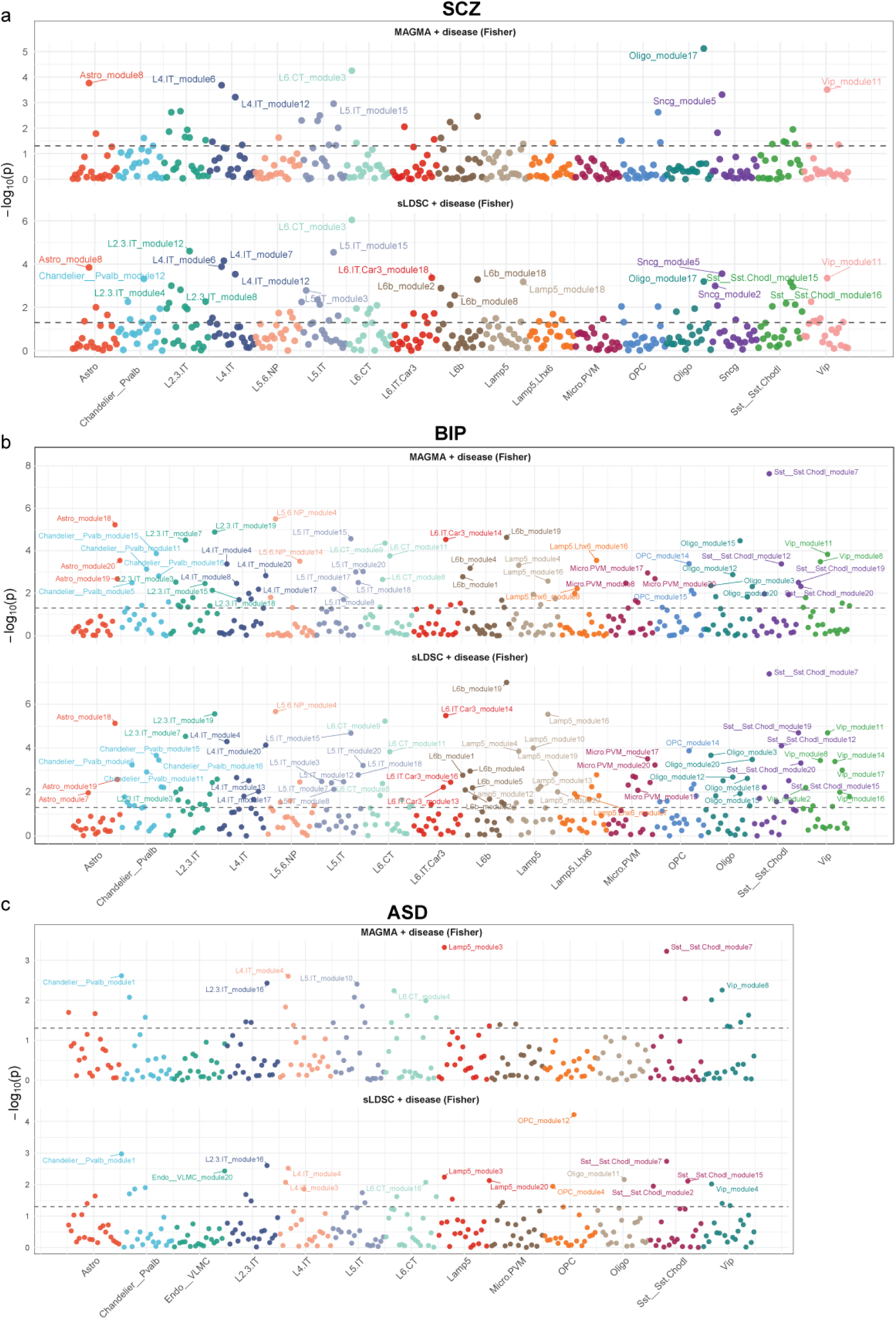
Manhattan-style scatter plots highlighting highly significant disease-associated modules supported by genetic evidence. (a–c) Manhattan-style multi-track plots showing integrated disease and genetic association signals for individual regulatory modules in schizophrenia (SCZ; **a**), bipolar disorder (BIP; **b**), and autism spectrum disorder (ASD; **c**). Modules are grouped and colored by cell type along the x-axis, with module order preserved within each cell type. For each disease, the upper track (“MAGMA + disease (Fisher)”) displays significance scores obtained by integrating disease-association *P*-values from REGA-derived sample-level module activities with MAGMA gene-level GWAS enrichment signals using Fisher’s combined probability test. The lower track (“sLDSC + disease (Fisher)”) shows integrated significance scores combining disease-association *P*-values with stratified LD score regression (sLDSC) enrichment of module-specific regulatory elements. The y-axis represents −log10(*P*) values, and dashed horizontal lines indicate nominal significance thresholds (*P* = 0.05). Modules passing global false discovery rate correction (Benjamini–Hochberg-adjusted *q* < 0.05) for either integrated analysis are labeled. These results highlight disease-associated regulatory modules jointly supported by REGA-derived regulatory dysregulation and independent genetic evidence.

**Supplementary Figure 15.**
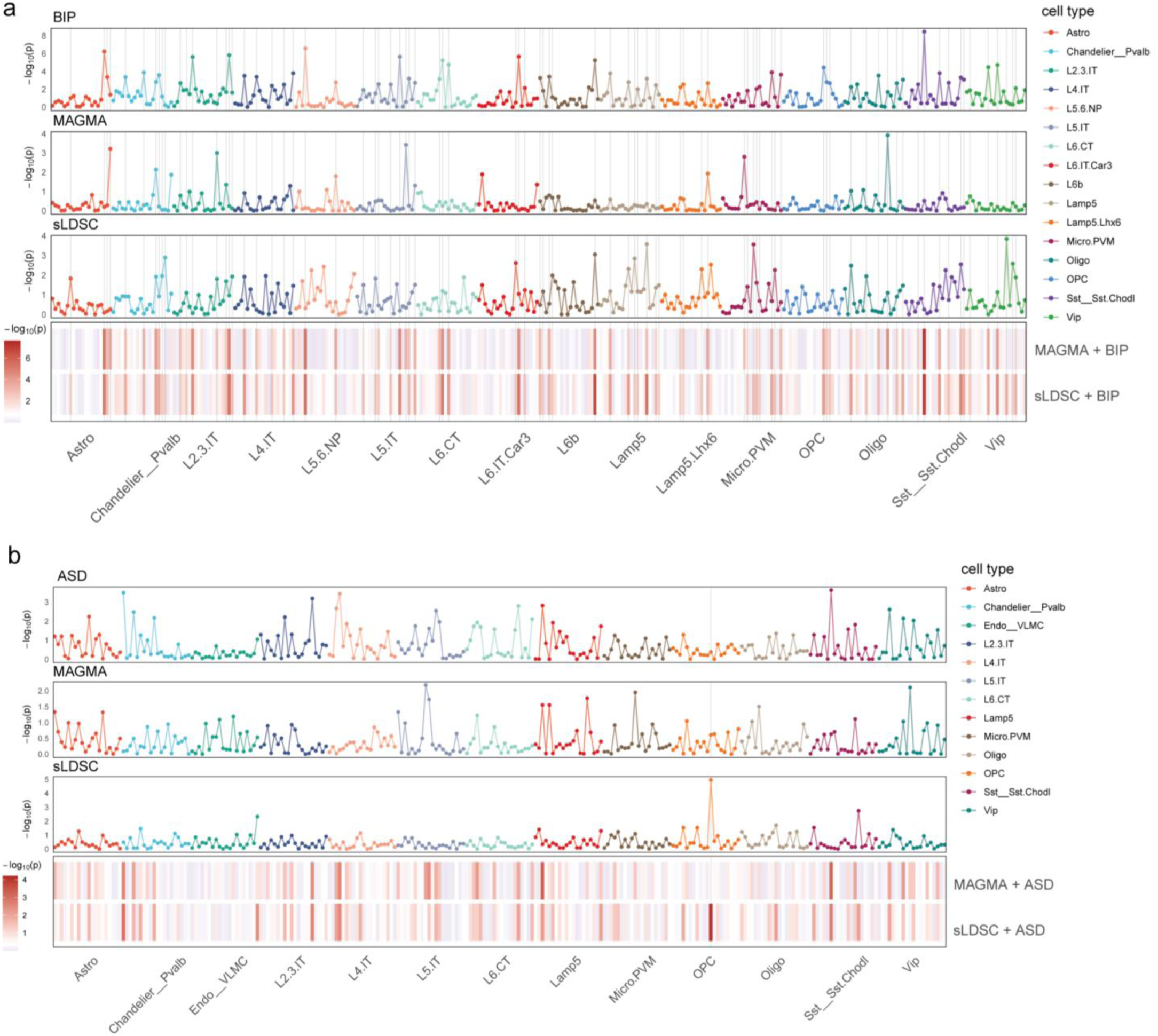
Integration of module–disease associations with genetic evidence across cell types for bipolar disorder (BIP) and autism spectrum disorder (ASD). (a–b) Multi-track plots showing the significance of disease associations for individual modules grouped and colored by cell type along the x-axis for BIP (**a**) and ASD (**b**). For each panel, the top three tracks display −log10(*P*) values across three independent layers of evidence: **(1)** BIP or ASD, representing associations between REGA-derived sample-level module activity and disease status; **(2)** MAGMA, showing gene-level GWAS enrichment of module target genes; and **(3)** sLDSC, showing heritability enrichment of module-specific regulatory elements assessed using stratified LD score regression. The bottom heatmaps show integrated significance scores obtained by combining disease-association *P*-values with MAGMA or sLDSC signals using Fisher’s combined probability test. Color intensity represents combined −log10(*P*) values, highlighting disease-associated modules supported by both REGA-inferred regulatory dysregulation and genetic risk variants.

**Supplementary Figure 16.**
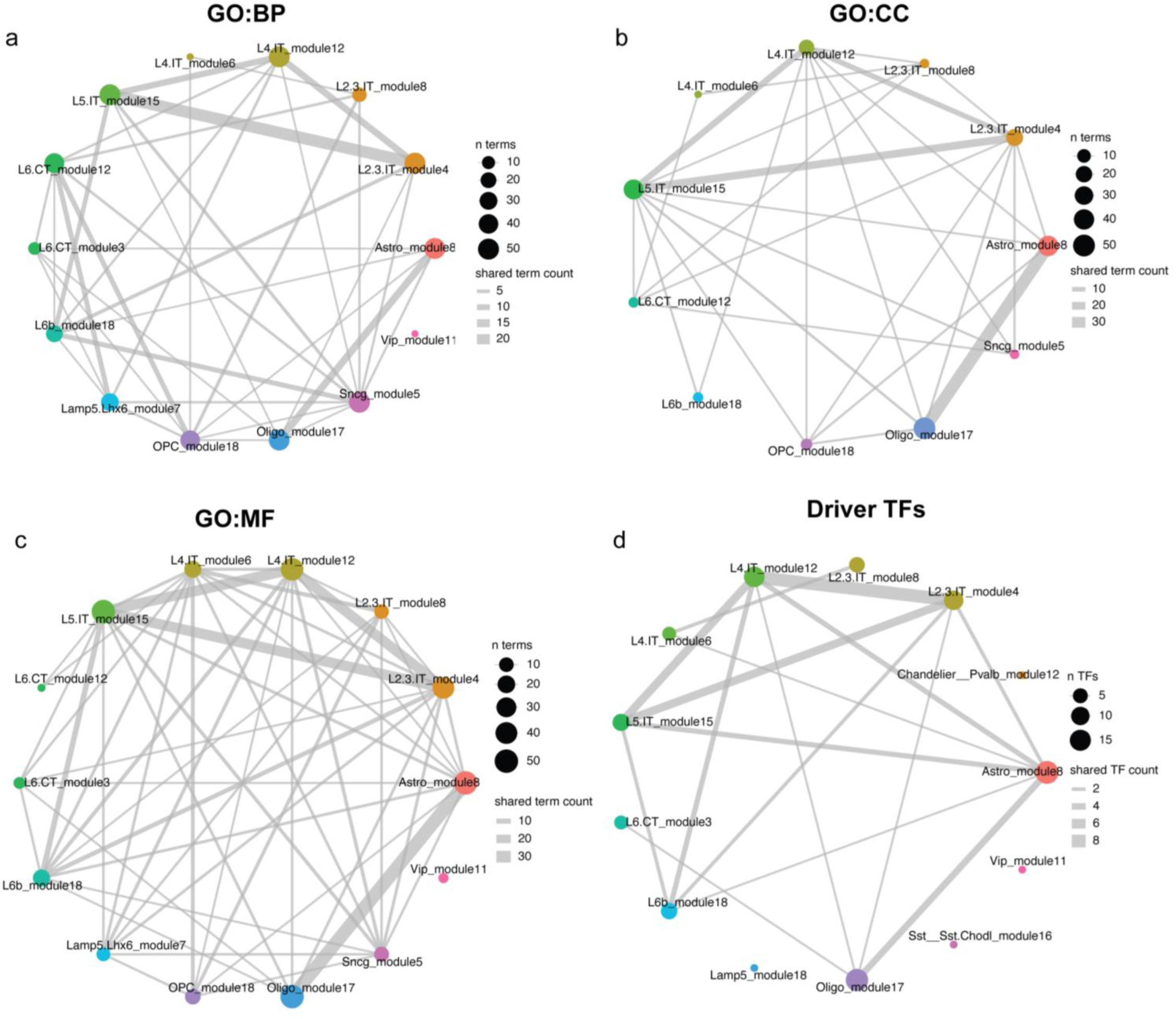
Shared functional terms and driver TF networks among genetically supported SCZ modules. (a–c) Network plots showing pairwise overlap among genetically supported SCZ-associated modules based on shared Gene Ontology (GO) terms identified by Gene Set Enrichment Analysis (GSEA): **(a)** GO:BP, **(b)** GO:CC, and **(c)** GO:MF. Nodes represent modules, with node size proportional to the number of enriched terms. Edges connect modules sharing enriched terms, and edge width reflects the number of shared terms. **(d)** Network plot showing pairwise overlap among the same modules based on shared driver transcription factors (TFs). Node size is proportional to the number of driver TFs, and edge width reflects the number of shared TFs. Modules are colored by cell type. Together, these networks highlight shared functional programs and regulatory architecture across genetically supported disease-associated modules.

**Supplementary Figure 17.**
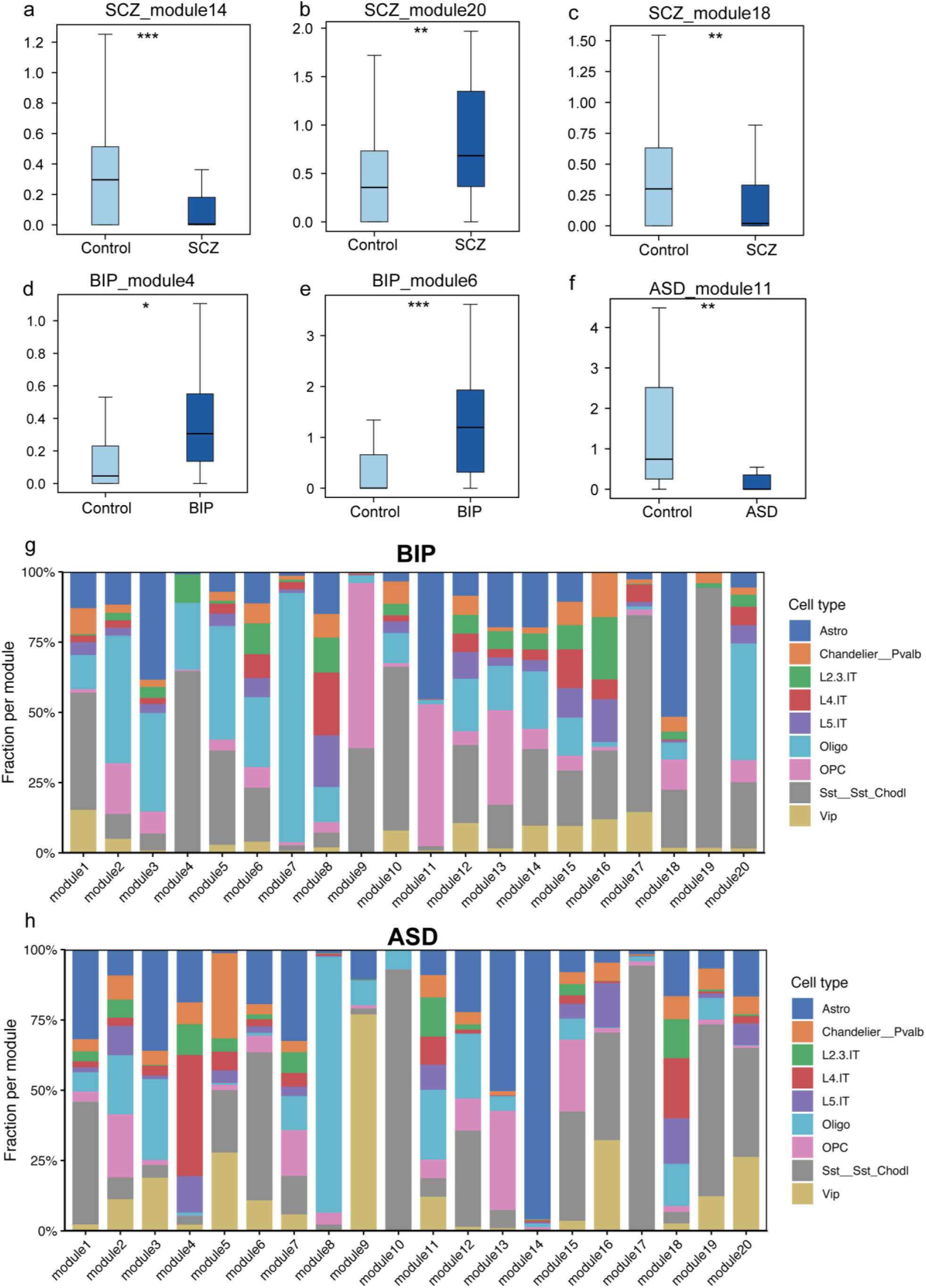
Differential activity and cell-type composition of cross-cell-type shared disease modules. (a–f) Boxplots showing the distribution of REGA-inferred activity for representative cross-cell-type shared modules between healthy control and disease samples. Panels correspond to SCZ modules 14, 20, and 18 (**a–c**), BIP modules 4 and 6 (**d–e**), and ASD module 11 (**f**). The central line indicates the median, box bounds represent the interquartile range (IQR), and whiskers extend to the minimum and maximum values. Statistical significance was assessed using two-sided Wilcoxon rank-sum tests (* *P* < 0.05, ** *P* < 0.01, *** *P* < 0.001). **(g–h)** Cell-type composition of cross-cell-type shared modules in BIP (**g**) and ASD (**h**). Stacked bar plots show the fraction of genes assigned to each module that originate from different cell types. Each bar represents a module, and colors denote distinct cell types, illustrating the cellular contributions to individual cross-cell-type shared regulatory modules.

**Supplementary Figure 18.**
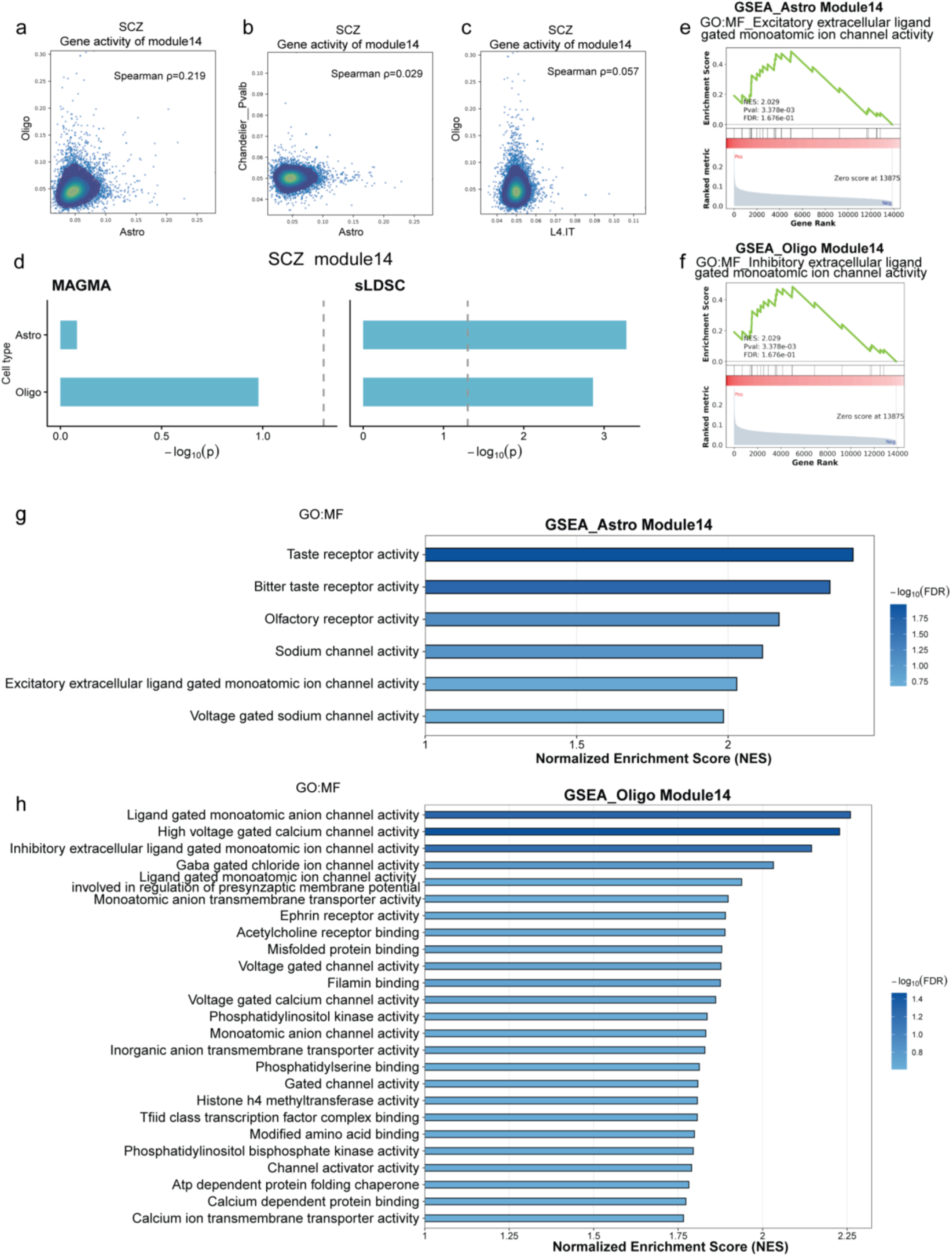
Cross-cell-type genetic and functional characterization of SCZ module 14. (a–c) Cross-cell-type comparison of gene loadings in SCZ module 14. Pairwise scatter plots show normalized gene loadings of shared genes between different cell types. Each point represents a gene, with axes indicating normalized gene loadings in the corresponding cell types. Spearman correlation coefficients (ρ) are shown in each panel to quantify concordance across cell types. Color gradients and contour lines represent local point density estimated by two-dimensional kernel density estimation, with warmer colors indicating higher gene density. Panels correspond to the following comparisons: **(a)** Oligo versus Astro, **(b)** Chandelier_Pvalb versus Astro, and **(c)** Oligo versus L4.IT. **(d)** Cell type-specific genetic enrichment of SCZ module 14. Bar plots show the significance of genetic associations across cell types based on MAGMA (left) and stratified LD score regression (sLDSC; right). The x-axis represents −log10(*P*) values, with higher values indicating stronger enrichment. Dashed vertical lines indicate the significance threshold (−log10(*P*) = 1.3, corresponding to *P* = 0.05). **(e–f)** Representative Gene Set Enrichment Analysis (GSEA) enrichment plots for SCZ module 14. Panels show enrichment of **(e)** excitatory extracellular ligand-gated monoatomic ion channel activity and **(f)** inhibitory extracellular ligand-gated monoatomic ion channel activity in Astro and Oligo module 14, respectively. **(g–h)** Cell type-specific functional enrichment of SCZ module 14. Bar plots show representative Gene Ontology Molecular Function (GO:MF) terms identified by GSEA in **(g)** Astro and **(h)** Oligo cell types. The x-axis represents normalized enrichment scores (NES), and color intensity indicates statistical significance (−log10 FDR).

**Supplementary Figure 19.**
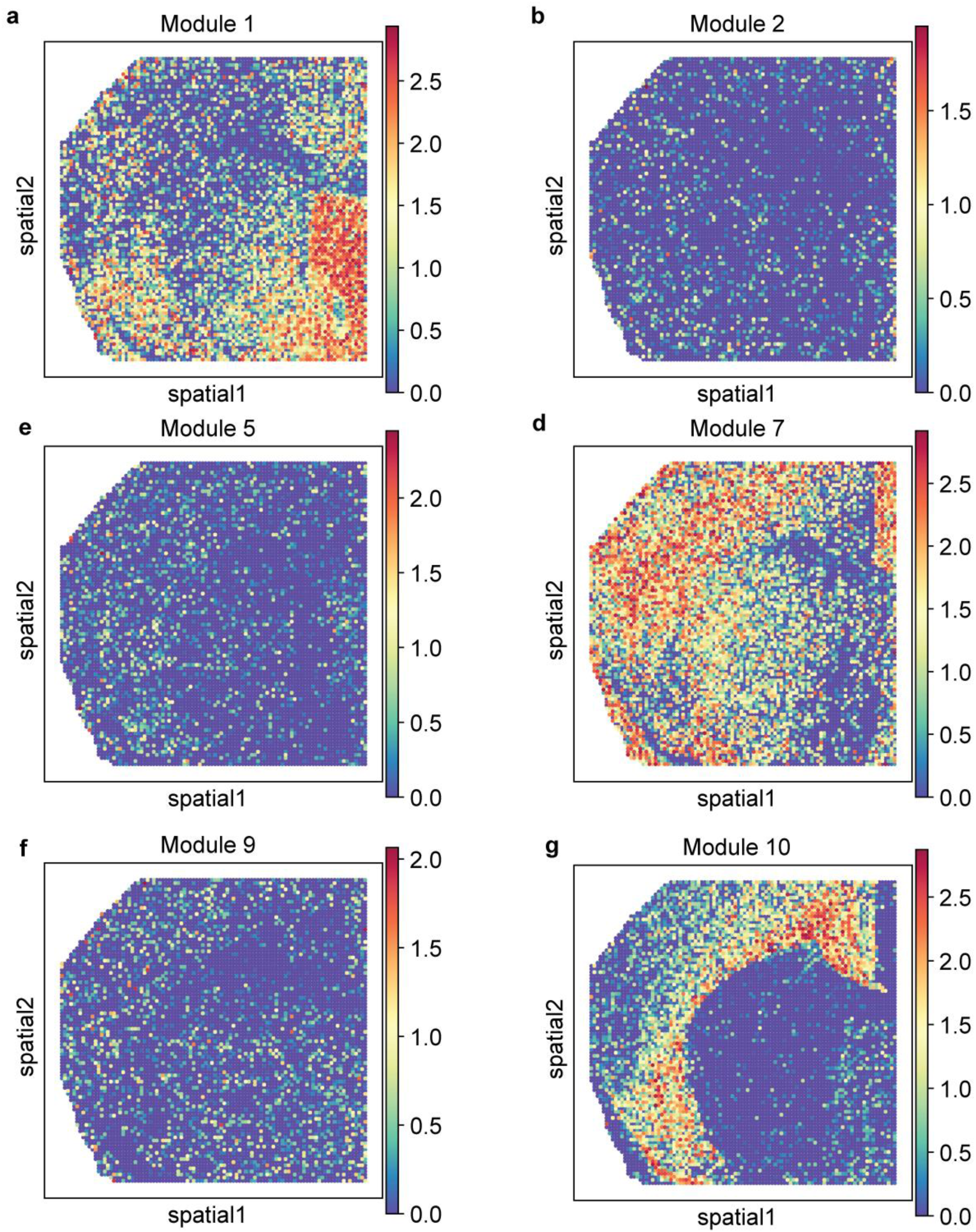
Spatial patterns of regulatory modules inferred from the REGA module activity matrix. (a–g) Spatial activity patterns of representative REGA-derived modules across the tissue section. Each panel shows the activity score of an individual module at each spatial spot, highlighting distinct spatial organization across tissue regions, including modules 1, 7, and 10.

**Supplementary Figure 20.**
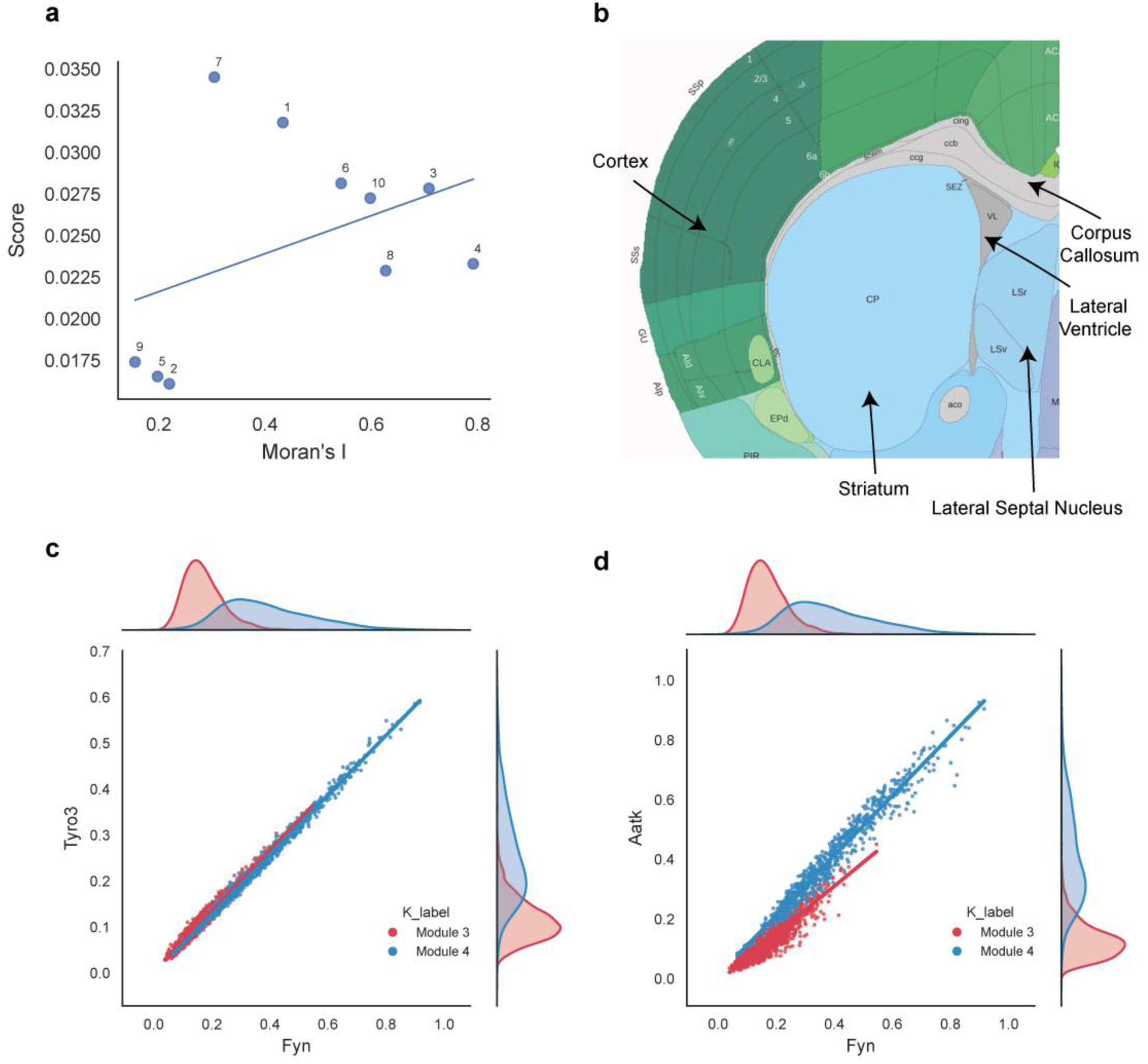
Spatial autocorrelation, anatomical context, and ligand–receptor expression patterns of REGA-derived modules. **(a)** Scatter plot showing the relationship between module Moran’s *I* and module contribution scores. Modules 2, 5, and 9 exhibit both low spatial autocorrelation and low contribution scores. **(b)** Anatomical regions from a coronal section of an adult mouse brain slice annotated using the Allen Brain Atlas. **(c–d)** Scatter plots showing the relationship between expression levels of indicated gene pairs in spots from Module 3 (red) and Module 4 (blue). Each point represents a single spatial spot. Solid lines indicate linear regression fits for each module, with shaded regions representing 95% confidence intervals. Marginal histograms along the x- and y-axes show the distribution of expression values for each gene within each module.

**Supplementary Figure 21.**
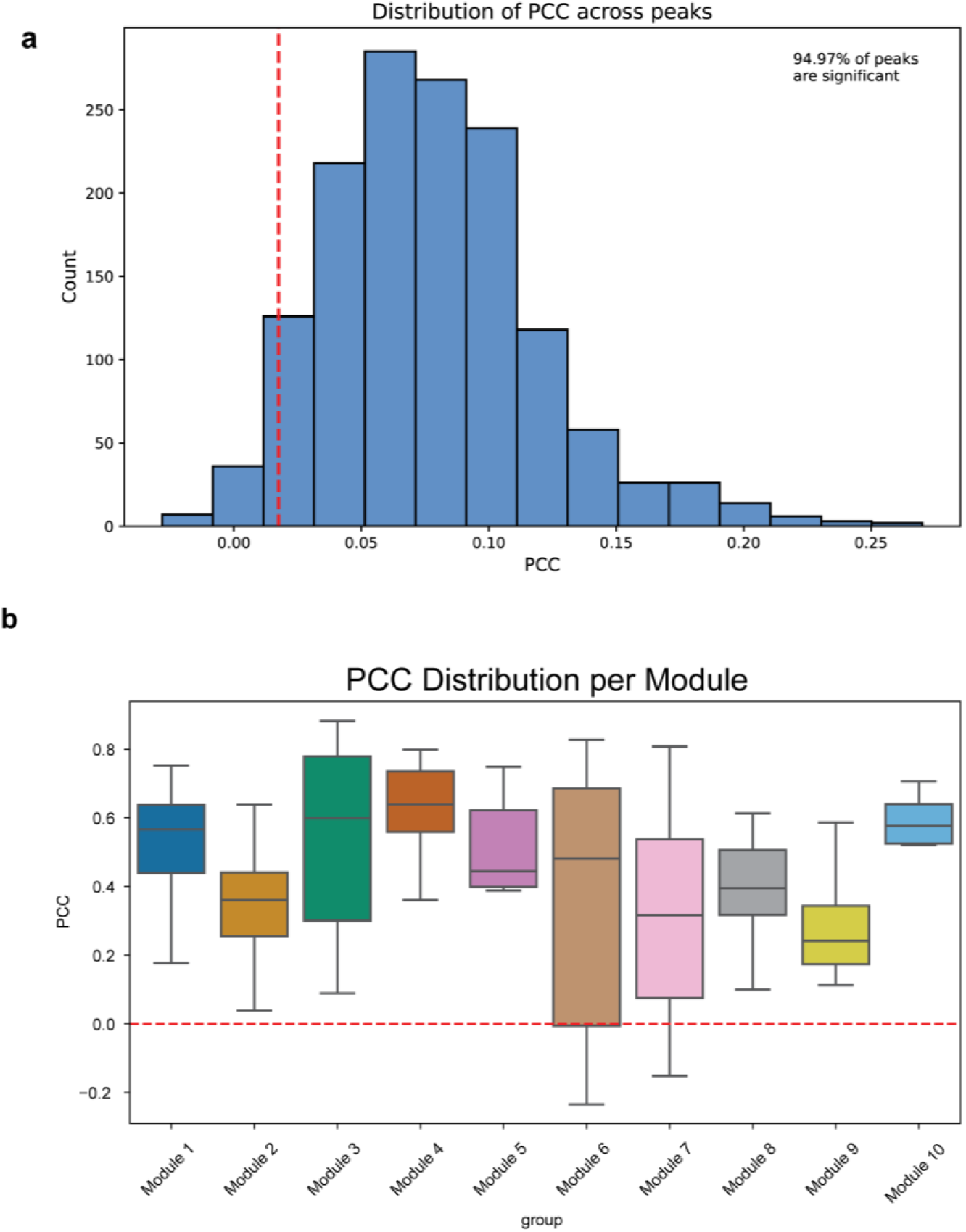
Evaluation of predicted spatial ATAC profiles and transcription factor activities inferred by REGA. **(a)** Histogram showing the distribution of Pearson correlation coefficients (PCCs) between predicted and observed ATAC values for selected top peaks (n = 1,432). The red vertical line indicates the FDR-based significance threshold at α = 0.05, defined as the minimum PCC among statistically significant peaks. **(b)** Boxplots showing the distribution of PCCs between chromVAR deviation scores computed from predicted spatial ATAC profiles and corresponding REGA-inferred transcription factor activities across modules. The red dashed line indicates PCC = 0.

**Supplementary Figure 22.**
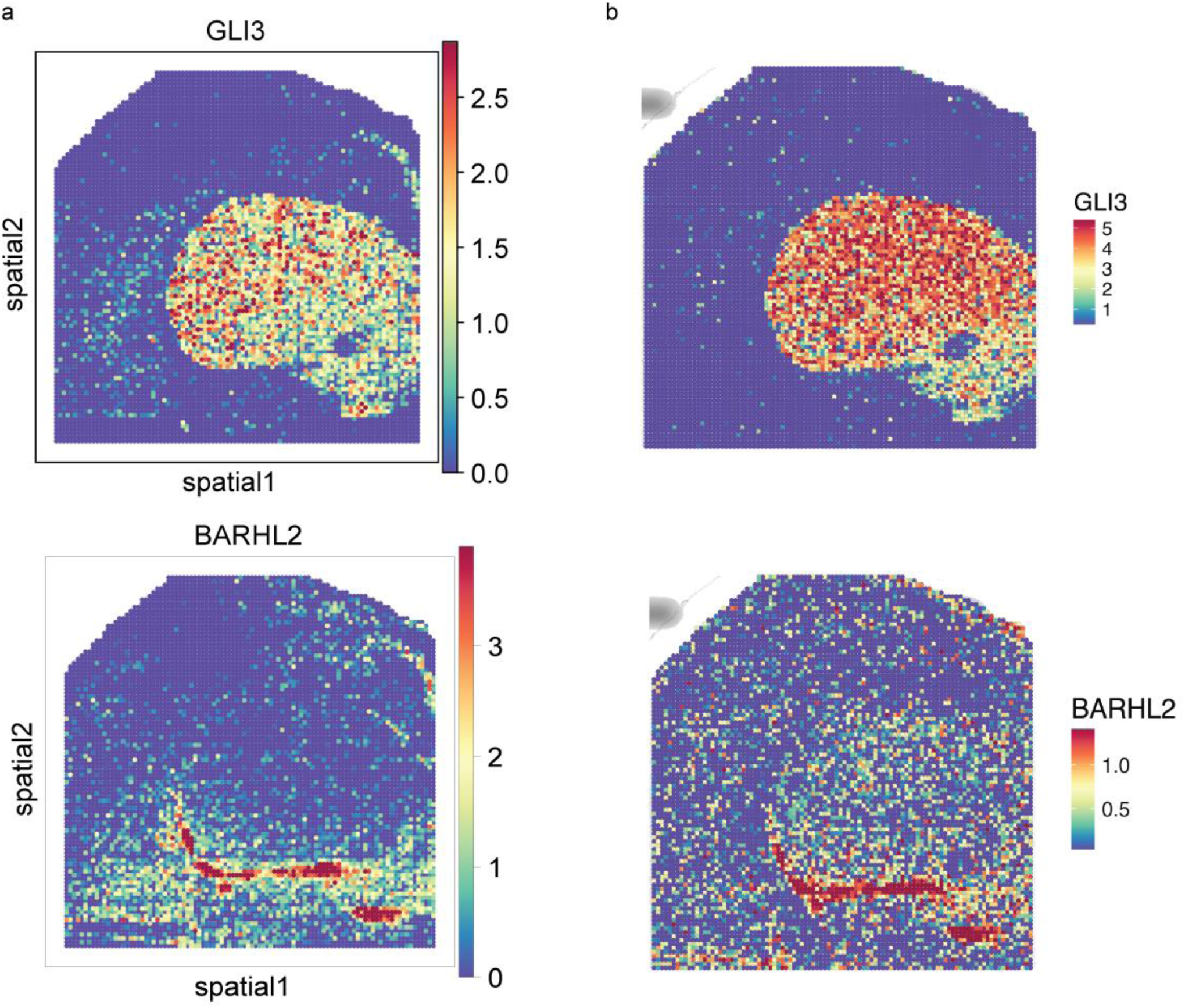
Agreement between spatial transcription factor activity patterns inferred by REGA and chromVAR. **(a)** Spatial activity patterns of selected transcription factors inferred by REGA. **(b)** Spatial activity patterns of the same transcription factors estimated by chromVAR using predicted spatial ATAC profiles. The inferred spatial patterns were highly consistent between the two approaches, as quantified by PCC values for GLI3 in module 3 (PCC = 0.88) and BARHL2 in module 8 (PCC = 0.61).

**Supplementary Figure 23.**
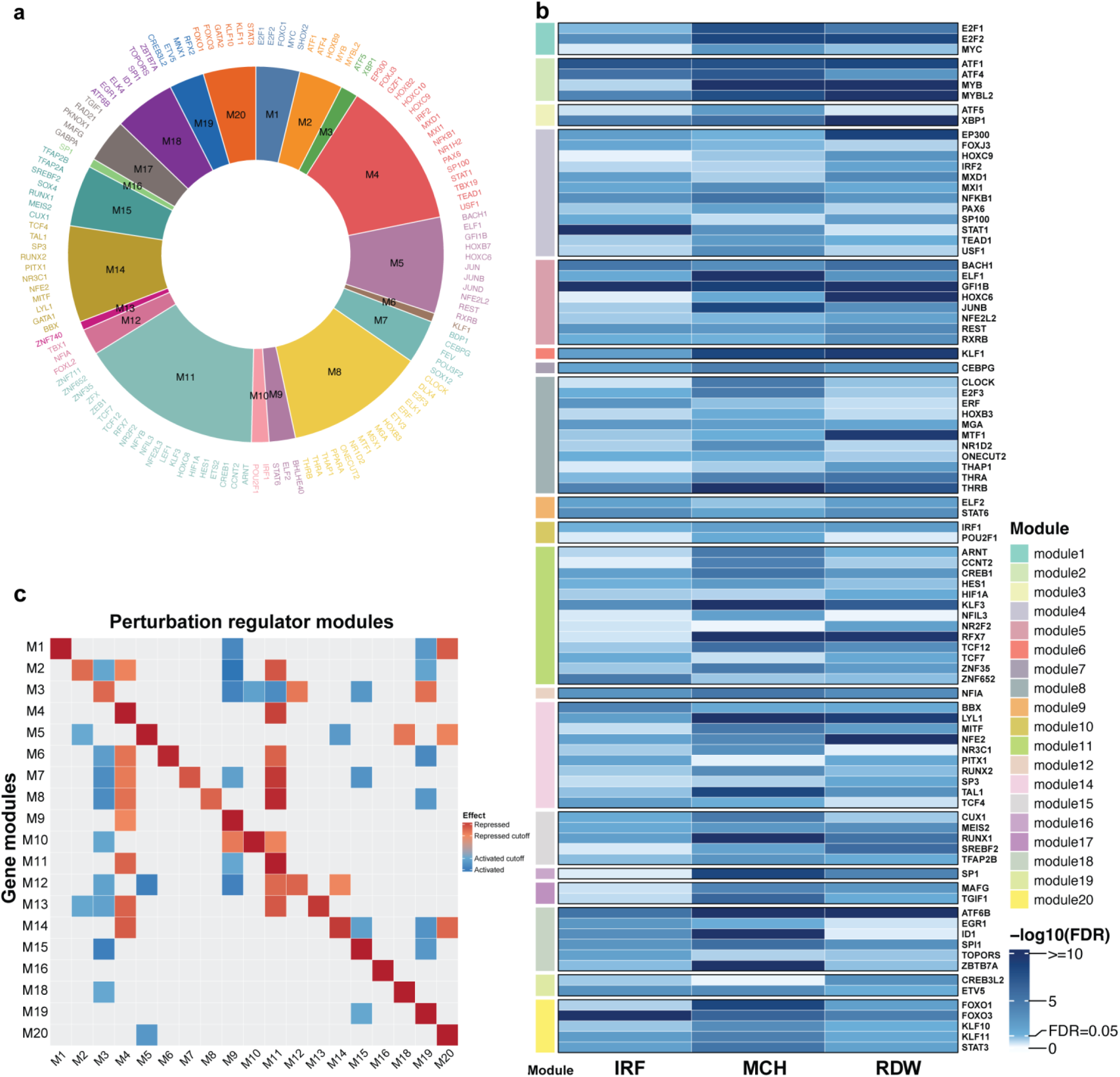
Regulatory modules derived from perturbation data. **(a)** Module assignment of transcription factors (TFs) across perturbation-derived regulator modules. Circular plot showing TFs assigned to individual REGA-derived modules (M1–M20), with colors indicating module identity. **(b)** Heatmap showing MAGMA gene-level GWAS significance of module-associated TFs across representative blood traits, including immature reticulocyte fraction (IRF), mean corpuscular hemoglobin (MCH), and red blood cell distribution width (RDW). Rows represent TFs, columns represent blood traits, and color intensity indicates statistical significance (−log10(FDR)). Only TFs significant in at least one trait are displayed. **(c)** Cross-module transcriptional responses between REGA-derived perturbation modules and gene modules. Gene expression profiles were aggregated according to REGA-inferred gene modules by averaging expression of genes assigned to the same module. Perturbation targets were grouped according to their REGA module assignments and averaged to generate a perturbation-module-by-gene-module response matrix. The resulting matrix was normalized using row-wise z-score scaling to highlight relative transcriptional responses across perturbation modules. Rows correspond to REGA-derived gene modules and columns correspond to REGA-derived perturbation target modules. Red indicates increased expression of a gene module following perturbation, implying that the corresponding perturbation target module normally represses that gene module. Blue indicates decreased expression following perturbation, implying that the corresponding perturbation target module normally activates that gene module. Only strong responses with absolute z-score ≥ 1 are displayed, while weaker responses are masked in grey. In the legend, “Repressed” and “Activated” denote the inferred regulatory effects of perturbation target modules on gene modules, whereas “Repressed cutoff” and “Activated cutoff” indicate the z-score thresholds (±1) used to define strong responses. Clustering was disabled to preserve the original module ordering.

**Supplementary Figure 24.**
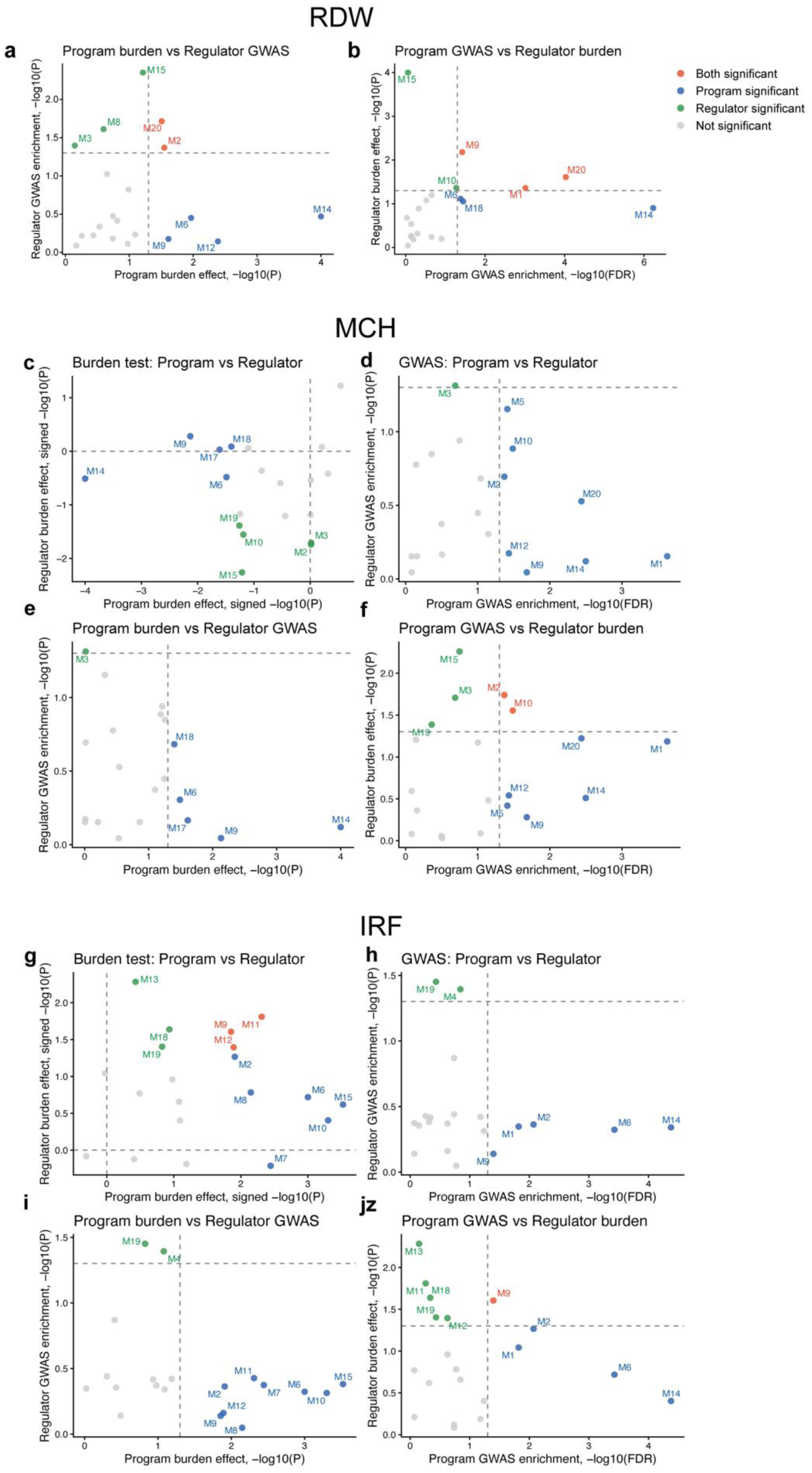
Genetic associations between module programs and regulators across blood traits. Scatter plots comparing genetic association signals between module target genes (“Program”, x-axis) and their corresponding regulators (“Regulator”, y-axis) across three representative blood traits: red blood cell distribution width (RDW; **a–b**), mean corpuscular hemoglobin (MCH; **c–f**), and immature reticulocyte fraction (IRF; **g–j**). Each point represents an individual module. Panels show **(a, c, g)** burden test effects for programs versus regulators, **(b, f, j)** program GWAS enrichment versus regulator burden effects, **(d, h)** GWAS enrichment for programs versus regulators, and **(e, i)** program burden effects versus regulator GWAS enrichment. For burden tests, values are represented as −log10(*P*) (or signed −log10(*P*) where indicated), with *P*-values derived from two-sided permutation tests (10,000 iterations). For GWAS analyses using MAGMA, program enrichment was evaluated using −log10(FDR), whereas regulator enrichment was evaluated using unadjusted −log10(*P*) values because of the limited number of regulator genes. Dashed lines indicate significance thresholds. Modules are color-coded according to joint significance status: both program and regulator significant (orange), only program significant (blue), only regulator significant (green), and not significant (grey).

